# p300/CBP degradation is required to disable the active AR enhanceosome in prostate cancer

**DOI:** 10.1101/2024.03.29.587346

**Authors:** Jie Luo, Zhixiang Chen, Yuanyuan Qiao, Jean Ching-Yi Tien, Eleanor Young, Rahul Mannan, Somnath Mahapatra, Tongchen He, Sanjana Eyunni, Yuping Zhang, Yang Zheng, Fengyun Su, Xuhong Cao, Rui Wang, Yunhui Cheng, Rithvik Seri, James George, Miriam Shahine, Stephanie J. Miner, Ulka Vaishampayan, Mi Wang, Shaomeng Wang, Abhijit Parolia, Arul M. Chinnaiyan

## Abstract

Prostate cancer is an exemplar of an enhancer-binding transcription factor-driven disease. The androgen receptor (AR) enhanceosome complex comprised of chromatin and epigenetic coregulators assembles at enhancer elements to drive disease progression. The paralog lysine acetyltransferases p300 and CBP deposit histone marks that are associated with enhancer activation. Here, we demonstrate that p300/CBP are determinant cofactors of the active AR enhanceosome in prostate cancer. Histone H2B N-terminus multisite lysine acetylation (H2BNTac), which is exclusively reliant on p300/CBP catalytic function, marked active enhancers and was notably elevated in prostate cancer lesions relative to the adjacent benign epithelia. Degradation of p300/CBP rapidly depleted acetylation marks associated with the active AR enhanceosome, which was only partially phenocopied by inhibition of their reader bromodomains. Notably, H2BNTac was effectively abrogated only upon p300/CBP degradation, which led to a stronger suppression of p300/CBP-dependent oncogenic gene programs relative to bromodomain inhibition or the inhibition of its catalytic domain. *In vivo* experiments using an orally active p300/CBP proteolysis targeting chimera (PROTAC) degrader (CBPD-409) showed that p300/CBP degradation potently inhibited tumor growth in preclinical models of castration-resistant prostate cancer and synergized with AR antagonists. While mouse p300/CBP orthologs were effectively degraded in host tissues, prolonged treatment with the PROTAC degrader was well tolerated with no significant signs of toxicity. Taken together, our study highlights the pivotal role of p300/CBP in maintaining the active AR enhanceosome and demonstrates how target degradation may have functionally distinct effects relative to target inhibition, thus supporting the development of p300/CBP degraders for the treatment of advanced prostate cancer.

## Introduction

Cancer is characterized by a major reconfiguration of the epigenetic landscape, which aids in the proliferation and survival of tumor cells^1,2^. A key change noted in many forms of cancer, including metastatic castration-resistant prostate cancer (mCRPC), is an escalated dependence on an aberrant enhancer landscape^3,4^. In mCRPC, crucial oncogenic transcription factors, including the androgen receptor (AR), FOXA1, HOXB13, ERG, and MYC, recruit and collaborate with epigenetic coregulators at enhancer sites in cancer-specific complexes known as *neo*-enhanceosomes to create active chromatin environments that drive hyper-expression of oncogenic genes through looping interactions with promoters^4-9^. Importantly, hyper-expression of the *AR*, *FOXA1*, and *MYC* genes themselves are wired through looping interactions with multi-enhancer clusters^4,7-9^. Earlier research demonstrated that when AR-positive prostate cancer cells were treated with a proteolysis-targeting chimera (PROTAC) that degraded the ATPase subunits of the SWI/SNF chromatin remodeling complex (namely SMARCA2 and SMARCA4), it led to physical compaction of enhancer elements^4^. This hindered transcription factor access and downstream activation of the oncogenic transcriptional programs regulated by AR, FOXA1, ERG, and MYC and led to inhibition of CRPC tumor growth^4^.

In addition to the physical remodeling of nucleosomes, chemical modifications to histones and the transcriptional machinery are essential for the activation of enhancers^10^. Two critical regulators involved in fueling enhancer activity in mCRPC are the paralog acetyltransferases p300 and CBP^11,12^. p300 and CBP function as critical coactivators of enhancer elements through orchestrating acetylation of histones and other transcriptional regulators, recruiting transcriptional machinery, and acting as scaffolds for protein complex assembly^13-15^. These enzymes are increasingly recognized as coregulators of AR^16-19^, and their inhibition significantly hinders the growth of AR-positive prostate cancer cells, including CRPC^17,20,21^. Despite this, the exact interplay between AR and p300/CBP within the AR enhanceosome remains elusive. Previously, two major types of p300/CBP inhibitors have been developed that target one of two domains of the proteins - inhibitors of the reader bromodomain or histone acetyltransferase (HAT) domain^17,20,21^. Small-molecule inhibitors of the reader bromodomain prevent CBP/p300 from recognizing acetylated lysine residues through their bromodomains, while HAT inhibitors attenuate their acetyltransferase activity^17,20,21^. Studies have shown that both domains are required for complete p300/CBP activities; therefore, combination therapy to target both domains can achieve significant synergistic effects^22^. Although reader bromodomain or HAT inhibitors can markedly alter the chromatin and transcriptional landscape and inhibit cell growth, both classes exhibit limitations. Studies have indicated that reader bromodomain inhibitors are only effective against a limited range of substrates acetylated by p300/CBP^12,23^, while the recently developed p300/CBP selective HAT domain inhibitor, A485, exhibited poor drug-like characteristics and modest effects on suppression of cancer cell proliferation^20,24^. Currently, there is only one p300/CBP bromodomain inhibitor in clinical trials, CCS1477^17^.

Here, we report that p300 and CBP are requisite HATs that define the active AR enhanceosome and hyperacetylate histone H2B N-terminus (H2BNTac) to drive a unique oncogenic transcriptome and promote cancer cell proliferation. Consistently, H2BNTac is significantly elevated in prostate cancer lesions compared to normal prostate epithelia. Employing an orally bioavailable p300/CBP PROTAC degrader, CBPD-409, with excellent selectivity and potency, we demonstrate that specific degradation of p300/CBP can achieve superior suppression of AR-positive prostate cancer cell growth relative to p300/CBP bromodomain inhibitors and HAT domain inhibitors. This enhanced efficacy is attributed to the stronger suppression of AR signaling and the targeted inhibition of a distinct set of oncogenes critical for cell growth regulation, which are dependent on H2BNTac. Intriguingly, our data suggests that degradation of p300/CBP is a safe approach for prostate cancer therapy without causing any evident toxicity, including thrombocytopenia and gastrointestinal tract toxicities. Collectively, our data underscore the significant impact of p300/CBP on the progression of prostate cancer and emphasize the potential of p300/CBP PROTAC degraders as a promising therapeutic approach.

## Results

### P300/CBP-catalyzed H2BNTac is significantly elevated in prostate cancer lesions compared to normal prostate epithelia

To identify which histone post-translational modifications (PTMs) were reprogrammed and possibly involved in prostate cancer development, we examined a range of histone PTMs related to transcriptional regulation in matched benign and prostate cancer tissues from patients^25-29^. In contrast to other histone modifications, such as H3K4me1, H3K4me3, H3K27me3, H3K27ac, and H3K18ac, there was a notable increase in a specific set of H2BNTac PTMs in prostate cancer tissues (**Figure S1A**). H2BNTac, which includes the recently characterized H2BK5ac, H2BK12ac, H2BK16ac, and H2BK20ac, is known to mark active intergenic enhancer regions^27^. Strikingly, immunofluorescent staining of prostatectomy samples revealed a marked increase in H2BK5ac and H2BK20ac levels in KRT8-positive malignant luminal cells relative to patient-matched adjacent benign epithelia (**Figures 1A-B and S1B**). Similar findings were observed in isogenic mouse organoid models. Compared to wild-type (WT) mouse prostate organoids, those genetically altered to inactivate the *Pten* or *Trp53* genes exhibited elevated levels of H2BNTac (**Figure 1C**). Importantly, from the DepMAP database, cumulative analysis of essentiality scores for all histone acetyltransferases (HATs) and histone deacetylases (HDACs) in AR-positive prostate cancer cell lines (namely VCaP and LNCaP) identified p300 as the most essential gene in these cell lines (**Figure 1D**). Considering recent findings that H2BNTac is exclusively dependent on p300/CBP^27^, we carried out a comprehensive analysis of H2BK5ac, H2BK20ac, H3K18ac, H3K27ac, and p300 genomic distribution in VCaP cells. This analysis revealed that a significant majority of H2BK5ac (82.3%) and H2BK20ac (76%) sites were co-localized with p300 peaks. In contrast, a smaller fraction of H3K18ac (46.7%) and H3K27ac (46.3%) sites showed overlap with p300, further supporting the specific dependency of H2BNTac on p300 binding and enzymatic activity in prostate cancer cells (**Figures 1E & S1C**). These data highlight the significance of the p300/H2BNTac epigenetic axis in the survival of AR-positive prostate cancer cells.

**Figure 1:**
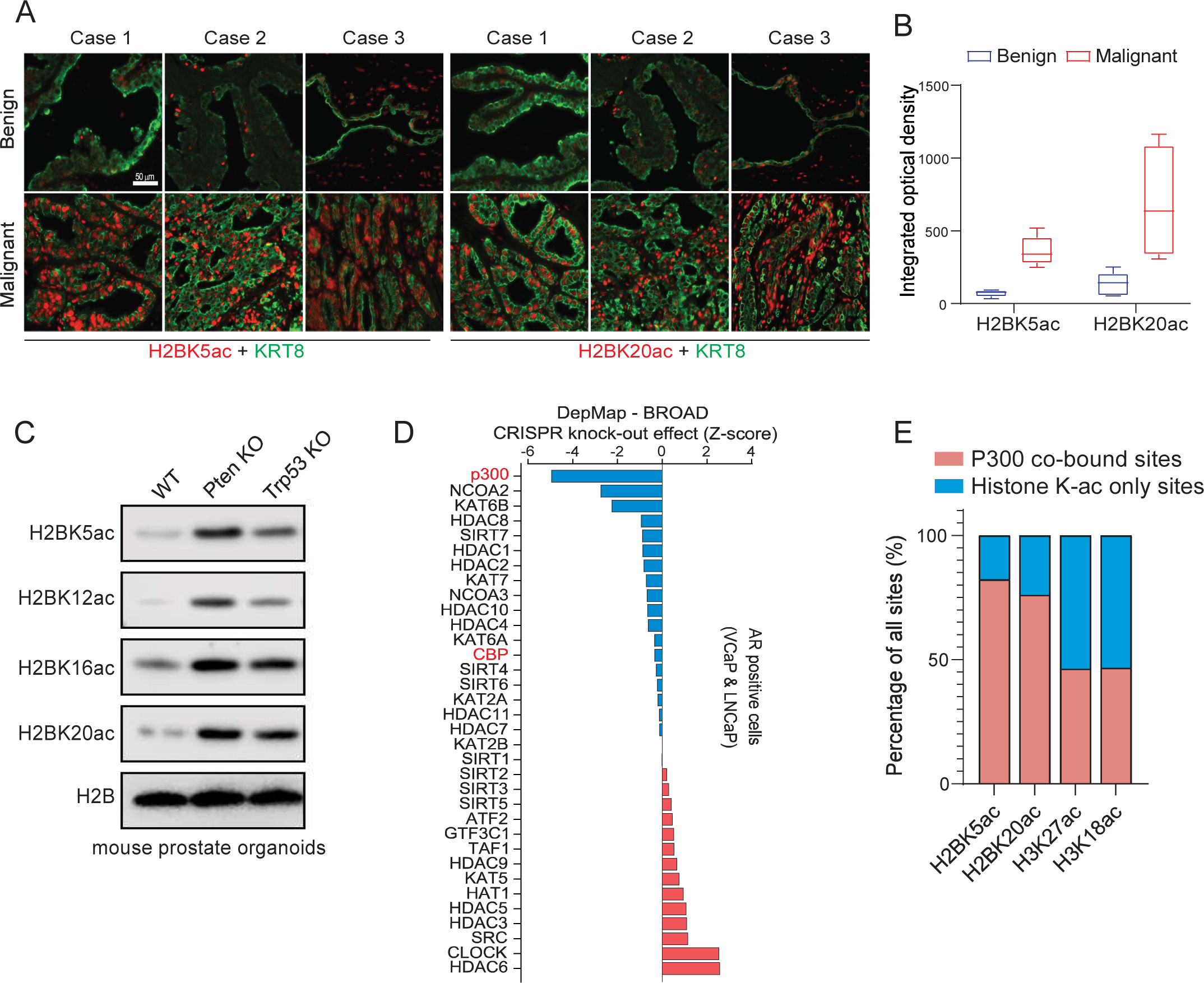
P300/CBP-dependent H2BNTac is significantly elevated in prostate cancer lesions compared to normal prostate epithelia. **a.** Representative multiplex immunofluorescence (IF) images of H2BK5ac (red)/KRT8 (green), and H2BK20ac (red)/KRT8 (green) staining in patient-matched adjacent benign and tumor tissues. Magnification: 200x. Scalebar = 50 µm. **b.** H2BK5ac and H2BK20ac IF mean intensity per field (integrated optical density) from images in panel a. n=5 (two-sided t-test). **c.** Immunoblot analysis of indicated H2BNTac in wild-type (WT), *Pten* KO, and *Trp53* KO mouse organoids. **d.** Cumulative DepMap CRISPR knockout essentiality scores of histone acetyltransferases and deacetylases in AR-positive prostate cancer cell lines. Aggregated z-scores for each gene derived from LNCaP and VCaP DepMap screens are shown, with p300 and CBP highlighted in red. **e.** Bar charts illustrating overlaps of genome-wide p300, H2BK5ac, H2BK20ac, H3K18ac, and H3K27ac ChIP-seq peaks in VCaP cells.

### P300 is a determinant cofactor of the activated AR enhanceosome in prostate cancer

As a transcriptional coactivator, p300 predominantly co-localizes with transcription factors at specific chromatin regions, playing a pivotal role in the regulation of gene expression^14,30,31^. We explored the interplay between p300 and AR or ERG, which are the two most critical oncogenic transcription factors in prostate cancer cells, in AR-positive/ERG-positive VCaP cells^32^. Based on normalized read densities, we categorized ChIP-seq peaks of prostate cancer oncogenic transcription factors (AR, ERG, and FOXA1) and transcriptional cofactors (SMARCA4, p300, and BRD4) at non-promoter regions into quartiles (Q1-Q4, where Q4 represents the top quartile and Q1 the bottom quartile). Subsequently, we assessed their presence at the binding sites of AR or ERG. As expected, the majority of the strongest AR or ERG peaks (Q4 peaks) were co-occupied by pioneer transcription factor FOXA1^7^ and the ATPase subunit of the SWI/SNF complex SMARCA4^4^, which are key components of AR and ERG transcriptional complexes (**Figures 2A and S1D-E**). Notably, we found p300 co-localized with over 60% of the strongest AR and ERG Q4 peaks (**Figures 2A and S1E**). In contrast, another cofactor, BRD4^33^, only bound a small fraction of AR (approximately 30%) and ERG (approximately 40%) Q4 binding sites (**Figures 2A and S1E**). Furthermore, unlike BRD4, the majority of the p300 peaks that overlapped with AR were of high confidence and signal strength and belonged to quartiles Q3 and Q4. These findings suggest that p300 acts as a critical cofactor for AR and ERG transcriptional machineries.

**Figure 2:**
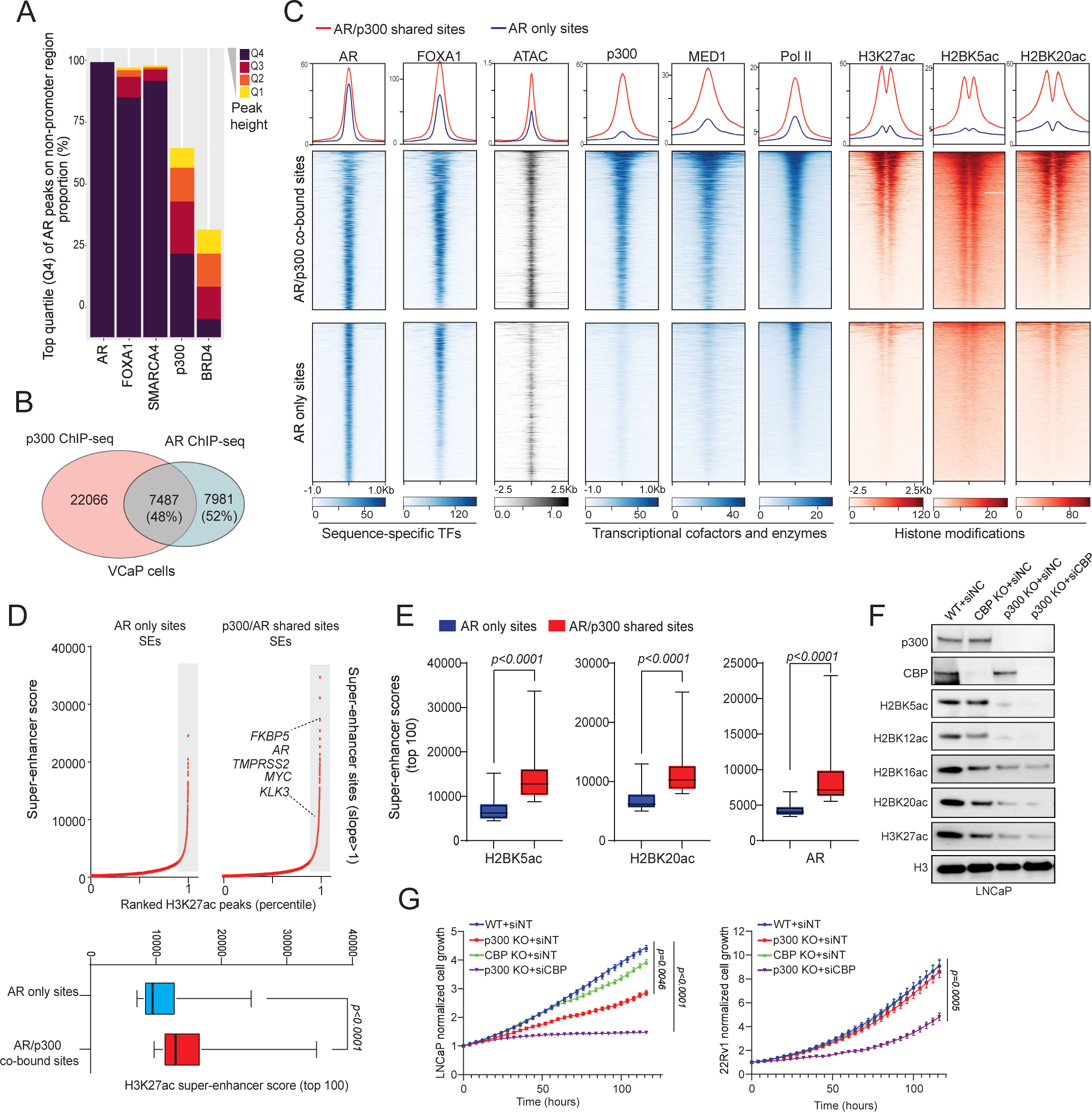
P300 is a determinant cofactor of the activated AR enhanceosome in prostate cancer. **a.** Proportion of FOXA1, SMARCA4, p300, and BRD4 ChIP-seq peaks that map to the top quartile of AR ChIP-seq peaks located in non-promoter regions in VCaP cells. **b.** Venn diagrams illustrating overlaps of genome-wide p300 and AR ChIP-seq peaks in VCaP cells. **c.** ChIP-seq and ATAC-seq read-density heatmaps at AR/p300 co-bound and AR only binding sites in VCaP cells. Transcription factors (TFs), transcription cofactors, RNA Pol II, and respective histone marks indicated. **d.** Ranked H3K27ac ChIP-seq signal of super-enhancers (SEs) on AR/p300 co-bound sites and AR only sites in VCaP cells identified by HOMER. Bottom panel: Activity scores of super-enhancers comprising either AR/p300 co-bound or AR only sites (two-sided t-test). **e.** Bar charts depicting the H2BK5ac, H2BK20ac, and AR super-enhancer (SEs) scores at AR only or AR/p300 co-bound sites (two-sided t test). **f.** Immunoblot analysis of p300, CBP, and the indicated histone marks in LNCaP WT (wild-type), p300 KO (knock-out), CBP KO, and p300 KO with siCBP (small interfering RNA) cells. **g.** Incucyte live-cell analysis of LNCaP and 22Rv1 cells with respective CRISPR KO and siRNAs indicated. siNT, non-targeting siRNA. Data are presented as mean +/− SD (n = 6). (two-sided t-test)

To validate this model, we profiled the p300 cistrome in prostate cancer cells and classified AR binding sites into shared (AR/p300 co-bound) and exclusive (AR only) categories (**Figure 2B**). While the binding intensity of AR was similar between the two groups, AR/p300 co-bound elements that comprised 48% of the AR cistrome were characterized by higher chromatin accessibility as well as parallel binding of the downstream transcriptional machinery, such as MED1 and Pol II (**Figure 2C**). Consistently, these enhancer sites also showed higher accumulation of active histone marks, namely H3K27ac, H2BK5ac, and H2BK20ac (**Figure 2C**). In contrast, enhancer sites that were accessed by AR without parallel loading of p300 had lower accessibility and significantly lower abundance of transcriptional cofactors and active histone acetylation marks (bottom heatmaps, **Figure 2C**). Super-enhancers comprise dense and closely arranged enhancer clusters that amplify the expression of dominant oncogenes in cancer cells^34^. Compared to AR only enhancers, we found the AR/p300 co-bound sites to more frequently comprise super-enhancer clusters, including the ones within recurrently rearranged loci such as *TMPRSS2*, *AR*, and *MYC* (**Figures 2D-E**). Similar analyses focusing on the intersection of p300 and ERG binding sites demonstrated that p300 also played a critical role in determining the activity of the ERG transcriptional complex (**Figure S1F**).

CRISPR/Cas9 and siRNA techniques were next employed to target p300 and its paralog, CBP, to understand their specific contribution to the survival and growth of prostate cancer cells. Notably, in LNCaP cells, p300 inhibition markedly reduced histone H3 and H2B acetylation as well as impeded cell proliferation, while simultaneous targeting of both p300 and CBP led to an even more pronounced loss of histone acetylation with subsequent attenuation of cell proliferation (**Figures 2F-G**). In 22Rv1 cells, inhibition of both p300 and CBP was necessary to suppress histone acetylation and cell growth (**Figures 2G and S1G**). This observation suggests a compensatory functional relationship between the two paralogs in prostate cancer cells, highlighting the need to target both enzymes to achieve optimal suppression of histone acetyltransferase activity and inhibition of cell growth.

### Degradation of p300/CBP, but not inhibition of their reader bromodomains, abolishes global H2BNTac

As mentioned above, the p300/CBP reader bromodomain inhibitor CCS1477 is currently in early-phase clinical trials; given this, we tested CCS1477 and another reader bromodomain inhibitor, GNE-049, for their ability to attenuate p300/CBP activity in AR-positive prostate cancer cells^17,21^. Surprisingly, while both CCS1477 and GNE-049 significantly decreased the p300/CBP-catalyzed H3K27ac mark, we saw modest to no decrease in the abundance of H2BNTac in the treated prostate cancer cells (**Figures 3A and S2A**), suggesting that reader bromodomain inhibitors lead to only a partial inhibition of p300/CBP activity. Thus, utilizing the bromodomain inhibitor GNE-049 as a warhead, we synthesized a novel cereblon (CRBN)-dependent proteolysis targeting chimera (PROTAC) degrader of p300/CBP, called CBPD-409^35^ (**Figure 3B**). In multiple cell lines, CBPD-409 exhibited marked potency in degrading both p300 and CBP proteins within an hour (**Figures 3C and S2B-E**). Mass spectrometry-based proteomics analysis in prostate cancer cell lines confirmed that CBPD-409 specifically and most significantly degraded p300 and CBP from over 7000 detectable proteins (**Figures 3D and S2F**). No changes in the abundance of other bromodomain-containing proteins were detected, including the BET family of proteins^36^ (**Figures 3E and S2G**), or in the level of known CRBN neo-substrates, including GSPT1 and Ikaros^37^ (**Figure S2H**). Further confirming the PROTAC’s mechanism of action, competition with the free thalidomide ligand or proteasomal inhibition using carfilzomib completely blocked degradation of p300/CBP by CBPD-409 (**Figures S2I-J**). The inactive isoform of CBPD-409, called CBPD-409-Me, with a methyl substitution in the cereblon ligand moiety did not degrade its targets (**Figures S2K-L**). Notably, compared to previously reported p300/CBP PROTAC degraders, dCBP1^38^ and JQAD1^39^, CBPD-409 exhibited enhanced degradation effects in prostate cancer cell lines (**Figure 3F**).

**Figure 3:**
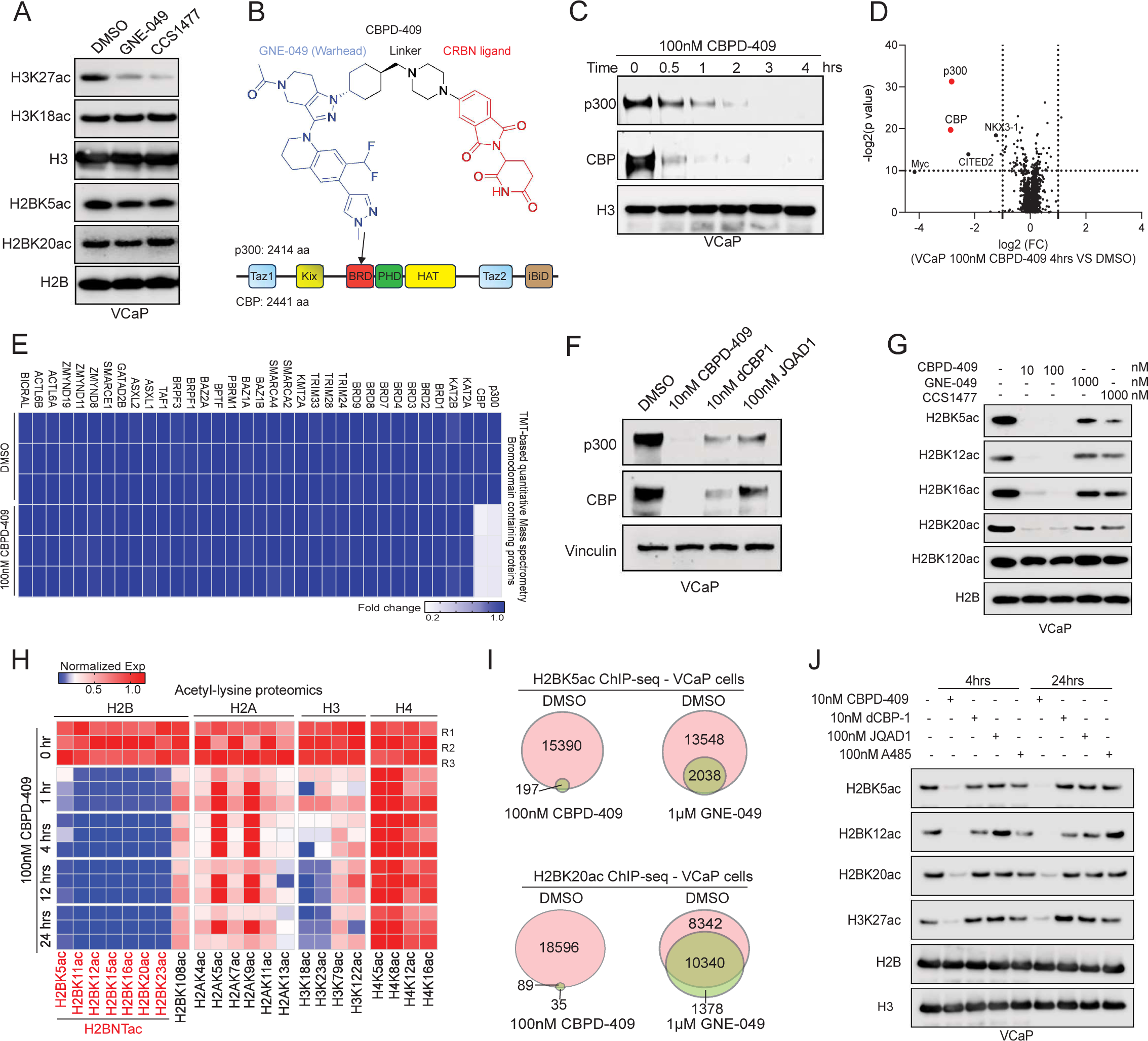
Degradation of p300/CBP, but not inhibition of their reader bromodomains, abolishes global H2BNTac. **a.** Immunoblot analysis of indicated histone acetylation marks in VCaP cells treated with DMSO, 1 µM GNE-049, or 1 µM CCS1477 for 24 hours. **b.** Structure of CBPD-409 and schematic of p300 and CBP domains. CBPD-409-targeted bromodomain (BRD) is highlighted by the arrow. Key domains of p300/CBP proteins are noted. Taz1: TAZ zinc finger domain; Kix: Kinase-inducible domain (KID) interacting domain; PHD: Plant homeodomain; HAT: Histone acetyltransferase domain; iBiD: N-terminal interferon-binding domain. CRBN, cereblon. **c.** Immunoblot analysis of p300 and CBP in VCaP cells treated with 100 nM CBPD-409 for the indicated durations. **d.** TMT (tandem mass tag) mass spectrometry assay to evaluate effects of CBPD-409 (100 nM, 4 hours) on the proteome of VCaP cells. Data are plotted as log2 of the fold change (FC) versus DMSO control against -log2 of the p-value per protein from n = 3 independent experiments. All t-tests performed were two-tailed t-tests assuming equal variances. **e.** Heatmap of relative abundance of all bromodomain-containing proteins detected in TMT-based quantitative MS after 4 hours CBPD-409 treatment of VCaP cells. **f.** Immunoblot analysis of p300 and CBP in VCaP cells treated with CBPD-409, dCBP-1, and JQAD1 at the indicated concentrations for 4 hours. **g.** Immunoblot analysis of labeled H2B N-terminus acetylation in VCaP cells treated with the indicated concentration of CBPD-409, GNE-049, or CCS1477 for 4 hours. **h.** Acetyl-lysine proteomics analysis of VCaP cells. Fold change heatmap illustrating alterations in acetyl-lysine levels of core histone proteins (H2A, H2B, H3, and H4) in VCaP cells treated with 100 nM CBPD-409 at indicated time points, compared to DMSO vehicle treated cells. Data plotted from n=3 independent samples. **i.** Venn diagrams of genome-wide changes of H2BK5ac and H2BK20ac ChIP-seq peaks after CBPD-409 (100 nM 4 hours) or GNE-049 (1 µM 4 hours) treatment of VCaP cells. **j.** Immunoblot analysis of indicated histone marks in LNCaP cells treated with 10 nM CBPD-409, 10 nM dCBP-1, 100 nM JQAD1, or 100 nM A485 for 4 or 24 hours.

Degradation of p300/CBP with CBPD-409 abolished H3K27ac as well as the H2BNTac marks in prostate cancer cell lines, without affecting the C-terminal H2BK120ac mark (**Figures 3G and S3A-B**). In contrast, neither GNE-049 or CCS1477 extinguished H2BNTac levels, even when 10-fold higher concentrations were used (**Figure 3G**). Immunoprecipitation (IP)-based mass spectrometry analysis focused on the acetylation of core histone proteins was employed to profile the repertoire of all histone marks that were catalyzed by p300/CBP. Here, we uncovered several acetyl marks on histone tails that were catalyzed by p300/CBP, including H2BNTac, which were rapidly lost within one hour of CBPD-409 treatment (**Figures 3H and S3C-D**). Using ChIP-seq, we next explored the specific effects of CBPD-409 on chromatin distribution of H2BNTac marks (namely H2BK5ac and H2BK20ac) and compared it to the changes seen upon treatment with GNE-409. CBPD-409 resulted in a complete loss of H2BK5ac and H2BK20ac peaks (>98% of peaks lost) on the chromatin, while GNE-049 showed only a modest impact on HBK5ac and H2BK20ac distribution across the chromatin (**Figures 3I and S3E**). Strikingly, CBPD-409 exhibited the strongest inhibitory effects in suppressing H2BNTac and H3K27ac compared to the HAT inhibitor A485 and p300/CBP degraders dCBP-1 and JQAD1 (**Figure 3J**). Altogether, these findings suggest that p300/CBP retain partial catalytic function in the presence of reader bromodomain inhibitors and highlight the efficacy of CBPD-409-triggered rapid p300/CBP degradation in completely extinguishing their oncogenic histone acetylation program in prostate cancer cells.

### Degradation of p300/CBP abolishes histone acetylation activity at the AR enhanceosome

While CBPD-409 and GNE-049 triggered a similar decrease in overall abundance of H3K27ac histone modifications (**Figures S4A-B**), we detected marked differences when interrogating the changes at AR binding sites. Unlike GNE-049, CBPD-409-triggered p300/CBP degradation caused an almost complete loss of H3K27ac at AR binding sites within just four hours of treatment (**Figure 4A**). This was accompanied by a parallel loss of H2BNTac at the same sites (**Figure 4A**). On the other hand, GNE-049 treatment resulted in only a modest suppression of H3K27ac and H2BNTac at AR enhancers (**Figure 4A**). Concordant findings were observed at AR super-enhancers, where CBPD-409 was found to more effectively repress H3K27ac, H2BK20ac, and H2BK5ac levels compared to GNE-049 (**Figure 4B**). p300/CBP degradation also severely attenuated H3K27ac at CRPC-specific AR enhancers defined using AR cistromes from patient tissues^5,6^ (**Figure S4C**). Altogether, these findings suggest that degradation of p300/CBP rapidly depletes active histone acetylation marks at AR enhancers in prostate cancer cells.

**Figure 4:**
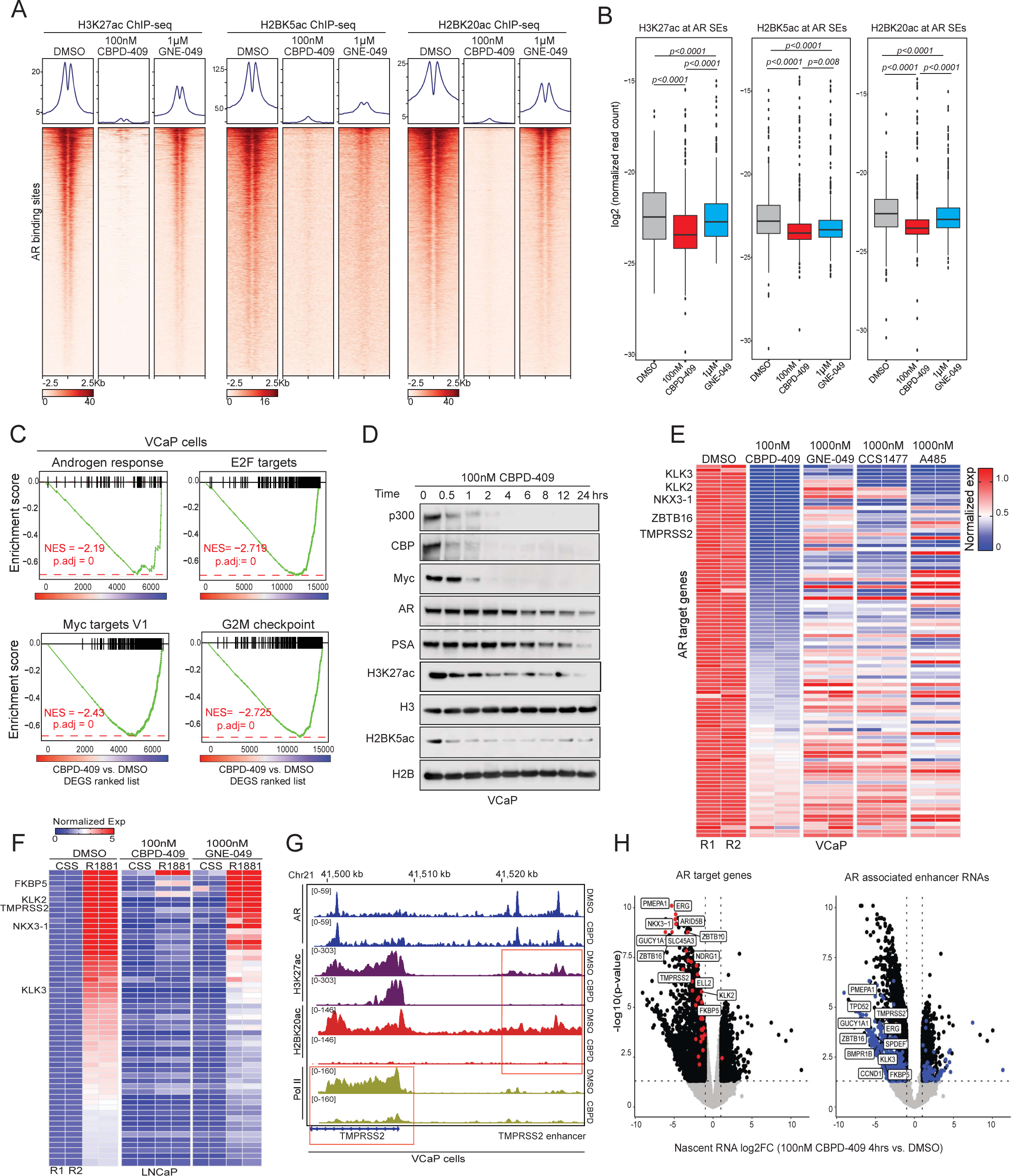
Degradation of p300/CBP abolishes histone acetylation activity at AR enhanceosome. **a.** ChIP-seq read-density heatmaps of H3K27ac, H2BK5ac, and H2BK20ac at AR *cis*-regulatory elements in VCaP cells with 4 hours of 100 nM CBPD-409 or 1 µM GNE-049 treatment. **b.** Normalized ChIP-seq read density of H3K27ac, H2BK5ac, and H2BK20ac at super-enhancers associated with AR binding sites, based on AR ChIP-seq data in VCaP cells with 4 hours of 100 nM CBPD-409 or 1 µM GNE-049 treatment (Wilcox test). **c.** Gene set enrichment analysis (GSEA) plots for AR, MYC, E2F, and G2M checkpoint pathway-related genes using the fold change rank-ordered gene signature from the CBPD-409 treated VCaP cells. NES, net enrichment score; adj P, adjusted p-value; DEGS, differentially expressed genes. n=2 biological replicates. **d.** Immunoblot analysis of indicated proteins and histone marks in VCaP cells treated with 100 nM CBPD-409 for the indicated times. **e.** RNA-seq heatmaps for classical AR target genes in VCaP cells treated with 100 nM CBPD-409, 1 µM GNE-049, 1 µM CCS1477, or 1 µM A485 for 24 hours. **f.** RNA-seq heatmaps for classical AR target genes (defined as upregulated by 1 nM R1881 in 12 hours) in LNCaP cells treated with 100 nM CBPD-409 or 1 µM GNE-049. Cells were cultured in medium with 10% charcoal-stripped FBS (CSS) overnight, then pre-treated with CBPD-409 or GNE-049 for 1 hour and subsequently stimulated with 1 nM R1881 for 12 hours. **g.** ChIP-seq tracks of AR, H3K27ac, H2BK20ac, and RNA Pol II within the *TMPRSS2* gene loci in VCaP cells treated with or without 100 nM CBPD-409 for 4 hours. **h.** Volcano plots of coding genes nascent RNA expression (left panel) and enhancer RNAs (eRNAs) expression (right panel) in VCaP cells treated with 100 nM CBPD-409 for 4 hours. The AR target genes which were significantly suppressed by CBPD-409 are highlighted by red in the left panel. The AR-associated eRNAs which were significantly suppressed by CBPD-409 are highlighted by blue in the right panel. The names of key AR target genes (left panel) or eRNAs associated target genes (right panel) are labeled. The data were generated from the 5-ethynyluridine (EU) labeled nascent RNA-seq. The volcano plots represent the log2 fold change (FC) of CBPD-409 relative to DMSO plotted against the - log10 p-value for each gene, based on n=2 independent experiments. Statistical tests were two-tailed t-tests with the assumption of equal variances.

Beyond histone acetylation, the regulation of chromatin accessibility and transcription factor loading also play crucial roles in enhancer-mediated transcription^40,41^. To understand the effects of p300/CBP protein loss on chromatin accessibility, we employed transposase-accessible chromatin sequencing (ATAC-seq). In contrast to the SWI/SNF ATPase degrader, AU-15330^4^, CBPD-409 treatment did not alter chromatin accessibility (**Figure S4D**). We also detected no change in the chromatin binding of AR or FOXA1 upon degradation of p300/CBP (**Figures S4D-E**), suggesting that p300/CBP degradation does not alter transcription factor access to chromatin.

Gene enrichment analysis (GSEA) based on RNA-seq data revealed that p300/CBP degradation led to a marked suppression of androgen response and proliferation-related pathways, including E2F, MYC, G2M checkpoint, and mTORC1 signaling (**Figures 4C and S5A-B**). We also observed strong inhibition of other prostate cancer oncogenic signaling, including ERG and FOXA1, after CBPD-409 treatment (**Figure S5B**). CRISPR/Cas9-mediated knockout of both p300 and CBP in LNCaP cells phenocopied the PROTAC in significantly suppressing the expression of AR and MYC target genes (**Figure S5C**). CBPD-409 also significantly reduced the protein levels of AR, PSA, and MYC in a time-dependent manner (**Figure 4D**). MYC and NKX3.1 (an AR target gene) proteins were amongst down-regulated proteins in our proteomics data generated from CBPD-409-treated VCaP cells (**Figure 3D**). Notably, consistent with complete attenuation of p300/CBP functions, the inhibitory effects of CBPD-409 were more pronounced when compared to treatment with the reader bromodomain inhibitors or HAT domain inhibitor (**Figures 4E and S5D**). Remarkably, unlike GNE-049, one hour pre-treatment with CBPD-409 completely attenuated ligand-induced transcriptional activity of AR in prostate cancer cells (**Figure 4F**). This was paralleled by marked reduction in loading of Pol II specifically at the promoters of AR up-regulated genes only in CBPD-409 treated cells (**Figures 4G and S5E-F**), without affecting Pol II loading and transcription at randomly selected genes (**Figure S5F**). To assess the immediate impact of CBPD-409 on active transcription in prostate cancer cells, we next performed nascent RNA-seq (5-ethynyluridine sequencing) and discovered hallmark AR-regulated transcripts amongst the topmost significantly down-regulated genes (**Figures 4H and S5G**). Notably, a large proportion of the significantly down-regulated nascent transcripts were comprised of enhancer RNAs templated from AR binding *cis*-regulatory elements, suggesting that degrading p300/CBP impairs their transcriptional function (**Figure 4H**). In summary, p300/CBP degradation triggers a rapid loss of acetylation marks at AR enhancer-flanking nucleosomes and disrupts the subsequent transcription of enhancer elements as well as their distal target genes.

### p300/CBP degradation leads to stronger inhibition of oncogenic gene programs compared to bromodomain inhibition

To identify the specific transcriptional effects of p300/CBP degradation, we carried out comparative analysis of the global transcriptomes from cells treated with CBPD-409 or reader bromodomain inhibitors, GNE-409 and CCS1477. Strikingly, over 45% of the genes downregulated by CBPD-409 were not comparably suppressed by reader bromodomain inhibitors (**Figure S6A**), with p300/CBP degradation showing a stronger inhibition of gene expression (**Figure 5A**). GSEA revealed unique CBPD-409 repressed genes to be associated with cell cycle and growth-associated signaling pathways in prostate cancer cells, which included the androgen response signature (**Figures 5B-C and S6B-C**). Amongst genes uniquely down-regulated by CBPD-409 versus reader bromodomain inhibitors, we identified *NKX3-1*, *CITED2*, and *CCND1* that are known for their driver roles in prostate cancer progression^42-44^ (**Figures 5A and S6D**). We were able to confirm the contrasting effects on these genes at the protein level, where only treatment with CBPD-409 entirely abolished their expression compared to GNE-049 (**Figures 5D and S6E-F**). More impressively, co-treatment with free thalidomide, which blocks the degradation activity of CBPD-409 and renders it to function as a bromodomain inhibitor, reversed the loss in H2BNTac as well as NKX3-1, CITED2, and CCND1 expression in VCaP cells (**Figures 5E-F**). In the same experiment, phenocopying the effect of reader bromodomain inhibitors, combinatorial treatment with CBPD-409 and thalidomide still triggered the loss of bromodomain-dependent H3K27ac and repressed MYC expression (**Figures 5E-F**), suggesting the residual H2BNT acetyltransferase activity of bromodomain-inhibited p300/CBP continues to support the expression of important cancer-promoting genes. As an orthogonal approach, treatment of prostate cancer cells with the inactive analogue of CBPD-409, CBPD-409-Me (i.e., the dead PROTAC) significantly reduced H3K27ac, yet only had modest effects on H2BNTac histone modifications and failed to repress the expression of degradation-specific gene targets (**Figure 5G**). A closer inspection of the *NKX3-1* and *CCND1* loci confirmed a significant depletion of H2BK20ac and Pol II loading at these genes upon treatment with CBPD-409 without affecting H3K27ac, which was not observed upon treatment with GNE-049 (**Figure S6G**). Notably, compared to other p300/CBP PROTAC degraders and the HAT domain inhibitor, CBDP-409 exhibited the most prominent effects in repressing NKX3-1, CITED2, and CCND1 levels (**Figure 5H**). Altogether, our findings suggest that p300/CBP complexes retain their acetyltransferase activity for the H2BNT lysine residues despite the attenuation of the reader bromodomain, which sustains AR enhancer activity and triggers a distinct oncogenic gene program in prostate cancer cells.

**Figure 5:**
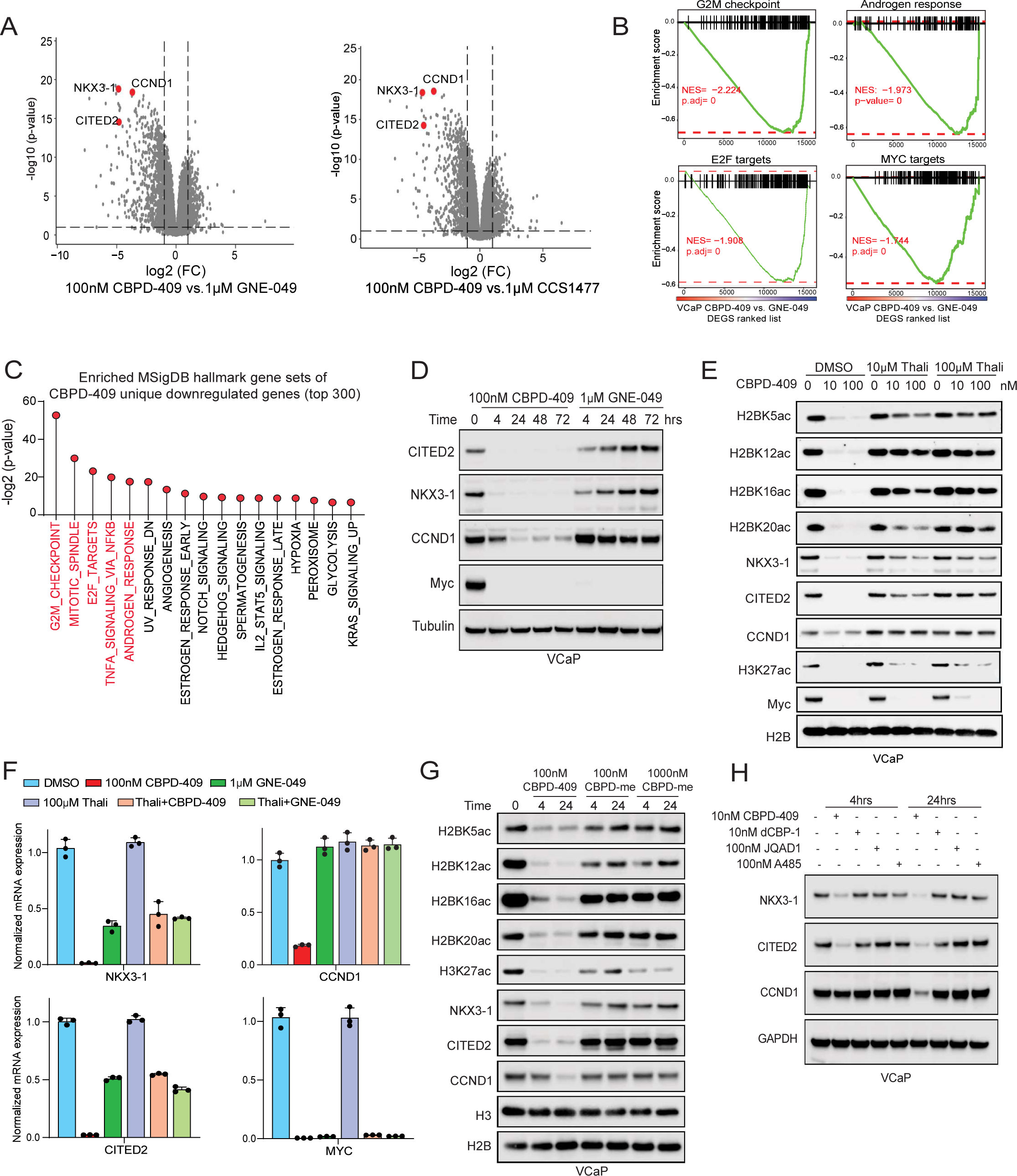
p300/CBP degradation leads to stronger suppression of oncogenic gene programs compared to bromodomain inhibition. **a.** Volcano plot of gene expression in VCaP cells treated for 24 hours with either 100 nM CBPD-409, 1 µM GNE-049, or 1 µM CCS1477. A set of genes, including *CCND1*, *CITED2*, and *NKX3-1*, are exclusively repressed by CBPD-409. The data represent the log2 fold change (FC) of CBPD-409 relative to GNE-049 or CCS1477 plotted against the -log10 p-value for each gene. Based on n=2 independent experiments. Statistical tests were two-tailed t-tests with the assumption of equal variances. **b.** GSEA plots for G2M checkpoint, AR, E2F, and MYC signaling pathways, based on rank-ordered fold change gene signatures in VCaP cells: comparison of 24 hours treatment with 100 nM CBPD-409 vs. 1 µM GNE-049. DEG: differentially expressed gene. **c.** Uniquely down-regulated genes in CBPD-409 relative to GNE-409 treated VCaP cells analyzed for overlap with MSigDB hallmark gene sets. The top five hallmark gene sets (ranked by p-value) are highlighted in red. **d.** Immunoblot analysis of indicated proteins in VCaP cells treated with 100 nM CBPD-409 or 1 µM GNE-049 for indicated times. **e.** Immunoblot analysis of labeled proteins and histone marks in VCaP cells pre-treated with different concentrations of thalidomide (Thali) for 1 hour, then treated with CBPD-409 at indicated concentrations for 4 hours. **f.** Quantitative-PCR (qPCR) of *NKX3-1*, *CCND1*, *CITED2*, and *MYC* expression in VCaP cells pre-treated with 100 µM thalidomide for 1 hour, then treated with CBPD-409 or GNE-409 for 4 hours. **g.** Immunoblot analysis of indicated proteins and histone marks in VCaP cells treated with CBPD-409 or CBPD-409-Me (inactive analogue) for indicated times. **h.** Immunoblot analysis of indicated proteins in VCaP cells treated with 10 nM CBPD-409, 10 nM dCBP-1, 100 nM JQAD1, or 100 nM A485 for noted durations.

### p300/CBP degradation inhibits prostate cancer growth without apparent toxicities

In line with the above molecular insights, treatment with CBPD-409 resulted in stronger cytotoxicity (IC50 ranging between 2 nM to 11 nM) in all tested AR-positive prostate cancer cell lines relative to GNE-049 (IC50 ranging between 650 nM to 1900 nM; **Figures 6A and S7A-B**). Notably, despite target degradation, CBPD-409 showed no efficacy in AR-negative prostate cancer (PC3 and DU145), neuroendocrine prostate cancer (NCI-H660 and LTL-331R-CL), or normal human prostate-derived cell lines (WPMY-1, PNT2, and RWPE1; **Figures 6A and S7C-D**). CBPD-409 also exhibited superior cytotoxicity, i.e. had the lowest IC50, when compared to other published p300/CBP degraders, bromodomain inhibitors, or HAT domain inhibitors in AR-positive prostate cancer cell lines (**Figure 6B**). As expected, the inactive PROTAC CBPD-409-Me had a similar cytotoxicity profile as GNE-049 (**Figure S7E**). Notably, CBPD-409 exhibited strong cytotoxicity in bromodomain inhibitor (GNE-049) and HAT domain inhibitor (A485) resistant LNCaP cells (**Figures S7F-G**).

**Figure 6:**
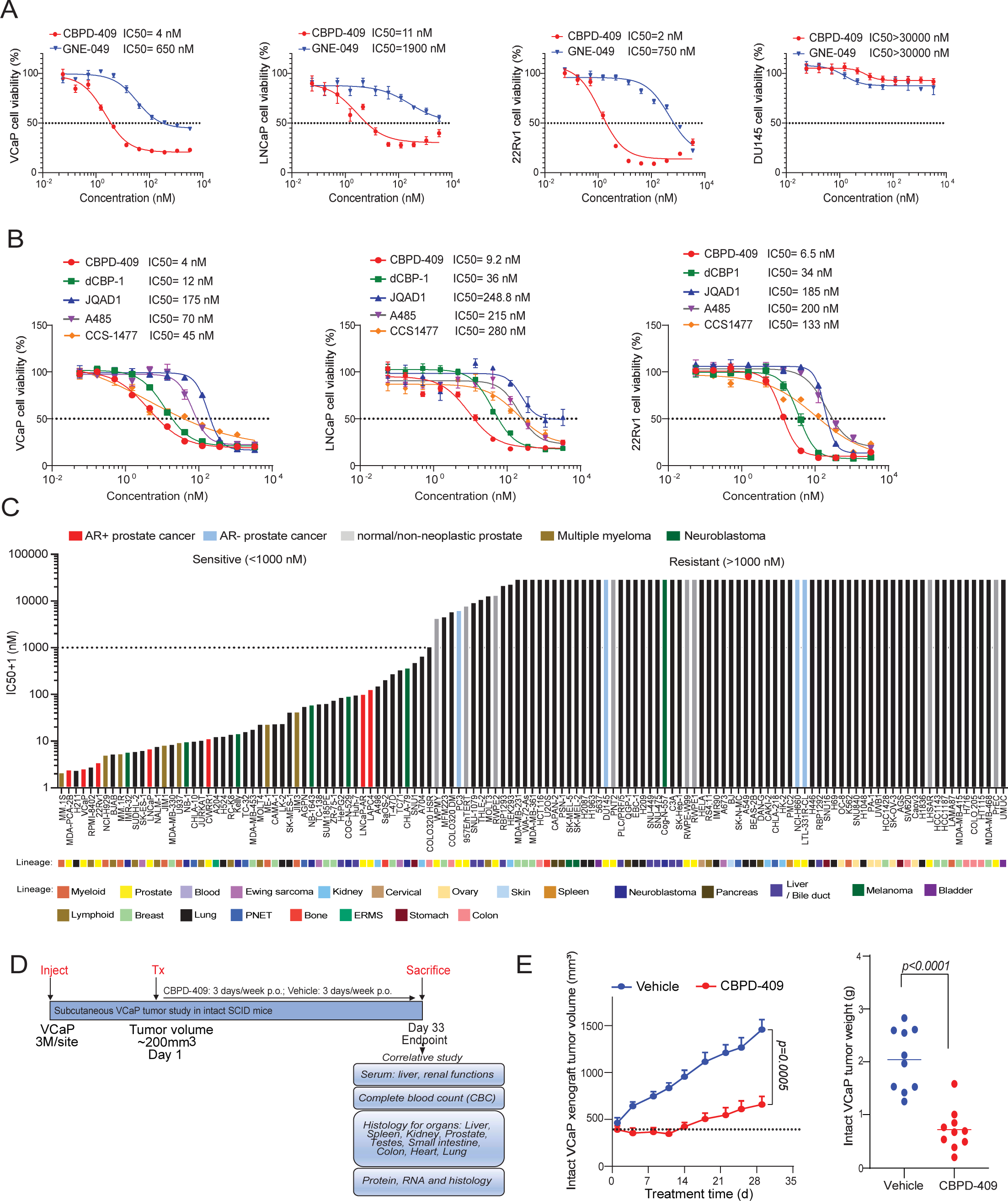
p300/CBP degradation inhibits prostate cancer growth. **a.** Dose-response curves and IC50 of prostate cancer cells treated with CBPD-409 and GNE-049. Data are presented as mean +/− SD (n = 6). **b.** Dose-response curves and IC50 of prostate cancer cells treated with p300/CBP degraders (CBPD-409, dCBP-1, and JQAD1), bromodomain inhibitor CCS1477, and HAT inhibitor A485. Data are presented as mean +/− SD (n = 6). **c.** Rank-order plot of IC50 values for CBPD-409 across 131 human normal and cancer cell lines following 5 days of treatment. Models of AR-positive prostate cancer, AR-negative prostate cancer, non-neoplastic prostatic cells, multiple myeloma, and neuroblastoma are highlighted in specific colors. The originating tissue lineages are indicated below. PNET, primitive neuroectodermal tumor; EMRS, embryonal rhabdomyosarcoma. **d.** Schematic of the CBPD-409 *in vivo* efficacy study in intact VCaP xenograft model. **e.** Graphs depicting the tumor volume curve (left) and tumor weights (right) in the intact VCaP xenograft model, measured biweekly by calipers. Treatments included vehicle and CBPD-409 (3 mg/kg, administered orally three days per week). Statistical analysis was performed using a two-sided t-test. Data are presented as mean ± SEM. Sample sizes are as follows: vehicle, n = 10; CBPD-409, n = 10.

More impressively, assessment of CBPD-409’s cytotoxicity in a large panel of 131 normal and cancer cell lines from 22 distinct lineages found AR-positive prostate cancer cells to be amongst the most sensitive models (**Figure 6C**). This screen also revealed multiple myeloma and neuroblastoma cell lines, which show acute dependence on p300/CBP^39,45^, to be markedly sensitive to CBPD-409 relative to the majority of tested normal and cancer cell lines (**Figure 6C**). Overall, we find p300/CBP degradation using CBPD-409 to be a potent therapeutic strategy with high selectivity to prevent cell growth of AR-positive prostate cancer.

CBPD-409 was specifically developed to possess high oral bioavailability (50%) and excellent pharmacokinetic properties^35^ to support its translational potential, which distinguishes it from the previously reported p300/CBP PROTACs^38,39^. In an initial *in vivo* efficacy study using VCaP-derived xenograft tumors in intact SCID mice, oral administration with 3 mg/kg of CBPD-409 significantly inhibited tumor growth (**Figures 6D-E and S8A**). Despite potent degradation of mouse p300/CBP orthologs, no signs of toxicity were identified as assessed by body weights, blood pathology, or vital organ functioning (**Figures S8B-G**).

To further evaluate the safety profile of CBPD-409, a thorough toxicity analysis was conducted in immune-competent CD1 mice (**Figure S9A**). Here, we confirmed the potent degradation of p300/CBP in multiple mouse organs (**Figures 7A and S9B**). Despite efficient target degradation, there were no significant changes in the weight of animals or vital organs during CBPD-409 treatment (**Figures 7B and S9C**). Furthermore, histological examination at the endpoint (day 32) revealed no evidence of toxicity in the mouse colon, liver, kidney, spleen, small intestine, mesenteric lymph nodes, or pancreas tissues (**Figure 7C**). Particularly, CBPD-409 did not affect the anatomical and spatial morphology of any of the prostatic lobes, and no evidence of atrophy, hyalinization, fibrosis, or emergence of any neoplastic phenomenon was identified (**Figure 7C**). Blood biochemistry analyses and complete blood cell counts also showed that CBPD-409 did not adversely affect liver and kidney functions or alter blood cell composition (**Figures 7D-E**). Notably, unlike the thrombocytopenia exhibited by p300/CBP BRDi or BET inhibitors^46,47^, CBPD-409 did not affect platelets in CD1 mice (**Figure 7E**), which was further confirmed by bone marrow H&E staining showing that CBPD-409 did not affect megakaryocytes^48^ (**Figure 7F**). Goblet cell depletion, which is another major side effect exhibited by BET inhibitors^49-52^, was furthermore not observed in CBPD-409 treated mice (**Figure 7G**). The only observed side effect was a reversible defect in germ cell maturation and testicular atrophy, which resolved after discontinuing CBPD-409 treatment (**Figure S9D**). We additionally evaluated the toxicity of CBPD-409 in CD rats. In rat tissues, p300 and CBP were efficiently degraded, but no evidence of toxicity in vital organs was observed (**Figures S9E-F**). Consistently, no significant alteration of liver and kidney functions or thrombocytopenia were observed in CBPD-409 treated rats (**Figures 7H-I**). We also tested the cytotoxicity of CBPD-409 in human primary CD3+ pan T cells, human primary NK cells, and immortalized human B cells (GM24694). In contrast to the significant inhibitory effects of ZBC-260 (a BRD4 degrader^53^) on CD3+ pan T cells and B cells, CBPD-409 exhibited no cytotoxicity in the three tested cell types (**Figure 7J**).

**Figure 7:**
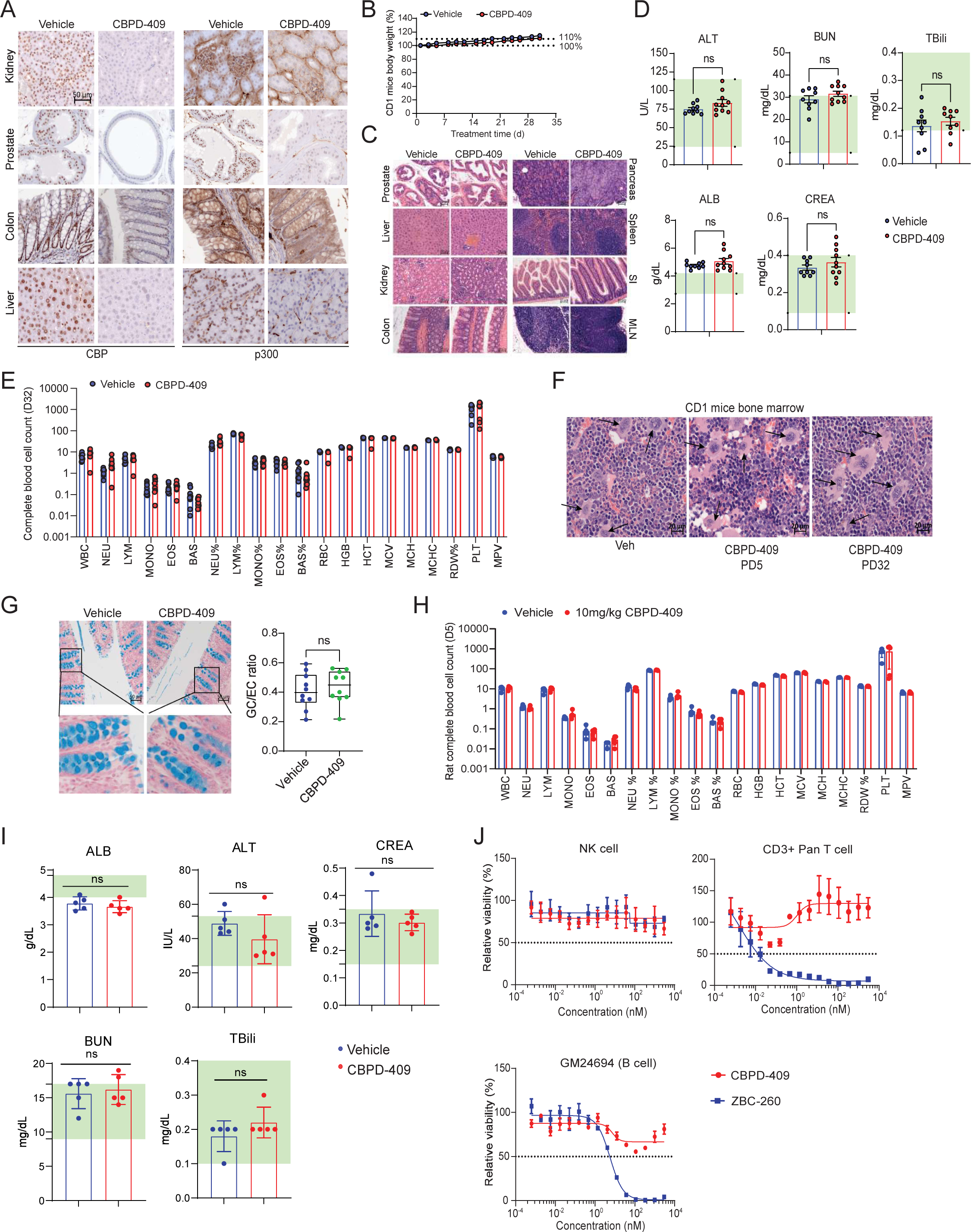
CBPD-409 displays no evident toxicity in murine models and human primary cells. **a.** Representative immunohistochemistry images showing staining of murine CBP and p300 in specified organs of CD-1 IGS mice treated with CBPD-409 (3 mg/kg, administered orally three days per week). n=10 per treatment. Scalebar = 50 µm. **b.** CD1 mice body weight (%) measurements throughout the treatment period from vehicle and CBPD-409 treated groups (two-sided t-test). Data are presented as mean +/− SEM (vehicle: n = 10, CBPD-409: n=10). **c.** Representative H&E staining of major organs from CD1 mouse study. Scalebar = 50 µm. **d.** Liver function and kidney function tests from CD1 mouse study. ALT: alanine transaminase; AST: aspartate transaminase; TBili: total bilirubin; BUN: blood urea nitrogen; CREA: creatinine. Vehicle: n=9; CBPD-409: n=9; two-sided t-test. **e.** Complete blood count from CBPD-409 PD32 CD1 mice. Data are presented as mean +/− SD (n = 8, biological replicates). **f.** Representative H&E staining of femur bone marrow samples from CD1 mouse study. The megakaryocytes are indicated by the arrows. Scalebar = 20 µm **g.** Representative alcian blue staining images from the large intestinal tract harvested from CD1 mice (n = 10/treatment group). Right, quantification of goblet:epithelial cell (GC/EC ratio) densities in the colon (two-sided t-test). Data are presented as mean +/− SEM. **h.** Complete blood count from CBPD-409 PD5 CD rats. CD rats were treated with vehicle or CBPD-409 (10 mg/kg, administered orally three days per week). Data are presented as mean +/− SD (n = 5, biological replicates). **i.** Liver function and kidney function tests from CD rat study. ALB: albumin, ALT: alanine transaminase; CREA: creatinine; BUN: blood urea nitrogen; Tbili: bilirubin. Vehicle: n=5; CBPD-409: n=5; two-sided t-test. **j.** Dose-response curves of human primary CD3+ pan T cells, human primary NK cells, and immortalized human B cells (GM24694) treated with CBPD-409 and BRD4 degrader ZBC-260. Data are presented as mean +/− SD (n = 3).

### p300/CBP degradation inhibits CRPC tumor growth and synergizes with enzalutamide

Next, we utilized a VCaP castration-resistant prostate cancer model (VCaP-CRPC) for further assessment of CBPD-409 anti-tumor efficacy alone or in combination with the AR antagonist enzalutamide (**Figure 8A**). While single agent CBPD-409 significantly suppressed the growth of VCaP-CRPC tumors, it showed dramatically higher anti-tumor potency when combined with enzalutamide, leading to disease regression in over 60% of the animals (11 out of 18) (**Figures 8B-C and S10A**). Consistent with our *in vitro* findings, immunohistochemistry (IHC) staining and western blotting of tumor xenografts five days post treatment confirmed significant downregulation of p300/CBP, AR, PSA, MYC, Ki67, CCND1, NKX3-1, CITED2, H3K27ac, and H2BK20ac (**Figures 8D and S10B-C**).

**Figure 8:**
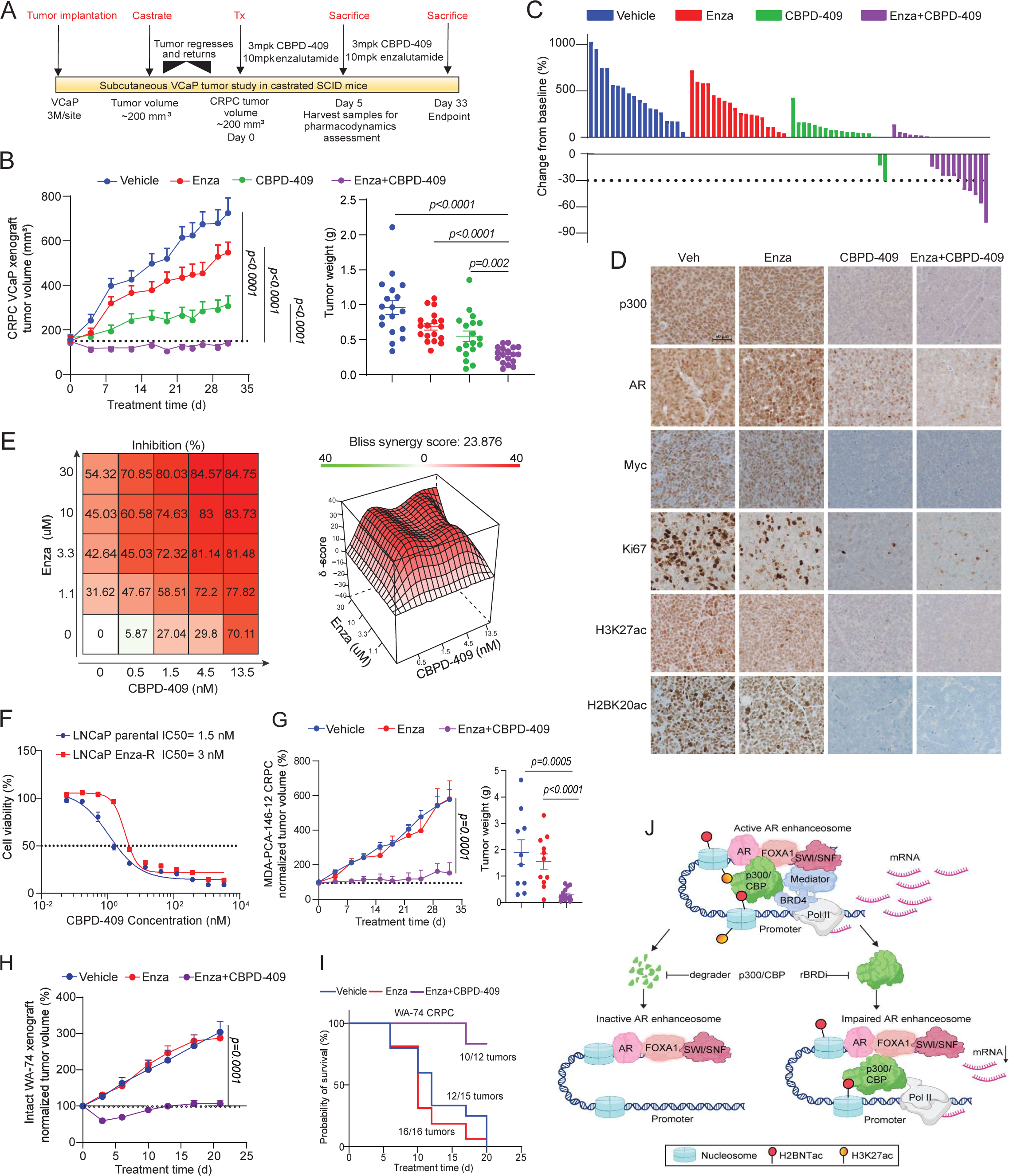
p300/CBP degradation inhibits CRPC tumor growth and synergizes with enzalutamide. **a.** Schematic of the CBPD-409 *in vivo* efficacy study utilizing the VCaP-CRPC xenograft model. VCaP cells were subcutaneously implanted in SCID mice, which underwent castration two weeks post-implantation once the tumors reached 200 mm³ to induce disease regression. Subsequently, castration-resistant tumors re-grew to 200 mm³, at which point the treatments commenced. Mpk, mg/kg. **b.** Graph depicting the tumor volume curves and tumor weights in the VCaP-CRPC model, measured biweekly using calipers. Treatments included vehicle, enzalutamide (Enza, 10 mg/kg, administered orally five days per week), CBPD-409 (3 mg/kg, administered orally three days per week), and CBPD-409 in combination with Enza. Statistical analysis was performed using a two-sided t-test. Data are presented as mean ± SEM. Sample sizes are as follows: vehicle, n = 18; CBPD-409, n = 18; Enza, n = 18; Enza + CBPD-409, n = 18. **c.** Waterfall plot illustrating the change in tumor volume after 33 days of treatment from the VCaP-CRPC study. **d.** Representative immunohistochemistry images from the VCaP-CRPC xenograft study for the indicated protein (n = 4 tumors per treatment). Scalebar = 50 µm. **e.** VCaP cells were treated with noted concentrations of CBPD-409 and/or Enza to evaluate drug synergism using the Bliss Independence method. The average synergy score of CBPD-409 and Enza is 23.876 (>10 indicates drug synergism). Data representation includes the mean of four replicates. **f.** Dose-response curves and IC50 of LNCaP parental and LNCaP enzalutamide-resistant (Enza-R) cells treated with CBPD-409. Data are presented as mean +/− SD (n = 6). **g.** Graph depicting tumor volume curves and tumor weights in the MDA-Pca-146-12 CRPC PDX model, measured biweekly by calipers. Treatments included vehicle, Enza, and combination Enza with CBPD-409. Statistical analysis was performed using a two-sided t-test. Data are presented as mean ± SEM. Sample sizes are as follows: vehicle, n = 10; Enza, n = 12; Enza + CBPD-409, n = 14. **h.** Tumor volume curve in the intact WA-74 PDX model, measured biweekly using calipers. Treatments included vehicle, Enza, and combination of Enza and CBPD-409. Statistical analysis was conducted by two-sided t-test. Data are presented as mean ± SEM. Sample sizes are as follows: vehicle, n = 12; Enza, n = 12; Enza + CBPD-409, n = 14. **i.** Kaplan-Meier survival plot illustrating the survival rates of mice bearing WA-74 CRPC PDX tumors, treated with vehicle, Enza, and combination of Enza and CBPD-409. Survival is measured up to the point where tumor volume reaches 2000 mm³. **j.** Mechanism of action of p300/CBP degrader versus reader bromodomain inhibitor (rBRDi) in disrupting activated AR enhanceosome in prostate cancer cells. Transient degradation of p300/CBP triggers selective loss of histone marks, followed by dislodgement of RNA Pol II. Conversely, inhibition of p300/CBP bromodomain only partially represses histone marks without affecting RNA Pol II loading.

Consistent with the *in vivo* data, combination treatment with CBPD-409 and enzalutamide showed marked synergism in inhibiting the growth of VCaP cells (Bliss synergy score = 23.8; **Figure 8E**). Notably, CBPD-409 retained comparable efficacy between parental and an enzalutamide-resistant LNCaP cell line (**Figure 8F**). To analyze the potential efficacy of CBPD-409 in treating enzalutamide-resistant CRPC *in vivo*, we treated the enzalutamide-resistant patient-derived xenograft (PDX) model, MDA-PCa-146-12^54^, with the combination therapy and observed profound suppression of tumor growth (**Figures 8G and S11A-C**). We further employed an in-house CRPC PDX model, WA-74, that is inherently resistant to enzalutamide. Here, the combined treatment of CBPD-409 and enzalutamide significantly inhibited the growth of WA-74 tumors (**Figure 8H**). In the more aggressive castration-resistant derivative of the WA-74 PDX tumors (labeled as WA74-CRPC), combination CBPD-409 and enzalutamide treatment markedly suppressed tumor growth and enhanced survival of tumor-bearing mice (**Figures 8I and S11D-E**). Taken together, our *in vitro* and *in vivo* efficacy data demonstrate that targeting p300/CBP proteins for degradation using CBPD-409 represents a promising therapeutic strategy for advanced AR-dependent prostate cancers, while exhibiting minimal on or off-target toxicity.

## Discussion

The ‘histone code’ forms the basis of epigenetics, positing that the activity of DNA is intricately regulated by post-translational modifications of histone proteins that are commonly referred to as histone marks^55-57^. These chemical modifications are believed to lead to various chromatin states, each with distinct functional implications for gene expression and cellular fate^52^. For instance, at *cis*-regulatory enhancer elements, the presence of H3K27ac on the flanking nucleosomes is associated with higher transcriptional activity^26^. More recently, several acetylation marks on the N-terminus of H2B were also shown to demarcate active enhancers and to be exclusively catalyzed by the p300/CBP enzymes^27^. In this study, screening a set of transcriptional related histone marks in prostatectomy specimens, we uncovered H2BNTac to be specifically and consistently elevated in prostate cancer lesions compared to the adjacent benign epithelia. Consistently, in CRISPR-based dependency maps (DepMap), p300 ranks as the most essential histone acetyltransferase in AR-driven prostate cancer cells, implying a pivotal role of p300/H2BNTac on prostate cancer cell viability.

Notably, about half of the AR enhancers were co-occupied by p300 in prostate cancer cells, which was deterministic of stronger transcriptional activation evidenced both by higher accessibility and subsequent recruitment of the mediator and RNA polymerase II complexes. The genome-wide distribution profiles of H2BK5ac and H2BK20ac in prostate cancer cells further confirmed a specific enrichment of H2BNTac at AR/p300 co-bound enhancers. Given the recent evidence that depletion of H3K27ac does not affect enhancer-associated gene expression^58-60^, our data suggests that the p300/CBP-catalyzed H2BNTac marks could play a more critical role in mediating its enhancer-associated transcriptional duties. In this context, one could also speculate elevated H2BNTac to be a predictive biomarker of enhancer-driven tumors and, thus, useful in selecting and monitoring patients on p300/CBP or other enhancer-targeted therapeutics.

Current p300/CBP inhibitors target various domains of p300/CBP, each with distinct mechanisms of action. For instance, reader bromodomain inhibitors target the bromodomain to impair chromatin reader functions^17,21^, HAT inhibitors target the histone acetyltransferase domain to reduce catalytic activities^20^, and Kix domain inhibitors disrupt protein-protein interactions^61,62^. Amongst these candidates, reader bromodomain inhibitors are known for their high specificity and drug-like properties^63^, with CCS1477 being tested in clinical trials^17^. However, in this study, we demonstrate that p300/CBP deposit H2BNTac on nucleosomes without requiring the activity of their reader bromodomain. This is consistent with prior reports that bromodomain inhibition does not dislodge p300 from the chromatin^23,64^ and, as we show, fails to extinguish its oncogenic gene programs in prostate cancer cells. Thus, our findings make a compelling mechanism-based case for the development of p300/CBP PROTACs. As proof-of-concept, using related bromodomain inhibitor and PROTAC compounds (i.e., GNE-049 and CBPD-409), we demonstrate only p300/CBP degradation to completely extinguish its catalytic activity and trigger a potent growth inhibitory effect in several preclinical prostate cancer models. Compared to other p300/CBP degraders^38,39,65^, including the CBPD-268 compound published by our group^65^, CBPD-409 is the only compound with good oral bioavailability and high target selectivity^35^, and CBPD-409 possesses better target degradation kinetics. CBPD-268 also has unintended activity against BRD family proteins^65^. Moreover, unlike the p300/CBP PROTAC targeting the HAT domain (JQAD1)^39^, bromodomain-binding p300/CBP PROTACs maintain the specificity and binding affinity of reader bromodomain inhibitors, thus also ensuring their strong on-target effects and safety.

Our group previously developed a highly potent STAT3 PROTAC degrader with excellent tumor inhibitory effects in leukemia and lymphoma cell lines, supporting the notion that PROTAC degraders have the potential to be developed for clinical use^67^. However, both on-target and off-target toxicities of PROTAC degraders have impeded their clinical testing and development. Here, using global proteomics, we demonstrate CBPD-409 to be highly selective for the p300 and CBP proteins, with no change detected even in the abundance of other bromodomain-containing proteins. Further, despite the degradation of p300/CBP in mouse tissues, we saw no dose limiting toxicities in any of the animal studies involving prolonged treatments with CBPD-409. This notably includes extensive toxicology studies for CBPD-409 carried out in immune-competent CD1 mice and rats, as well as cytotoxicity assessments in primary human cells. For CBPD-409, we also found no evidence of goblet and platelet cell toxicities that are associated with BRD4 inhibitors/degraders^49-52^ or p300/CBP bromodomain inhibitors^46^. We did, however, detect reversible inhibition of spermatogenesis in the mouse testis upon treatment with CBPD-409. Thus, our studies have established the groundwork for IND-enabling studies to be conducted with CBPD-409 and its related analogues and warrants safety and efficacy assessments of p300/CBP PROTACs in higher-order primates and, eventually, in early phase human clinical trials.

In summary, we report a striking elevation of the p300/CBP-catalyzed H2BNTac histone marks in prostate cancer lesions. p300 co-binding identifies the highly active AR enhancer circuitry in malignant cells, where p300/CBP deposit H2BNTac at the flanking nucleosomes in a bromodomain-independent manner. Accordingly, bromodomain inhibitors neither effectively deplete H2BNTac nor fully inhibit p300/CBP-dependent transcriptional programs in prostate cancer cells. As an alternative therapeutic strategy, using a p300/CBP bromodomain-binding warhead, we develop CBPD-409 as a PROTAC degrader that has excellent oral bioavailability and pharmacokinetic properties. In a large panel comprising 131 human-derived normal and cancer cell lines, CBPD-409 shows preferential cytotoxicity in AR-driven prostate cancer versus normal prostate and other cancer models. Mechanistically, p300/CBP degradation triggers a rapid and complete loss of the p300/CBP-dependent histone acetylome at AR enhancers and disrupts the recruitment of coactivators (namely MED1 and Pol II) that constitute a functional enhanceosome complex (**Figure 8J**). To our knowledge, this is the first study that provides a mechanistic rationale for the development of p300/CBP degraders instead of the bromodomain inhibitors, which has direct implications for ongoing pharmacological and clinical efforts focused on targeting these enzymes in human cancers. Given the tolerability and more potent anti-tumor effects of CBPD-409, the recently developed p300/CBP PROTACs^35,65,68^ could be powerful pharmacologic agents to treat enhancer-addicted cancers.

## Methods

### Cell lines and compounds

All cell lines were originally obtained from ATCC, DSMZ, ECACC, Lonza, or internal stock. CWR-R1 and LNCaP parental/enzalutamide-resistant (LNCaP-EnzR) cells were gifts from D. Vander Griend (University of Illinois at Chicago). All cells were genotyped every 6 months to ensure their identity at the University of Michigan Sequencing Core and tested every 2 weeks for Mycoplasma contamination. Gibco RPMI-1640 + 10% FBS (ThermoFisher) were used for LNCaP, 22RV-1, CWR-R1, PC-3, and DU145 cells. VCaP was grown in Gibco DMEM Glutamax + 10% FBS (ThermoFisher). CBPD-409 and CBPD-409-Me were synthesized in Dr. Shaomeng Wang’s lab (see **Supplementary Methods**). GNE-049, CCS1477, A485, JQAD1, enzalutamide, carfilzomib, and thalidomide were purchased from Selleck Chemicals. dCBP1 was purchased from MedChemExpress.

### Antibodies

For immunoblotting, the following antibodies were used: p300 (Invitrogen: MA1-16608); CBP (Invitrogen: PA5-27369); AR (Abcam: ab133273); H2BK5ac (Cell Signaling Technology: 12799S); H2BK20ac (Cell Signaling Technology: 34156S); H2BK12ac (Abcam: ab40883); H2BK16ac (Abcam: ab177427); H3K27ac (Cell Signaling Technology: 8173S); H3K18ac (Active Motif: 39755); H3K4me1 (Abcam: ab8895); H3K4me3 (Active Motif: 39060); H3K27me3(Millipore: 07-449); H2B (Active Motif: 39210); H3 (Cell Signaling Technology: 3638S); H2BK120ac (Active Motif: 39119); Myc (Cell Signaling Technology: 9402S); KLK3/PSA (Dako: A0562); CITED2 (Abcam: ab108345); NKX3-1 (Cell Signaling Technology:83700S); CCND1 (Abcam: ab16663); FOXA1 (Thermo Fisher Scientific: PA5-27157); Vinculin (Cell Signaling Technology: 18799S); GAPDH (Santa Cruz Biotechnology: sc-47724); BRD2 (Bethyl Laboratories: A700-008); BRD3 (Bethyl Laboratories: A302-368A); BRD4 (Bethyl Laboratories: A700-004CF); GSPT1 (Proteintech: 28130-1-ap); Aiolos (Cell Signaling Technology: 15103S); Ikaros (Cell Signaling Technology: 14859S).

For ChIP-seq, the following antibodies were used: H3K27ac (Diagenode: C15410196); p300 (Abcam: ab14984); H2BK20ac (Cell Signaling Technology: 34156S); H2BK5ac (Cell Signaling Technology: 12799S); AR (Millipore: 06-680); FOXA1 (Thermo Fisher Scientific: PA5-27157); BRD4 (Diagenode: C15410337); H3K18ac (Active Motif: 39755); MED1 (Active motif: 61065); ERG (Cell Signaling Technology: 97249S); RNA Pol II (Active motif: 39097); H3K18ac (Active Motif: 39755).

For immunohistology and immunofluorescent staining, the following antibodies were used: p300 (Invitrogen: 33-7600); CBP (Invitrogen: PA5-27369); AR (Abcam: ab133273); H3K27ac (Abcam: ab4729); Ki67 (Ventana Medical Systems: 790-4286); H2BK20ac (Abcam: ab177430); H2BK5ac (Abcam: ab40886); CCND1 (Cell Marque: 241R-18).

### Cell viability assay

A total of 4000 cells per well were plated into poly-D lysine coated 96-well plates (Corning) in culture medium and incubated at 37 °C with 5% CO_2_. 24 hours after seeding, the cells were treated with a serial dilution of compounds with 6 replicates of each concentration. The cells were then incubated for 120 hours, and the CellTiter-Glo assay (Promega) was performed based on the manufacturer’s instruction to determine cell survival rate. The luminescence signal from each well was read by the Infinite M1000 Pro plate reader (Tecan). The results were analyzed by GraphPad Prism software (GraphPad Software).

### Incucyte proliferation assays

A total of 4000 cells per well were seeded in clear flat bottom poly-D lysine coated 96-well plates. After 24 hours incubation, different concentrations of compounds were added to the cells. Every 4 hours, phase object confluence (percentage area) for proliferation was measured.

### Immunoblotting

Cells were lysed by RIPA buffers (ThermoFisher) with 1x Halt™ Protease Inhibitor Cocktail (ThermoFisher) and denatured in the complete NuPage 1× LDS/reducing agent buffer (Invitrogen) with 15-minute incubation at 70 °C. The protein concentration was measured by Pierce BCA Protein Assay Kit (ThermoFisher). 15-30 µg protein was loaded and separated on NuPAGE 3 to 8%, Tris-Acetate Protein Gel (ThermoFisher) or NuPAGE 4 to 12%, Bis-Tris Protein Gel (ThermoFisher) and transferred to 0.45-μm nitrocellulose membranes (ThermoFisher). The membranes were blocked with blocking buffer (Tris-buffered saline, 0.1% Tween (TBS-T), 5% non-fat dry milk) for 1 hour and then blotted with primary antibodies in 4°C overnight. After incubation with HRP-conjugated secondary antibodies, membranes were imaged on an Odyssey CLx Imager (LiCOR Biosciences).

### RNA extraction and quantitative polymerase chain reaction

Total RNA was extracted from cells using QIAzol Lysis Reagent (QIAGEN). Initially, 1x10^6 cells were lysed with 700 µL of QIAzol. The lysates were incubated at room temperature for 5 minutes to dissociate nucleoproteins, then mixed with 140 µL chloroform and centrifuged at 12,000 g for 15 minutes at 4°C to achieve phase separation. The aqueous phase was then mixed with 1.5x volume of ethanol and applied to a RNeasy Mini spin column (QIAGEN) for RNA purification, following the miRNeasy Mini Kit protocol. The RNA was then eluted in RNase-free water, and its concentration and purity assessed using a NanoDrop spectrophotometer.

For qPCR analysis, cDNA was synthesized using the extracted RNA as a template. Reverse transcription was carried out using Maxima First Strand cDNA Synthesis Kit (ThermoFisher), following the manufacturer’s instructions. The resulting cDNA was then used for quantitative PCR (qPCR) using SYBR™ Green PCR Master Mix (Applied Biosystems), depending on the specific gene targets. The qPCR reactions were performed in QuantStudio 5 Real-Time PCR system (Applied Biosystems), and the data were analyzed using the ΔΔCt method to quantify gene expression levels, normalizing to the expression of the *GAPDH* gene.

Primers used in this study are listed below: *GAPDH* forward (F): ACAACTTTGGTATCGTGGAAGG and reverse (R): GCCATCACGCCACAGTTTC; *KLK3*/PSA F: GACCAAGTTCATGCTGTGTGC and R: CCACTCACCTTTCCCCTCAAG; *KLK2* F: TCAGAGCCTGCCAAGATCAC and R: CACAAGTGTCTTTACCACCTGT; *TMPRSS2* F: GTCCCCACTGTCTACGAGGT and R: CAGACGACGGGGTTGGAAG; *CITED2* F: CCTAATGGGCGAGCACATACA and R: GGGGTAGGGGTGATGGTTGA; *NKX3-1* F: CCCACACTCAGGTGATCGAG and R: GAGCTGCTTTCGCTTAGTCTT; *MYC* F: CGGAAGGACTATCCTGCTGC and R: CAAGACGTTGTGTGTTCGCC; *CCND1* F: GCTGCGAAGTGGAAACCATC and R: CCTCCTTCTGCACACATTTGAA.

### CRISPR knockout

Short guide RNAs (sgRNAs) targeting the exons of human p300 or CBP were designed by Benchling (https://www.benchling.com/). Non-targeting control sgRNA, p300, and CBP sgRNAs were cloned into lentiCRISPR v2 plasmid based on a previous report^69^. LNCaP and 22Rv1 cells were transiently transfected with lentiCRISPR v2 encoding control or pair of two independent p300 or CBP-targeting sgRNAs. Forty-eight hours after transfection, cells were selected with 1 μg ml−1 puromycin for three days. Western blot was performed to identify the knockout efficiency.

sgRNA sequences used are as follows: sgNC#1:5′-GTAGCGAACGTGTCCGGCGT-3′; sgNC#2:5′-GACCGGAACGATCTCGCGTA-3′; sgp300#1: 5’-CACCGTTCAATTGGAGCAGGCCGA-3’; sgp300#2: 5’-CACCGCATCCCTGTGTTCATTCCCA-3’; sgCBP#1: 5’-CACCGCGCGTGACCAGTCATTTGCG-3’; sgCBP#2: 5’-CACCGCAACTGTCGGAGCTTCTACG-3’.

### siRNA-mediated gene knockdown

The human non-targeting control (Cat#: D-001810-10-05), p300 (Cat#: L-003486-00-0005), and CBP (Cat#: L-003477-00-0005) ON-TARGETplus SMARTPool siRNAs were ordered from Horizon Discovery. Cells were plated in a 6-well plate at the density of 300,000 cells per well. After 24 hours, cells were transfected with 30 nM of siRNAs using the RNAiMAX transfection reagents (Life Technologies) on two consecutive days. The protein was extracted on day 3 to identify efficient (>80%) knockdown of the target genes.

### Acetyl (lysine)-proteomics analysis

#### 1. Chemicals and Instrumentation

DL-dithiothreitol (DTT), iodoacetamide (IAA), trifluoroacetic Acid (TFA), acetonitrile (ACN), and methanol were purchased from Sigma. Trypsin was purchased from Promega. C18 SPE Cartridge was purchased from Waters (Milford, MA, USA). Anti acetyl-Lysine beads were purchased from Cell Signaling Technology. Ultrapure water was prepared from a Millipore purification system. An Ultimate 3000 nano UHPLC system coupled with a Q Exactive HF mass spectrometer (Thermo Fisher Scientific) with an ESI nanospray source was used.

#### 2. Protein Extraction

Samples were taken from -80°C and homogenized in 1 mL lysis buffer (8 M urea, 100 mM Tris, pH 8.0, 1% protease inhibitor) by sonication. Any cell debris were removed by centrifugation at 15,000 rpm for 15 min at 4°C. The protein concentration was then determined by using the BCA assay.

#### 3. Protein Digestion

2 mg protein of each sample was diluted to 1 mL with lysis buffer. Disulfide bridges were reduced by 4.5 mM DTT at 37°C for 1 h. Reduced cysteine residues were alkylated by 10 mM iodoacetamide (IAA) in the dark at room temperature for 30 min. The solution was then diluted to 4 mL with 50 mM ammonium bicarbonate and subjected to overnight digestion at 37°C with trypsin (Promega) using an enzyme to substrate ratio of 1:200 (w/w). After digestion, TFA was added to 1% final concentration. Any precipitate was removed by centrifugation at 1,780 g for 15 min.

#### 4. Peptide Desalting

Purification of peptides was performed at room temperature on C18 reversed-phase columns. The columns were first conditioned by 100% ACN followed by 0.1% TFA, 50% ACN and then equalized by 0.1% TFA. The acidified and cleared digests were then loaded onto columns. After a wash by 0.1% TFA, the peptides were eluted from the column by 0.1% TFA, 50% ACN.

#### 5. Acetylated Peptide Enrichment

The eluent was lyophilized to dry and resuspended in IAP buffer (50 mM NaCl, 10 mM Na_2_HPO_4_, 50 mM MOPS, pH 7.2). Before incubation, the beads were washed in pre-chilled PBS 4 times. The peptide solution was then added into the vial containing motif antibody beads and incubated on a rotator for 4 hours at 4°C. After incubation, the beads were washed with pre-chilled IAP buffer 3 times and HPLC grade water 4 times. The enriched peptides were eluted from the beads by 0.1% TFA.

#### 6. Nano LC-MS/MS Analysis

##### 6.1 NanoLC

Nanoflow UPLC: Ultimate 3000 nano UHPLC system (ThermoFisher Scientific)

Nanocolumn: trapping column (PepMap C18, 100Å, 100 μm × 2 cm, 5 μm) and an analytical column (PepMap C18, 100Å, 75 μm × 50 cm, 2 μm)

Loaded sample volume: 1 μg

Mobile phase: A: 0.1% formic acid in water; B: 0.1% formic acid in 80% acetonitrile

Total flow rate: 250 nL/min

LC linear gradient: from 2 to 8% buffer B in 5 min, from 8% to 40% buffer B in 120 min, then from 40% to 90% buffer B in 5 min

##### 6.2 Mass Spectrometry

The full scan was performed between 300-1,650 m/z at the resolution 60,000 at 200 m/z. The automatic gain control target for the full scan was set to 3e6. The MS/MS scan was operated in Top 20 mode using the following settings: resolution 15,000 at 200 m/z; automatic gain control target 1e5; maximum injection time 19ms; normalized collision energy at 28%; isolation window of 1.4 Th; charge sate exclusion: unassigned, 1, > 6; dynamic exclusion 30 s.

#### 7. Data Analysis

Raw MS files were analyzed and searched against the Homo sapiens protein database based on the species of the samples using Maxquant (2.3.0.0). The parameters were set as follows: the protein modifications were Carbamidomethylation (C), Oxidation (M) (variables), Acetyl (K) (variables), Acetyl (Protein N-term) (variables), Acetyl (N-term) (variables); the enzyme specificity was set to trypsin; the maximum missed cleavages were set to 5; the precursor ion mass tolerance was set to 10 ppm, and MS/MS tolerance was 0.5 Da.

### RNA-seq and data analysis

RNA extraction was performed as previously described^4^. Following extraction, ribosomal RNA (rRNA) was depleted from the total RNA samples using the RiboErase module of the KAPA RNA Hyper+RiboErase HMR Kit (Roche Cat. No. 08098140702). The rRNA-depleted RNA was then applied for library preparation. The procedure was conducted following the protocol provided with the KAPA RNA Hyper+RiboErase HMR Kit. The prepared libraries were validated for quality and quantification using the Agilent 2100 Bioanalyzer. The libraries were then sequenced on an Illumina NovaSeq 6000, utilizing a paired-end sequencing strategy (2 × 150 nucleotide read length with sequence depth of 15–20M paired reads).

RNA-seq data was handled via Kallisto (version 0.46.1)^70^. R package EdgeR (edgeR_3.39.6) was used to generate normalized and filtered read counts (counts >10)^71^. Differential expression was performed using Limma-Voom (limma_3.53.10)^72^. R package fgsea (fgsea_1.24.0) was used for Gene Set Enrichment Analyis (GSEA). Further R packages tidyverse, gtable, gplots, ggplot2 and EnhancedVolcano (EnhancedVolcano_1.15.0) were also used for generating figures.

### ChIP-seq and data analysis

Chromatin immunoprecipitation experiments were conducted using the Ideal ChIP-seq Kit for Transcription Factors (for AR, FOXA1, p300, ERG, BRD4, and MED1 ChIP-seq) or Histones (for H3K27ac, H2BK5ac, H3K20ac, and RNA Pol II ChIP-seq) (Diagenode) following the manufacturer’s protocol. Briefly, 4 × 10^6^ cells (for transcription factors) and 1x10^6^ cells (for histones) for each ChIP reaction were applied for cross-linking for 10 minutes in 1% formaldehyde solution, followed by 1/10 volume 1.25 M glycine for 5 minutes at room temperature to quench the formaldehyde. The chromatin was extracted by cell lysis and sonication (Bioruptor, Diagenode) to break down chromatin into the size of 200-600 bp. Sheared chromatin was then used for immunoprecipitation with the individual antibody (8 μg for transcription factor or 1 μg for histone), with overnight incubation at 4 °C. DNA fragments were de-crosslinked and purified. Purified DNA was then subjected for sequencing following the manufacturer’s instructions (Illumina). The ChIP-seq libraries were prepared from purified ChIP samples (1–10 ng) as described previously^4^. Libraries were quantified and quality checked using the Bioanalyzer 2100 (Agilent) and sequenced on the Illumina NovaSeq 6000 Sequencer (2x150-nucleotide read length with sequence depth of 25-35M paired reads).

ChIP-seq data analysis started with trimming using Trimmomatic version 0.39 (settings TruSeq3-PE-2.fa:2:30:10, minlen 50)^73^. BWA was used to align reads to hg38 (GRCh38) human genome reference (“bwa mem” command with options -5SP -T0, version 0.7.17-r1198-dirty)^74^. Alignments were filtered using samtools (quality score cutoff of 20) and picard MarkDuplicates (removed duplicates)^75,76^. MACS2 was used for peak calling with narrowpeak setting for narrow peaks and a second set of parameters for histone peaks (eg H3K27Ac, --broad -B --cutoff-analysis --broad-cutoff 0.05 --max-gap 500)^77^. In addition, bedtools was used to remove blacklisted regions of the genome from the peaks list (Encode’s exclusion list ENCFF356LFX.bed)^78,79^. UCSC’s tool wigtoBigwig was used for conversion to bigwig formats^80^.

### ATAC-seq and data analysis

ATAC-seq was conducted as previously described^4^. Briefly, 50,000 cancer cells were lysed using CER-I cytoplasmic lysis buffer from the NE-PER kit (Invitrogen) and incubated for 5 minutes on ice with occasional gentle mixing. After centrifugation at 1,300g for 5 minutes at 4°C, the nuclear pellets were isolated. The nuclei were then treated with 50 μl of 1× TD buffer and 2 μl Tn5 enzyme for 30 minutes at 37°C, using the Nextera DNA Library Preparation Kit. After transposition, samples were immediately purified using a Qiagen minElute column and subjected to PCR amplification with NEBNext High-Fidelity 2X PCR Master Mix (NEB). Optimal PCR cycles were determined via qPCR to avoid over-amplification. The amplified libraries were further purified using a Qiagen minElute column and SPRI beads (Beckman Coulter). Libraries were quantified and quality checked using the Bioanalyzer 2100 (Agilent). Finally, the ATAC-seq libraries were sequenced on the Illumina HiSeq 2500 platform, utilizing a 2x50-nucleotide paired-end read length with sequence depth of 30-35M.

Sequencing of ATAC-seq libraries generated fastq files, which were initially processed using Trimmomatic (version 0.39) for trimming^73^. These files were then aligned to the GRCh38/hg38 human genome using bwa mem (version 0.7.17-r1198-dirty)^74^, and the alignments were converted to binary format with SAMtools (version 1.9)^75^. We next eliminated reads from mitochondrial DNA and duplicated reads, using SAMtools and PICARD MarkDuplicates (version 2.26.0-1-gbaf4d27-SNAPSHOT)^75,76^. Peaks in the ATAC-seq data were identified using MACS2 (version 2.1.1.20160309)^77^. Finally, conversion of data to bigwig format was accomplished using the UCSC tool wigtoBigwig^80^.

### Nascent RNA-seq data analysis

Nascent RNA sequencing was conducted using the Click-iT™ Nascent RNA Capture Kit (Invitrogen). Cells were treated with or without CBPD-409 for 2 and 4 hours, with 0.5 mM 5-Ethynyl Uridine (EU) added to the medium 45 minutes before cell harvest. After treatment, cells were washed with PBS, harvested, and lysed with the kit’s lysis buffer. RNA isolation was performed using QIAzol reagent and RNeasy Mini spin column (Qiagen), following previously described methods^4^. For ribosomal RNA depletion, 5 μg of total RNA were processed using the Ribominus™ Eukaryote System v2 (ThermoFisher). The rRNA-depleted, EU-labeled RNA (250-500 ng) was then incubated with 0.25 mM biotin azide in the Click-iT® reaction cocktail for 30 minutes with gentle vortexing. Biotin-conjugated EU-RNA was precipitated using UltraPure™ Glycogen (1 μL, ThermoFisher), 7.5 M ammonium acetate (50 μL, Sigma-Aldrich), and chilled 100% ethanol (700 μL) at –70°C overnight. The sample was then centrifuged at 13,000 × g for 20 minutes at 4°C, and the RNA pellet was washed with 75% ethanol. Finally, 100-200 ng of the purified EU-RNA was used for library preparation and sequencing, as detailed in the RNA-seq section.

The sequencing reads were mapped to the reference genome (GRCh38.p14) downloaded from Gencode^81^ using Burrows-Wheeler Alignment Tool (bwa mem) with default parameters^74^. The resulting sam files were then converted to bam format and sorted using Samtools^75^; bigwig files were then generated using bamCoverage of deeptools^82^ with RPKM normalization method. Quantification of gene and enhancer expression levels was conducted with FeatureCounts^83^. AR binding sites at non-promoter regions were defined as AR enhancers. The coordinates of AR enhancers were based on ChIP-seq peaks of AR, and peaks that overlap with gene promoters (+/- 3 kb of transcription start site (TSS)) were excluded. Normalized enhancer expression levels in TPM were used for data visualization. Meta profile plots were generated with the plotProfile function of deeptools^82^, in which the inputs were merged bigwig files from duplicate libraries using UCSC BigwigMerge tool^80^.

### Enrichment heatmaps and profile plots

The read density heatmaps and enrichment plots were all created using the software Deeptools. The referencePoint parameter was set to +/- 2.5 kb for histone signals and +/- 1 kb for other signals. Other settings included using ‘skipzeros’, ‘averagetype mean’, and ‘plotype se’. The Encode blacklist ENCFF356LFX was used. Non-promoter regions were selected based on annotation by R package ChIPseeker^84^ with +/- 1kb windows to gene regions.

### Peak annotations and overlaps

The R package ChIPpeakAnno^85^ was used to compare samples’ peak lists from MACS^77^. Peaks were reduced within 500 bp, and overlaps were calculated using settings maxgap=-1L, minoverlap=0L, ignore.strand=TRUE, connectedPeaks=c(’keepAll’, ‘min’, ‘merge’). An additional R package, ChIPseeker^84^, was used for comparisons of the enrichment sites to the known gene database (TxDb.Hsapiens. UCSC.hg38.knownGene) with a +/- 1kb relative distance from gene regions.

### Analysis of AR binding sites from human CRPC and benign prostate tissues

Additional published datasets were used from Pomerantz et al^5,6^ and processed using our ChIP-seq pipelines. The Picard utility MergeSamFiles was utilized to combine the aligned bam files, and samtools view -bs was used to subsample the combined bam file to a depth of approximately 100M reads for AR datasets. Peak calling was then repeated using macs2 callpeak, and output bedgraphs were converted using wigToBigWig.

### Super-enhancer analysis

Regions were selected from the overlap comparison of ChIP-seq peaks as peaks only in the AR sample, or peaks found both in the AR sample and the p300 sample. The AR sample aligned bam file was subset to regions from bed files. Super-enhancers were called using HOMER^86^. The subset alignment files were then converted to tag directories using makeTagDirectory command, and peaks were called with findPeaks (options -style super -superSlope 1000 -typical). Line with slope for y=1 cutoff was used to classify the results as super-enhancers and plotted in R (ggplot).

### ChIP-seq peaks quartile categorization and heatmaps

Quartile comparisons were made by taking results from MACS peak calling into R as granges objects and labeled into quartile bins by MACS score. Overlap analysis was performed by IRanges’s subsetbyoverlaps^87^. After some table manipulation and calculations for percentile in bash, data was reloaded into R and plotted as barplots with ggplot.

### Immunohistochemistry (IHC) and immunofluorescence (IF)

IHC and IF were carried out on 4-micron formalin-fixed, paraffin-embedded (FFPE) tissue slices on the Ventana ULTRA automated slide stainer platform. The antigen retrieval was done by heating tissues for cell conditioning media 1 (CC1) and 2 (CC2) with primary antibody incubation done. Anti-rabbit or anti-mouse secondary antibodies, wherever applicable, were used to develop the immune complex. For the IHC, the OmniMap and UltraMap Universal DAB RTU detection kits were used, while for IF, Ventana FITC and Red 610 RTU detection kits were used to develop the signal. For double-IF, H2BK5ac, H2BK20ac, and KRT8 were employed. H2BK5ac/CK8 double IF was carried out using CC1 95°C for antigen retrieval, followed by first antibody (H2BK5ac) incubation, OmniMap anti-rabbit horseradish peroxidase (HRP), and signal development using Discovery 610 Kit (RTU, details below). The second antibody staining was performed consecutively with heat denaturation before the second primary antibody (KRT8) incubation, OmniMap anti-rabbit HRP kit, and signal development using Discovery FITC (RTU, detail below). Similarly, H2BK20ac was used as the first primary antibody to develop Red 610, and KRT8 was used as second primary antibody to develop FITC. An additional step of counterstaining by DAPI kit was used. The essential reagents and their catalog numbers are listed below.

| **Reagent** | **Source** | **Identifier** |
| --- | --- | --- |
| Discovery CC1 | Roche-Ventana Medical System | Catalog No. 06414575001 |
| Discovery CC2 | Roche-Ventana Medical System | Catalog No. 950-223 |
| Discovery Inhibitor | Roche-Ventana Medical System | Catalog No. 760-4840 |
| Discovery OmniMap anti-rabbit HRP | Roche-Ventana Medical System | Catalog No. 760-4311 |
| Discovery OmniMap anti-mouse HRP | Roche-Ventana Medical System | Catalog No. 760-4310 |
| UltraView Universal DAB Detection Kit | Roche-Ventana Medical System | Catalog No. 760-500 |
| Discovery FITC Kit (RTU) | Roche-Ventana Medical System | Catalog No. 760-232 |
| Discovery Red 610 Kit (RTU) | Roche-Ventana Medical System | Catalog No. 760-245 |
| DISC. QD DAPI RUO 05268826001 | Roche-Ventana Medical System | Catalog No. 760-4196 |

### CBPD-409 and enzalutamide formula for *in vivo* studies

CBPD-409 was freshly dissolved in 100% PEG400 before administration to mice. Enzalutamide was prepared in 1% carboxymethyl cellulose (CMC) with 0.25% Tween-80 and homogenized by sonication. Both CBPD-409 and enzalutamide were administered to mice by oral gavage.

### Human prostate tumor xenograft models

Six-to-eight-week-old male severe combined immunodeficiency (SCID) mice were acquired from the breeding colony at the University of Michigan. They were used to establish subcutaneous tumors on both sides of dorsal flanks. Tumor sizes were measured weekly using digital calipers, applying the formula (π/6) (Length × Width²). After study completion, mice were euthanized, and the tumors were harvested and weighed. The University of Michigan’s International Animal Care and Use Committee (IACUC) approved these procedures.

In the VCaP non-castrated tumor model, mice received a subcutaneous injection of 3 million VCaP cells in a mix of serum-free medium and 50% Matrigel (BD Biosciences). Once tumors grew to about 200 mm³, the mice were grouped randomly and received either CBPD-409 (3 mg/kg) or a control treatment through oral gavage three times weekly for five weeks.

For the VCaP castration-resistant model, a similar injection of VCaP cells was administered. After the tumors became palpable (150-200mm³), mice underwent castration. When tumors regrew to their initial size, mice were treated with either 3 mg/kg CBPD-409 or a control, via oral gavage three times weekly, and additionally with or without 10 mg/kg enzalutamide orally five times weekly for five weeks.

### Prostate patient-derived xenograft models

The patient-derived xenograft (PDX) MDA-PCa-146-12 model was obtained from the University of Texas M.D. Anderson Cancer Center. This AR-positive PDX was developed from a CRPC patient as previously described^54^.

The prostate cancer sample used for WA74 PDX development was collected during a rapid autopsy case as part of the Michigan Legacy Tissue Program (MTLP). For this line, metastases were excised from the intestinal mesentery and immediately placed into cold DMEM. Within 2-3 hours, tumor chunks were implanted into subcutaneous pockets of male NSG mice. Mice were monitored weekly. Out of 5 mice, only one animal showed tumor growth after 10 months. This tumor was harvested and sequentially passaged in both NSG and CB17SCID mice. This WA74 PDX line bears *BRCA2* somatic mutation and *TMPRSS2-ERG* fusion.

For the intact WA74 model, the PDXs were propagated in male SCID mice. This involved surgical implantation of 2 mm^3^ tumor pieces, encapsulated in 100% Matrigel, into the flanks of the mice. After the tumor grew to 200 mm^3^, mice were randomized into different treatment groups. These groups were administered with vehicle, 10 mg/kg enzalutamide (5 times per week), or 3 mg/kg CBPD-409 (3 times per week) with 10 mg/kg enzalutamide by oral gavage for 4 weeks.

For CRPC MDA-PCa-146-12 and CRPC WA-74 models, tumors were established in castrated male SCID mice. When tumor sizes reached about 100 mm^3^, similar randomization and division into treatment groups were conducted. These groups received the same treatments as intact WA-74 tumors for 4-5 weeks.

Throughout all studies, the treatment protocols strictly followed the IACUC guidelines, ensuring that the maximum tumor size did not surpass 2.0 cm in any dimension. Mice with xenografts reaching this threshold were humanely euthanized.

### Drug toxicity analysis in CD1 mice and CD rats

#### Complete blood cell count

Blood samples were collected from mice or rats using the submandibular bleeding method, employing K3 EDTA tubes to prevent coagulation. Each animal was gently restrained, and approximately 100-200 µL of blood was carefully obtained from the cheek pouch, ensuring minimal discomfort to the animal. This method was chosen for its minimal invasiveness and reliability in obtaining sufficient blood volume for analysis. Following collection, the blood samples were immediately mixed with EDTA by gentle inversion to prevent clotting. The complete blood cell count (CBC) was performed on the same day as the blood collection to maintain sample integrity. Samples were analyzed by the University of Michigan *in vivo* animal core (IVAC) using a calibrated hematology analyzer suitable for mouse blood.

#### Serum chemistry analysis

Serum was obtained for biochemical analysis through a standardized collection and separation process. More than 200 µL of blood was drawn into serum separator tubes, and tubes were set aside to allow the blood to clot for over 30 minutes. Following the clotting period, the tubes were centrifuged at 1800-3000g for 10 minutes. The cell-free serum was then extracted from the tubes for analysis at IVAC.

#### Histopathological analysis of organs

Multiple organs including the liver, spleen, kidney, colon, small intestine, mesenteric lymph nodes, pancreas, prostate, and testis were systematically evaluated for histopathological changes as previously described^4^. Two pathologists, blinded to the control and treatment groups, examined the stained sections under a brightfield microscope. They assessed general tissue morphology and architectural coherence across all organs. Detailed analyses were conducted at cellular and sub-cellular levels, focusing on specific aspects unique to each organ. Johnsen scoring was done according to the well-established schema detailed in published literature^88,89^.

#### Alcian blue staining

Alcian blue staining was performed using the Alcian Blue Stain Kit (pH 2.5, Abcam) following the manufacturer’s protocol. Tissue sections on slides were first incubated overnight at 58°C, then deparaffinized with xylene and rehydrated through a series of ethanol solutions (100%, 70%) and water, each for 5 minutes. The slides were then treated with acetic acid solution for 3 minutes, followed by incubation in Alcian blue stain (pH 2.5) for 30 minutes at room temperature. After staining, the slides were rinsed in acetic acid for 1 minute and washed three times with water, 2 minutes each. Nuclear Fast Red was applied as a counterstain for 5 minutes, followed by washing, dehydration in ethanol and xylene, and mounting with EcoMount (Thermo Fisher).

The Alcian blue goblet cell: epithelial ratio (GC:EC) was calculated by recording the number of goblet cells and epithelial cells per colonic crypt. In total, 10 colonic crypts were assessed per mouse sample, and the final average score was rendered.

### Analysis of drug synergism

To evaluate potential synergistic interactions between two pharmacological agents, cells were subjected to escalating concentrations of each drug individually over a period of 5 days. The assessment of cell viability post-treatment was conducted using the CellTiter-Glo Luminescent Cell Viability Assay (Promega). This experimental procedure was replicated across four biological replicates. The resulting data were analyzed to quantify the percentage of inhibition relative to untreated control cells. To determine the presence and extent of drug synergy, the data were processed using the online version of the Synergy Finder tool. This analysis employed the Bliss independence model to interpret interactive effects between the two drugs.

### Human research statement

From the pathology archives, patient tissues from either prostatectomy or biopsy of metastatic prostate cancer lesions acquired at the University of Michigan (U-M) Hospital were used for immunohistochemical assessment of luminal markers and histone marks. The U-M Institutional Review Board approved the acquisition and use of clinical formalin-fixed paraffin embedded specimens from the archives not needing patient consent.

## Data availability

All sequencing data generated in this study have been deposited in the following National Center for Biotechnology Information Gene Expression Omnibus (NCBI GEO) repository: GSE255134.

## Acknowledgements

We thank Lanbo Xiao, Ingrid Apel, Jing Hu, Lisa McMurry, Amanda Miller, Christine Caldwell-Smith, Kenneth Gu, Jocelyn Cai, Xiaoming Wang, Prathibha Gajjala, Shuqing Li, Rohit Mehra, and Javed Siddiqui from the Michigan Center for Translational Pathology at the University of Michigan as well as Venky Basrur and the Rogel Cancer Center Proteomics Shared Resource for providing technical assistance. This work was supported by the following mechanisms: Prostate Cancer Foundation (PCF) 2023 Challenge Award, National Cancer Institute (NCI) Specialized Programs of Research Excellence Grant P50-CA186786, NCI Outstanding Investigator Award R35-CA231996 (A.M.C.), NIH F99/K00 Fellowship (A.P.), Rogel Fellowship (A.P.), and PCF 2021 and 2023 Young Investigator Awards (A.P. and J.L.). A.M.C. is a Howard Hughes Medical Institute Investigator, A. Alfred Taubman Scholar, and American Cancer Society Professor.

## Author contributions

J.L., A.P., Y.Q., S.W., and A.M.C. conceived and designed the studies; J.L. performed all *in vitro* and functional genomics experiments with assistance from A.P., S.E., R.S., and M.W.; Y.Q., J.C.-Y.T., and J.L. performed all animal efficacy and toxicity studies with assistance from T.H., Y. Zheng, and Y.C.; E.Y., A.P., and Y. Zhang carried out all bioinformatics analyses with assistance from J.G., M.S., and S.E.; R.M. and S.M. carried out all histopathological evaluations of drug toxicity and quantified all histology-based data; R.M. performed all immunohistochemistry and immunofluorescence staining; F.S. generated next-generation sequencing libraries, and R.W. and X.C. performed the sequencing; U.V. guided the design of drug efficacy studies; Z.C. and M.W. were involved in the discovery, synthesis, and initial profiling of the CBPD-409 compound; J.L., A.P., S.J.M., and A.M.C. wrote the manuscript and organized the final figures.

## Declaration of interests

S.W. was a co-founder and served as a paid consultant to Oncopia Therapeutics, Inc. S.W. and the University of Michigan owned equity in Oncopia Therapeutics, Inc., which was acquired by Roivant Sciences, Inc., and Proteovant Therapeutics, Inc. S.W. is a paid consultant to Proteovant Therapeutics, Inc. The University of Michigan has received a research contract from Oncopia Therapeutics, Inc., for which S.W. serves as the principal investigator. A.M.C. is a co-founder and serves on the scientific advisory board of LynxDx, Oncopia Therapeutics, Flamingo Therapeutics, Medsyn Pharma, and Esanik Therapeutics. A.M.C. serves as an advisor to Tempus, Aurigene Oncology, Proteovant, and Ascentage. The remaining authors declare no conflicts of interests. The authors have filed patents on: 1. orally active p300/CBP degraders and 2. elevation of H2NTac in prostate cancer

---

**Figure S1: p300 is the critical histone acetyltransferase for H2BNTac and assessment of the active ERG cistrome in prostate cancer.**

**a.** Immunoblot analysis of key histone marks in four pairs of matched prostate cancer (T) and benign adjacent tissues (N). Quantitation and fold change (FC) of the respective histone marks is provided to the right. PTMs, post-translational modifications.

**b.** Representative immunofluorescence images of H2BK5ac (green) and H2BK20ac (green) expression in nuclei of paired prostate cancer and benign adjacent tissues. Magnification: 200x. Scalebar = 50 µm.

**c.** Venn diagrams illustrating overlaps of genome-wide p300, H2BK5ac, H2BK20ac, H3K18ac, and H3K27ac ChIP-seq peaks in VCaP cells.

**d.** Bar charts depicting proportion of FOXA1, SMARCA4, p300, and BRD4 ChIP-seq peaks that map to all AR peaks located in non-promoter regions.

**e.** Bar charts depicting proportion of FOXA1, SMARCA4, p300, and BRD4 peaks that map to the top quartile of ERG ChIP-seq peaks located in non-promoter regions.

**f.** ChIP-seq (ERG, FOXA1, p300, MED1, H3K27ac, and H2BK20ac) and ATAC-seq read-density heatmaps at ERG/p300 co-bound and ERG only binding sites in VCaP cells.

**g.** Immunoblot analysis of p300, CBP, and indicated histone marks in 22Rv1 WT, p300 KO, CBP KO, and p300 KO with siCBP cells.

**Figure S2: Characterization of the on-target degradation effects of the p300/CBP degrader, CBPD-409.**

**a.** ChIP-seq read-density heatmaps of H3K27ac at AR/p300 co-bound sites in VCaP cells with 4 hours of 1 µM GNE-049 or 1 µM CCS1477 treatments.

**b.** Immunoblot analysis of p300 and CBP in LNCaP cells treated with 100 nM CBPD-409 for the indicated durations.

**c.** Immunoblot analysis of p300 and CBP in VCaP cells treated with 10 nM or 100 nM CBPD-409 for 4 hours.

**d.** Immunoblot analysis of p300 and CBP in non-neoplastic prostatic cells treated with 10 nM or 100 nM CBPD-409 for 4 hours.

**e.** Immunoblot analysis of p300 and CBP in murine prostate cancer cells (Myc-Cap and TRAMPC2) treated with 10 nM or 100 nM CBPD-409 for 4 hours.

**f.** TMT mass spectrometry (MS) assay to evaluate effects of CBPD-409 (100 nM, 4 hours) on the proteome of 22Rv1 cells. Data plotted log2 of the fold change (FC) versus DMSO control against -log2 of the p-value per protein from n = 3 independent experiments. All t-tests performed were two-tailed t-tests assuming equal variances. CBP and p300 are highlighted in red.

**g.** Immunoblot analysis of BET proteins (BRD2, BRD3, and BRD4) in VCaP cells treated with 100 nM CBPD-409 for the indicated durations.

**h.** Immunoblot analysis of cereblon (CRBN) neo-substrates (GSPT1 and Ikaros) in LNCaP cells treated with 100 nM CBPD-409 for 24 hours.

**i.** Immunoblot analysis of p300 and CBP in VCaP cells pre-treated with different concentrations of thalidomide (Thali) for 1 hour, then treated with CBPD-409 at the indicated concentrations for 4 hours.

**j.** Immunoblot analysis of p300 and CBP in VCaP cells pre-treated with different concentrations of carfilzomib (Carfil) for 1 hour, then treated with CBPD-409 at the indicated concentrations for 4 hours.

**k.** Schematic of CBPD-409-Me structure (inactive analogue).

**l.** Immunoblot analysis of p300 and CBP in LNCaP cells treated with CBPD-409 and CBPD-409-Me at indicated concentrations for the noted durations.

**Figure S3: CBPD-409 suppresses histone and non-histone protein acetylation in VCaP cells.**

**a.** Immunoblot analysis of H3K27ac in VCaP cells treated with 100 nM CBPD-409 or 1 µM GNE-049 for the noted durations.

**b.** Immunoblot analysis of H2BK5ac, H2BK20ac, and H2BK120ac in VCaP cells treated with 100 nM CBPD-409 for the noted durations.

**c.** Summary table of differentially acetylated lysines (acetylated proteins) from mass spectrometry acetylome analysis in VCaP cells treated with DMSO or CBPD-409 for the indicated time. In this experiment, a total of 739 acetylated lysine residues across 426 acetylated proteins were identified with high confidence (localization probability > 0.75).

**d.** Fold change heatmap of all detected acetylated peptides in VCaP cells treated with or without 100 nM CBPD-409 for the indicated durations. The results were obtained from three replicated experiments for each condition.

**e.** ChIP-seq read-density heatmaps of genome-wide H2BK20ac in VCaP cells with 4 hours of 100 nM CBPD-409 or 1 µM GNE-049 treatments.

**Figure S4: CBPD-409 inhibits histone acetylation at the AR enhanceosome without affecting chromatin accessibility and AR/FOXA1 chromatin distribution.**

**a.** Venn diagrams depicting genome-wide changes of H3K27ac ChIP-seq peaks in VCaP cells treated with 100 nM CBPD-409 or 1 µM GNE-049 for 4 hours.

**b.** ChIP-seq read-density heatmaps of genome-wide H3K27ac in VCaP cells with 4 hours of 100 nM CBPD-409 or 1 µM GNE-049 treatments.

**c.** ChIP-seq read-density heatmaps of H3K27ac within CRPC-specific AR binding sites in VCaP cells after 4 hours of 100 nM CBPD-409 or 1 µM GNE-049 treatments.

**d.** ATAC-seq read-density heatmaps and ChIP-seq read-density heatmaps of AR and FOXA1 peaks at AR *cis*-regulatory elements in VCaP cells with 100 nM CBPD-409 and 1 µM SWI/SNF ATPase degrader AU-15330 treatments for 4 hours.

**e.** Immunoblot analysis of AR and FOXA1 sub-cellular localization in VCaP cells with 6 hours of CBPD-409 (100 nM), GNE-049 (1 µM), and A485 (1 µM) treatments.

**Figure S5: CBPD-409 significantly suppresses AR transactivation.**

**a.** GSEA net enrichment score (NES) plot of significantly altered hallmark pathways in VCaP cells treated with 100 nM CBPD-409 for 24 hours.

**b.** GSEA plots for mTORC1, FOXA1, MYC, and ERG signaling pathways using the fold change rank-ordered gene signature from VCaP cells treated with CBPD-409 for 24 hours.

**c.** qPCR of *KLK3*, *KLK2*, *CCND1*, *NKX3-1*, *TMPRSS2*, and *MYC* expression in LNCaP WT, p300 KO, CBP KO, and p300 KO with siCBP cells.

**d.** Immunoblot analysis of indicated proteins in VCaP cells treated with 100 nM CBPD-409, 100 nM GNE-049, or 100 nM A485 for noted durations.

**e.** ChIP-seq tracks of AR, H3K27ac, H2BK20ac, and Pol II within the *TMPRSS2* gene locus in VCaP cells treated with or without 1 µM GNE-049 for 4 hours.

**f.** Average Pol II ChIP-seq coverage profiles of AR down-regulated genes (AD_DN), AR activated genes (AR_UP), and random genes in VCaP cells treated with 100 nM CBPD-409 or 1 µM GNE-049 for 4 hours.

**g.** Densities of 5-ethynyluridine (EU) labeled nascent RNA-seq reads of classical AR target genes (n=200) and highly expressed random genes (n=200) in VCaP cells treated with or without 100 nM CBPD-409 for 2 and 4 hours. Classical AR target genes were defined as those up-regulated by 1 nM R1881. Highly expressed genes were defined as those with log2(CPM) > 6 from RNA-seq data, from which 200 random genes were selected. The data were derived from two independent samples for each group.

**Figure S6: CBPD-409 targets a unique set of cell cycle-related genes to suppress proliferation.**

**a.** Venn diagrams showing overlaps between genes suppressed by CBPD-409 and those suppressed by GNE-049 or CCS1477 in VCaP cells. Data obtained from RNA-seq with two independent samples for each condition.

**b.** GSEA net enrichment score (NES) plot comparing hallmark pathway alterations in VCaP cells: 100 nM CBPD-409 vs. 1 µM GNE-049 treatment for 24 hours.

**c.** Compute overlaps of ranked top 300 CBPD-409 unique down-regulated genes with gene ontology (GO) and Reactome gene sets.

**d.** qPCR of *KLK3*, *CITED2*, *CCND1*, and *NKX3-1* expression in LNCaP cells treated with CBPD-409, GNE-049, or CCS1477 for 24 hours.

**e.** Immunoblot analysis of NKX3-1, CCND1, and CITED2 in VCaP cells treated with 100 nM CBPD-409 for the noted durations.

**f.** Immunoblot analysis of indicated proteins and histone marks in LNCaP cells treated with 100 nM CBPD-409, 1 µM GNE-049, or 1 µM CCS1477 for 24 hours.

**g.** ChIP-seq tracks of H3K27ac, H2BK20ac, and Pol II within the *NKX3-1* and *CCND1* gene loci in VCaP cells treated with 100 nM CBPD-409 or 1 µM GNE-049 for 4 hours.

**Figure S7: CBPD-409 exhibits superior cytotoxicity in AR-positive prostate cancer cells compared to other p300/CBP inhibitors.**

**a.** Dose-response curves of AR-positive prostate cancer cells treated with CBPD-409. Data are presented as mean +/− SD (n = 6).

**b.** Growth curves of LNCaP and VCaP cells treated with CBPD-409, which were measured by Incucyte Live-Cell Analysis. Data are presented as mean +/− SD (n = 6).

**c.** Dose-response curves of AR-negative prostate cancer cells and non-neoplastic prostate cells treated with CBPD-409. Data are presented as mean +/− SD (n = 6).

**d.** Table summary of CBPD-409 IC50 in prostate cancer and non-neoplastic prostate cells.

**e.** Dose-response curves and IC50 of LNCaP cells treated with CBPD-409, CBPD-409-Me, and GNE-049. Data are presented as mean +/− SD (n = 6).

**f.** Dose-response curves and IC50 of LNCaP parental and GNE-049 resistant (R) cells treated with GNE-049 (left) or CBPD-409 (right). Data are presented as mean +/− SD (n = 6).

**g.** Dose-response curves and IC50 of LNCaP parental and A485 resistant (R) cells treated with A485 (left) or CBPD-409 (right). Data are presented as mean +/− SD (n = 6).

**Figure S8: CBPD-409 monotherapy in the intact VCaP tumor model.**

**a.** Individual tumor images from vehicle and CBPD-409 groups in intact VCaP xenograft study.

**b.** Representative immunohistochemistry images showing staining of murine p300 in specified organs treated with CBPD-409 (3 mg/kg, administered orally three days per week). n=10 per treatment. Scalebar = 50 µm.

**c.** Percent body weight measurement throughout the treatment period of intact VCaP tumor bearing mice in vehicle and CBPD-409 groups (two-sided t-test). Data are presented as mean +/− SEM (vehicle: n = 10; CBPD-409: n = 10).

**d.** Individual prostate images from vehicle (n=5) and CBPD-409 (n=5) groups from intact VCaP xenograft study.

**e.** Percent major organ weight measurements (organ weight vs. body weight) from intact VCaP xenograft study. Data are presented as mean +/− SD (n = 6).

**f.** Liver function and kidney function tests from intact VCaP xenograft study. ALT: alanine transaminase; AST: aspartate transaminase; BUN: blood urea nitrogen; CREA: creatinine. Vehicle: n=5; CBPD-409: n=5; two-sided t-test.

**g.** Complete blood count from intact VCaP xenograft study. EOS: eosinophils; BAS: basophils; NEU: neutrophils; LYM: lymphocytes; MONO: monocytes; RBC, red blood cells; HGB: hemoglobin; HCT: hematocrit; MCV: mean corpuscular volume; MCH: mean corpuscular hemoglobin; MCHC: mean corpuscular hemoglobin concentration; RDW: red cell distribution width; PLT, platelets; MPV: mean platelet volume. Data are presented as mean +/− SD (n = 6, biological replicates).

**Figure S9: CBPD-409 does not exhibit toxicity in CD1 mice and CD rats.**

**a.** Schematic of experimental design for CD1 mouse study. GI, gastrointestinal; GU, genitourinary; LN, lymph node.

**b.** Representative immunohistochemistry images showing staining of murine CBP in specified organs of CD-1 IGS mice treated with vehicle or CBPD-409 (3 mg/kg, administered orally three days per week). n=10 per treatment. Scalebar = 50 µm.

**c.** Percent major organ weight measurements (organ weight vs. body weight) from CD1 mouse study. Data are presented as mean +/− SD (n = 10, biological replicates).

**d.** Representative H&E images of mouse testes from CD1 study. Right panel showing quantification of germ cell density and maturation using the Johnsen scoring system (two-sided t-test). Testes were harvested at 5 days and 32 days post-CBPD-409 treatment. Treatment was discontinued after day 32, and a subsequent set of testes was collected 26 days later (total 58 days) for evaluation. Scalebar = 50 µm.

**e.** Representative immunohistochemistry images showing staining of CBP and p300 in specified organs of CD rats treated with vehicle or CBPD-409 (10 mg/kg, administered orally three days per week). n=5 per treatment. Scalebar = 20 µm.

**f.** Representative H&E staining of major organs from CD rat study. Scalebar = 20 µm.

**Figure S10: CBPD-409 efficacy in castrated VCaP-CRPC tumor model.**

**a.** Percent body weight measurement throughout the treatment period of VCaP-CRPC tumor bearing mice in vehicle, Enza, CBPD-409, and combination of Enza with CBPD-409 groups (two-sided t-test). Data are presented as mean +/− SEM. Vehicle: n = 9; Enza: n=9; CBPD-409: n = 9; Enza+CBPD-409: n=9.

**b.** Representative haematoxylin and eosin (H&E) staining and immunohistochemistry for the indicated protein from the VCaP-CRPC xenograft study (n = 4 tumors per treatment). Scalebar = 50 µm.

**c.** Immunoblots of indicated proteins and histone marks from VCaP-CRPC xenograft study. The tumors were collected after 5 days of treatment. Vehicle: n = 4; Enza: n=4; CBPD-409: n = 4; Enza+CBPD-409: n=6.

**Figure S11: CBPD-409 efficacy in enzalutamide-resistant tumor models.**

**a.** Percent body weight measurement throughout the treatment period from MDA-PCa-146-12 CRPC xenograft study (two-sided t-test). Data are presented as mean +/− SEM. Vehicle: n = 6; Enza: n=6; Enza+CBPD-409: n=7.

**b.** Individual tumor images of vehicle, Enza, and Enza+CBPD-409 groups from MDA-PCa-146-12 CRPC study.

**c.** Representative H&E staining and immunohistochemistry staining from the MDA-PCa-146-12 CRPC xenograft study (n = 10 tumors per treatment). Scalebar = 200 µm.

**d.** Graph depicting the tumor volume curve in the WA-74 CRPC xenograft model, measured biweekly by calipers. Statistical analysis was performed using a two-sided t-test. Data are presented as mean ± SEM. Sample sizes are as follows: vehicle, n = 15; Enza, n=16; CBPD-409, n = 12.

**e.** Representative immunohistochemistry from the WA-74 CRPC xenograft study (n = 10 tumors per treatment). Scalebar = 50 µm.

## Notes

### Summary of Updates

The revised manuscript contains several new pieces of data in the main and supplemental figures.

