## Supplemental Figures S1 - S11 for "p300/CBP degradation is required to disable the active AR enhanceosome in prostate cancer"

### Figure S1

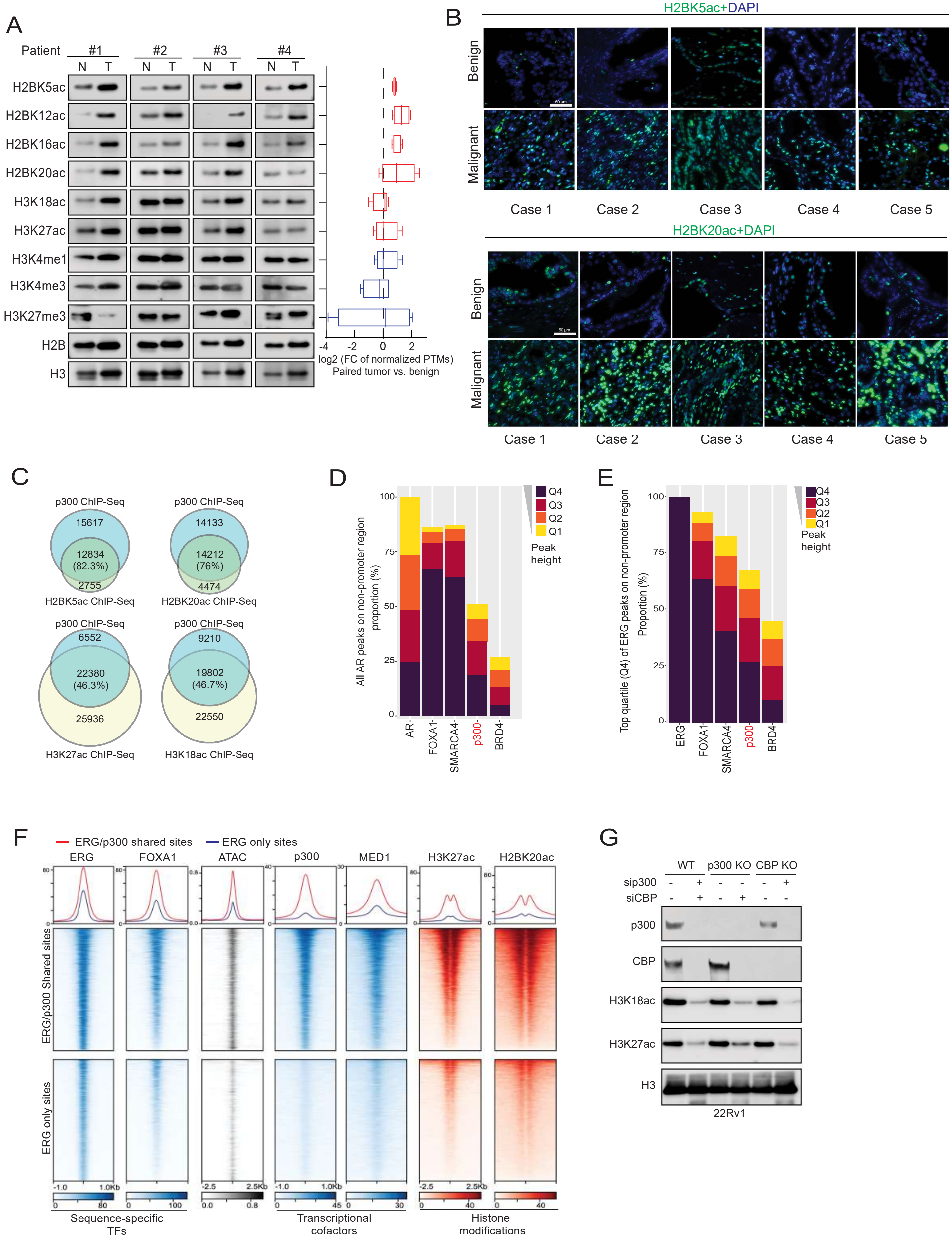

A
Patient #1 #2 #3 #4
N T N T N T N T
H2BK5ac
H2BK12ac
H2BK16ac
H2BK20ac
H3K18ac
H3K27ac
H3K4me1
H3K4me3
H3K27me3
H2B
H3
-4 -2 0 2
log2 (FC of normalized PTMs)
Paired tumor vs. benign
B
H2BK5ac+DAPI
Benign
Malignant
Case 1 Case 2 Case 3 Case 4 Case 5
H2BK20ac+DAPI
Benign
Malignant
Case 1 Case 2 Case 3 Case 4 Case 5
C
p300 ChIP-Seq
15617
12834 (82.3%)
2755
H2BK5ac ChIP-Seq
p300 ChIP-Seq
14133
14212 (76%)
4474
H2BK20ac ChIP-Seq
p300 ChIP-Seq
6552
22380 (46.3%)
25936
H3K27ac ChIP-Seq
p300 ChIP-Seq
9210
19802 (46.7%)
22550
H3K18ac ChIP-Seq
D
All AR peaks on non-promoter region proportion (%)
AR FOXA1 SMARCA4 p300 BRD4
Q4 Q3 Q2 Q1
Peak height
E
Top quartile (Q4) of ERG peaks on non-promoter region Proportion (%)
ERG FOXA1 SMARCA4 p300 BRD4
Q4 Q3 Q2 Q1
Peak height
F
ERG/p300 shared sites
ERG only sites
ERG FOXA1 ATAC p300 MED1 H3K27ac H2BK20ac
ERG/p300 Shared sites
ERG only sites
-1.0 1.0Kb
-2.5 2.5Kb
Sequence-specific TFs
Transcriptional cofactors
Histone modifications
G
WT p300 KO CBP KO
sip300 - + - - - +
siCBP - + - + - -
p300
CBP
H3K18ac
H3K27ac
H3
22Rv1

Figure S2

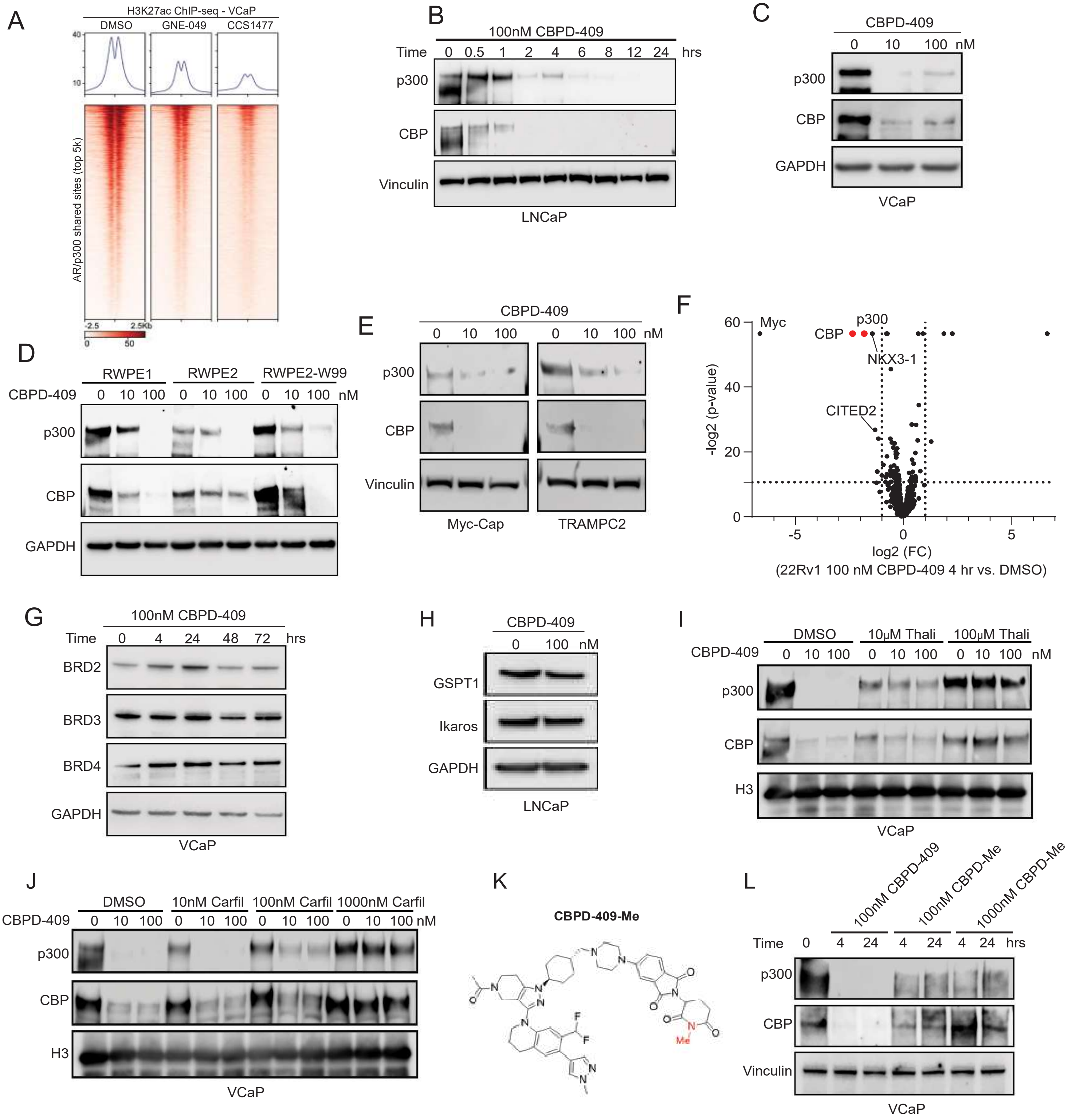

A
H3K27ac ChIP-seq - VCaP
DMSO
GNE-049
CCS1477
40
10
AR/p300 shared sites (top 5k)
-2.5
2.5Kb
0
50
B
100nM CBPD-409
Time 0 0.5 1 2 4 6 8 12 24 hrs
p300
CBP
Vinculin
LNCaP
C
CBPD-409
0 10 100 nM
p300
CBP
GAPDH
VCaP
D
RWPE1
RWPE2
RWPE2-W99
CBPD-409 0 10 100 0 10 100 0 10 100 nM
p300
CBP
GAPDH
E
CBPD-409
0 10 100 0 10 100 nM
p300
CBP
Vinculin
Myc-Cap
TRAMPC2
F
60
40
20
0
Myc
CBP
p300
NKX3-1
CITED2
-log2 (p-value)
-5
0
5
log2 (FC)
(22Rv1 100 nM CBPD-409 4 hr vs. DMSO)
G
100nM CBPD-409
Time 0 4 24 48 72 hrs
BRD2
BRD3
BRD4
GAPDH
VCaP
H
CBPD-409
0 100 nM
GSPT1
Ikaros
GAPDH
LNCaP
I
DMSO
10μM Thali
100μM Thali
CBPD-409 0 10 100 0 10 100 0 10 100 nM
p300
CBP
H3
VCaP
J
DMSO
10nM Carfil
100nM Carfil
1000nM Carfil
CBPD-409 0 10 100 0 10 100 0 10 100 0 10 100 nM
p300
CBP
H3
VCaP
K
CBPD-409-Me
F
F
N
N
O
Me
L
100nM CBPD-409
100nM CBPD-Me
1000nM CBPD-Me
Time 0 4 24 4 24 4 24 hrs
p300
CBP
Vinculin
VCaP

### Figure S3

A

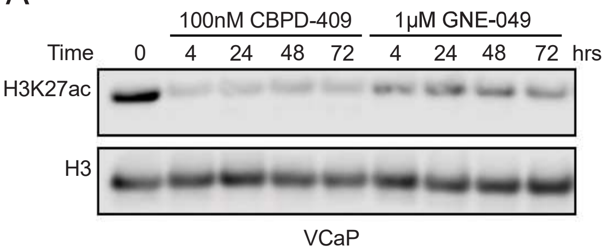


B

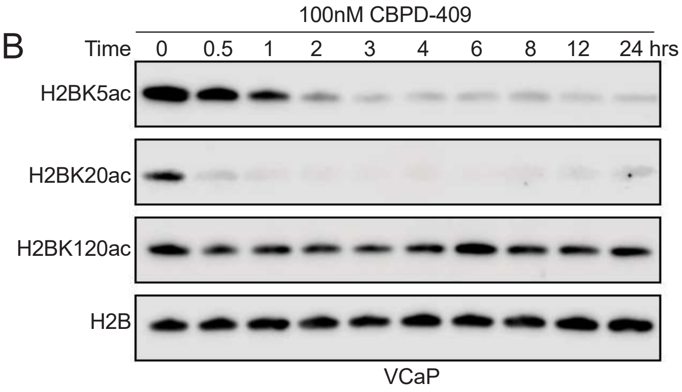


C

Summary of differential acetylated sites (acetylated proteins)

| Group Name | Up-regulated (FC>1.5, p<0.05) | Down-regulated (FC<1/1.5, p<0.05) |
|---|---|---|
| DMSO vs CBPD 1hr | 16 (11) | 134 (80) |
| DMSO vs CBPD 4hrs | 4 (2) | 243 (151) |
| DMSO vs CBPD 12hrs | 5 (5) | 302 (197) |
| DMSO vs CBPD 24hrs | 11 (8) | 337 (214) |

D

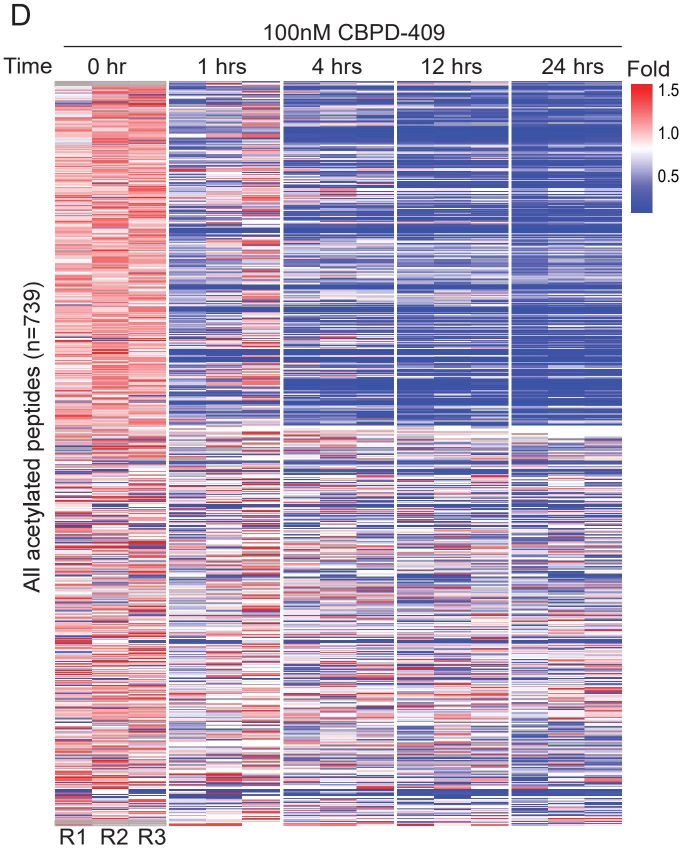


E

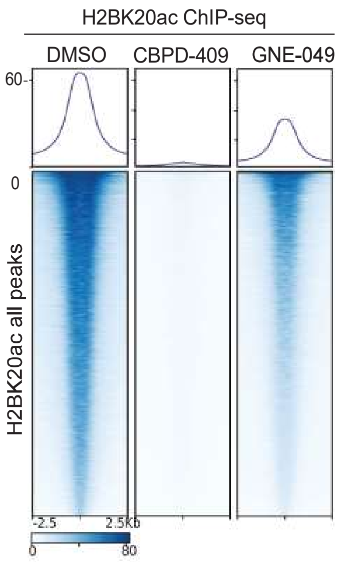

Figure S4

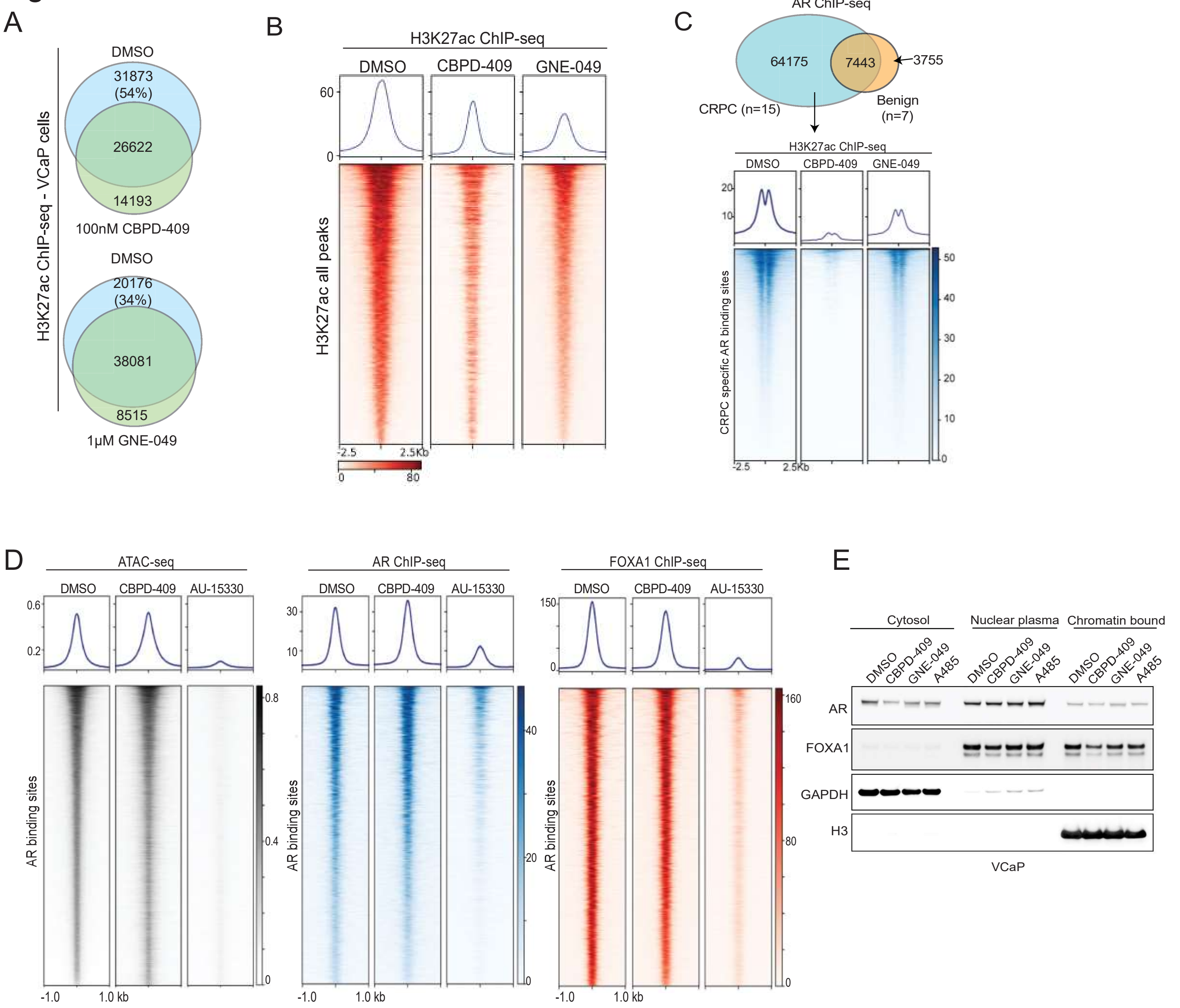

A
H3K27ac ChIP-seq - VCaP cells
DMSO
31873
(54%)
26622
14193
100nM CBPD-409
DMSO
20176
(34%)
38081
8515
1μM GNE-049
B
H3K27ac ChIP-seq
DMSO
CBPD-409
GNE-049
60
0
H3K27ac all peaks
-2.5
2.5Kb
0
80
C
AR ChIP-seq
64175
7443
3755
CRPC (n=15)
Benign
(n=7)
H3K27ac ChIP-seq
DMSO
CBPD-409
GNE-049
20
10
CRPC specific AR binding sites
50
40
30
20
10
0
-2.5
2.5Kb
D
ATAC-seq
DMSO
CBPD-409
AU-15330
0.6
0.2
AR binding sites
0.8
0.4
0
-1.0
1.0 kb
AR ChIP-seq
DMSO
CBPD-409
AU-15330
30
10
AR binding sites
40
20
0
-1.0
1.0 kb
FOXA1 ChIP-seq
DMSO
CBPD-409
AU-15330
150
0
AR binding sites
160
80
0
-1.0
1.0 kb
E
Cytosol
Nuclear plasma
Chromatin bound
DMSO
CBPD-409
GNE-049
A485
AR
FOXA1
GAPDH
H3
VCaP

### Figure S5

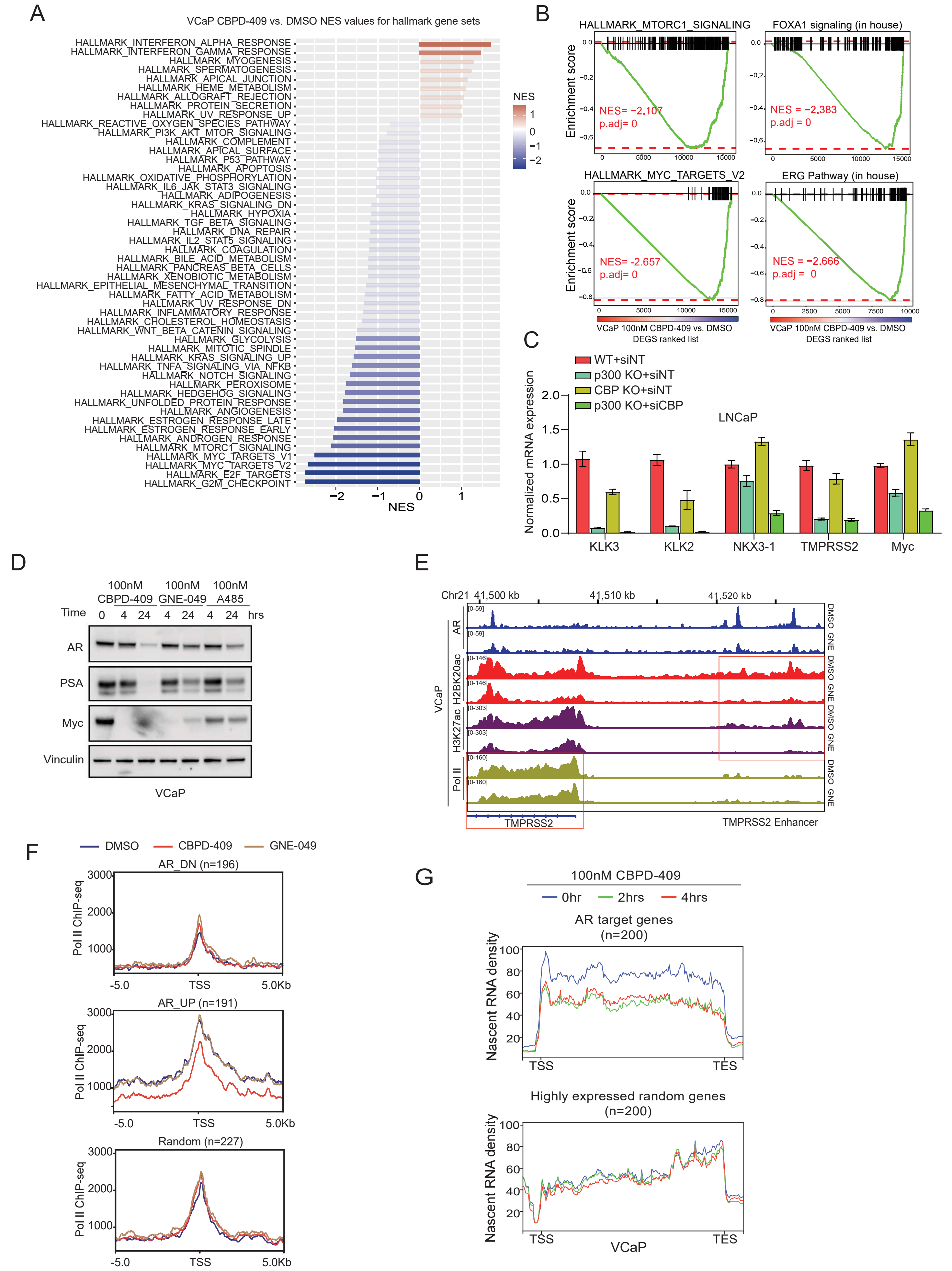
A
VCaP CBPD-409 vs. DMSO NES values for hallmark gene sets
HALLMARK_INTERFERON_ALPHA_RESPONSE
HALLMARK_INTERFERON_GAMMA_RESPONSE
HALLMARK_MYOGENESIS
HALLMARK_SPERMATOGENESIS
HALLMARK_APICAL_JUNCTION
HALLMARK_HEME_METABOLISM
HALLMARK_ALLOGRAFT_REJECTION
HALLMARK_PROTEIN_SECRETION
HALLMARK_UV_RESPONSE_UP
HALLMARK_REACTIVE_OXYGEN_SPECIES_PATHWAY
HALLMARK_PI3K_AKT_MTOR_SIGNALING
HALLMARK_COMPLEMENT
HALLMARK_APICAL_SURFACE
HALLMARK_P53_PATHWAY
HALLMARK_APOPTOSIS
HALLMARK_OXIDATIVE_PHOSPHORYLATION
HALLMARK_IL6_JAK_STAT3_SIGNALING
HALLMARK_ADIPOGENESIS
HALLMARK_KRAS_SIGNALING_DN
HALLMARK_HYPOXIA
HALLMARK_TGF_BETA_SIGNALING
HALLMARK_DNA_REPAIR
HALLMARK_IL2_STAT5_SIGNALING
HALLMARK_COAGULATION
HALLMARK_BILE_ACID_METABOLISM
HALLMARK_PANCREAS_BETA_CELLS
HALLMARK_XENOBIOTIC_METABOLISM
HALLMARK_EPITHELIAL_MESENCHYMAL_TRANSITION
HALLMARK_FATTY_ACID_METABOLISM
HALLMARK_UV_RESPONSE_DN
HALLMARK_INFLAMMATORY_RESPONSE
HALLMARK_CHOLESTEROL_HOMEOSTASIS
HALLMARK_WNT_BETA_CATENIN_SIGNALING
HALLMARK_GLYCOLYSIS
HALLMARK_MITOTIC_SPINDLE
HALLMARK_KRAS_SIGNALING_UP
HALLMARK_TNFA_SIGNALING_VIA_NFKB
HALLMARK_NOTCH_SIGNALING
HALLMARK_PEROXISOME
HALLMARK_HEDGEHOG_SIGNALING
HALLMARK_UNFOLDED_PROTEIN_RESPONSE
HALLMARK_ANGIOGENESIS
HALLMARK_ESTROGEN_RESPONSE_LATE
HALLMARK_ESTROGEN_RESPONSE_EARLY
HALLMARK_ANDROGEN_RESPONSE
HALLMARK_MTORC1_SIGNALING
HALLMARK_MYC_TARGETS_V1
HALLMARK_MYC_TARGETS_V2
HALLMARK_E2F_TARGETS
HALLMARK_G2M_CHECKPOINT
−2 −1 0 1
NES
NES 1 0 −1 −2
B
HALLMARK_MTORC1_SIGNALING
NES= −2.107
p.adj= 0
FOXA1 signaling (in house)
NES = −2.383
p.adj = 0
HALLMARK_MYC_TARGETS_V2
NES= -2.657
p.adj= 0
ERG Pathway (in house)
NES = −2.666
p.adj = 0
Enrichment score
VCaP 100nM CBPD-409 vs. DMSO
DEGS ranked list
C
WT+siNT
p300 KO+siNT
CBP KO+siNT
p300 KO+siCBP
LNCaP
Normalized mRNA expression
KLK3
KLK2
NKX3-1
TMPRSS2
Myc
D
100nM CBPD-409
100nM GNE-049
100nM A485
Time 0 4 24 4 24 4 24 hrs
AR
PSA
Myc
Vinculin
VCaP
E
Chr21 41,500 kb 41,510 kb 41,520 kb
AR
H2BK20ac
H3K27ac
Pol II
VCaP
DMSO
GNE
TMPRSS2
TMPRSS2 Enhancer
F
DMSO
CBPD-409
GNE-049
AR_DN (n=196)
AR_UP (n=191)
Random (n=227)
Pol II ChIP-seq
-5.0 TSS 5.0Kb
G
100nM CBPD-409
0hr 2hrs 4hrs
AR target genes (n=200)
Highly expressed random genes (n=200)
Nascent RNA density
TSS TES
VCaP

### Figure S6

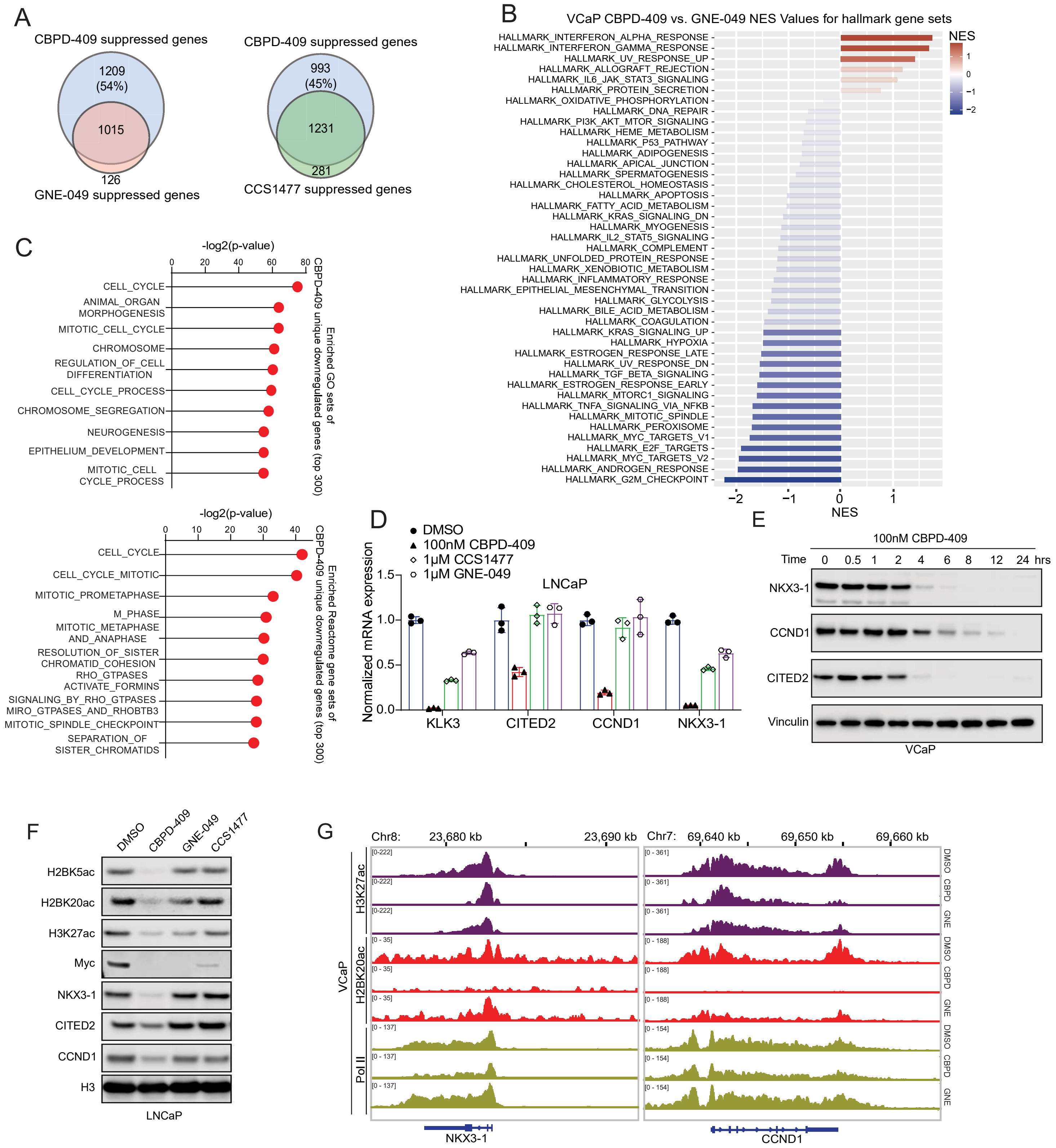

A
CBPD-409 suppressed genes
1209 (54%)
1015
126
GNE-049 suppressed genes
CBPD-409 suppressed genes
993 (45%)
1231
281
CCS1477 suppressed genes
B
VCaP CBPD-409 vs. GNE-049 NES Values for hallmark gene sets
HALLMARK_INTERFERON_ALPHA_RESPONSE
HALLMARK_INTERFERON_GAMMA_RESPONSE
HALLMARK_UV_RESPONSE_UP
HALLMARK_ALLOGRAFT_REJECTION
HALLMARK_IL6_JAK_STAT3_SIGNALING
HALLMARK_PROTEIN_SECRETION
HALLMARK_OXIDATIVE_PHOSPHORYLATION
HALLMARK_DNA_REPAIR
HALLMARK_PI3K_AKT_MTOR_SIGNALING
HALLMARK_HEME_METABOLISM
HALLMARK_P53_PATHWAY
HALLMARK_ADIPOGENESIS
HALLMARK_APICAL_JUNCTION
HALLMARK_SPERMATOGENESIS
HALLMARK_CHOLESTEROL_HOMEOSTASIS
HALLMARK_APOPTOSIS
HALLMARK_FATTY_ACID_METABOLISM
HALLMARK_KRAS_SIGNALING_DN
HALLMARK_MYOGENESIS
HALLMARK_IL2_STAT5_SIGNALING
HALLMARK_COMPLEMENT
HALLMARK_UNFOLDED_PROTEIN_RESPONSE
HALLMARK_XENOBIOTIC_METABOLISM
HALLMARK_INFLAMMATORY_RESPONSE
HALLMARK_EPITHELIAL_MESENCHYMAL_TRANSITION
HALLMARK_GLYCOLYSIS
HALLMARK_BILE_ACID_METABOLISM
HALLMARK_COAGULATION
HALLMARK_KRAS_SIGNALING_UP
HALLMARK_HYPOXIA
HALLMARK_ESTROGEN_RESPONSE_LATE
HALLMARK_UV_RESPONSE_DN
HALLMARK_TGF_BETA_SIGNALING
HALLMARK_ESTROGEN_RESPONSE_EARLY
HALLMARK_MTORC1_SIGNALING
HALLMARK_TNFA_SIGNALING_VIA_NFKB
HALLMARK_MITOTIC_SPINDLE
HALLMARK_PEROXISOME
HALLMARK_MYC_TARGETS_V1
HALLMARK_E2F_TARGETS
HALLMARK_MYC_TARGETS_V2
HALLMARK_ANDROGEN_RESPONSE
HALLMARK_G2M_CHECKPOINT
−2 −1 0 1
NES
NES 1 0 −1 −2
C
-log2(p-value)
0 20 40 60 80
CELL_CYCLE
ANIMAL_ORGAN MORPHOGENESIS
MITOTIC_CELL_CYCLE
CHROMOSOME
REGULATION_OF_CELL DIFFERENTIATION
CELL_CYCLE_PROCESS
CHROMOSOME_SEGREGATION
NEUROGENESIS
EPITHELIUM_DEVELOPMENT
MITOTIC_CELL CYCLE_PROCESS
Enriched GO sets of CBPD-409 unique downregulated genes (top 300)
-log2(p-value)
0 10 20 30 40
CELL_CYCLE
CELL_CYCLE_MITOTIC
MITOTIC_PROMETAPHASE
M_PHASE
MITOTIC_METAPHASE AND_ANAPHASE
RESOLUTION_OF_SISTER CHROMATID_COHESION
RHO_GTPASES ACTIVATE_FORMINS
SIGNALING_BY_RHO_GTPASES MIRO_GTPASES_AND_RHOBTB3
MITOTIC_SPINDLE_CHECKPOINT
SEPARATION_OF SISTER_CHROMATIDS
Enriched Reactome gene sets of CBPD-409 unique downregulated genes (top 300)
D
DMSO
100nM CBPD-409
1μM CCS1477
1μM GNE-049
LNCaP
Normalized mRNA expression
0.0 0.5 1.0 1.5
KLK3 CITED2 CCND1 NKX3-1
E
100nM CBPD-409
Time 0 0.5 1 2 4 6 8 12 24 hrs
NKX3-1
CCND1
CITED2
Vinculin
VCaP
F
DMSO CBPD-409 GNE-049 CCS1477
H2BK5ac
H2BK20ac
H3K27ac
Myc
NKX3-1
CITED2
CCND1
H3
LNCaP
G
Chr8: 23,680 kb 23,690 kb
Chr7: 69,640 kb 69,650 kb 69,660 kb
VCaP
H3K27ac
H2BK20ac
Pol II
[0-222]
[0 - 361]
[0 - 35]
[0 - 188]
[0 - 137]
[0 - 154]
DMSO CBPD GNE
NKX3-1
CCND1

### Figure S7

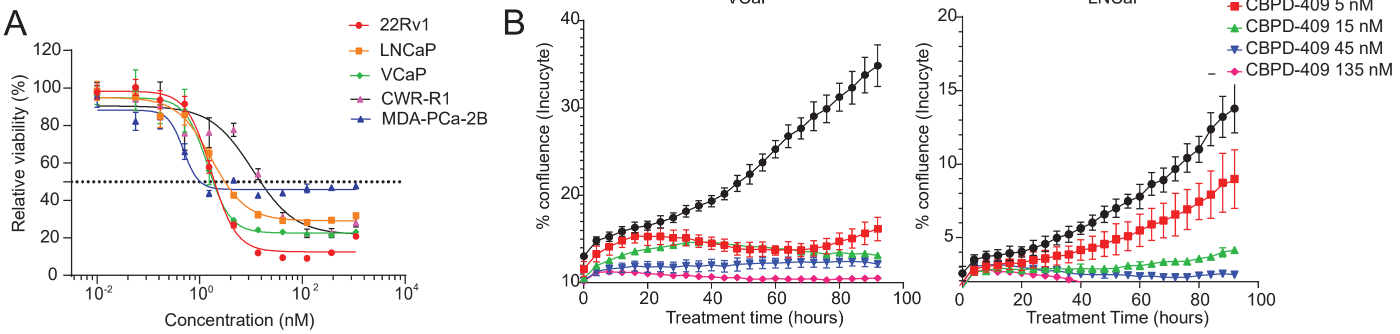


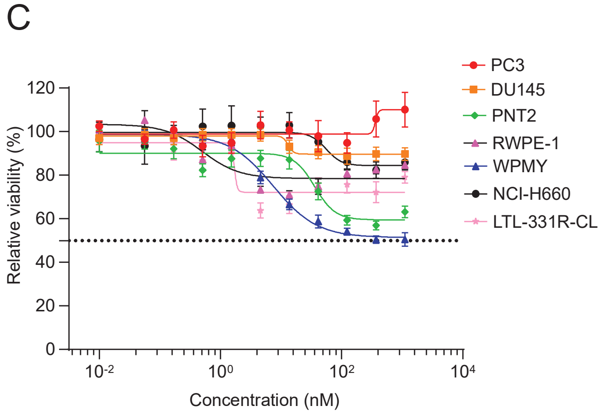


D

| Group | Cell line | CBPD-409 IC50 (nM) |
|---|---|---|
| AR+ prostate cancer | 22Rv1 | 2.2 |
| | MDA-PCA-2B | 2.5 |
| | VCaP | 4 |
| | LNCaP | 11 |
| | CWR-R1 | 11.5 |
| AR- prostate cancer | PC3 | 6368 |
| | DU145 | >30000 |
| | NCI-H660 | >30000 |
| | LTL-331R-CL | >30000 |
| non-neoplastic | WPMY-1 | 4365 |
| | PNT2 | >30000 |
| | RWPE-1 | >30000 |

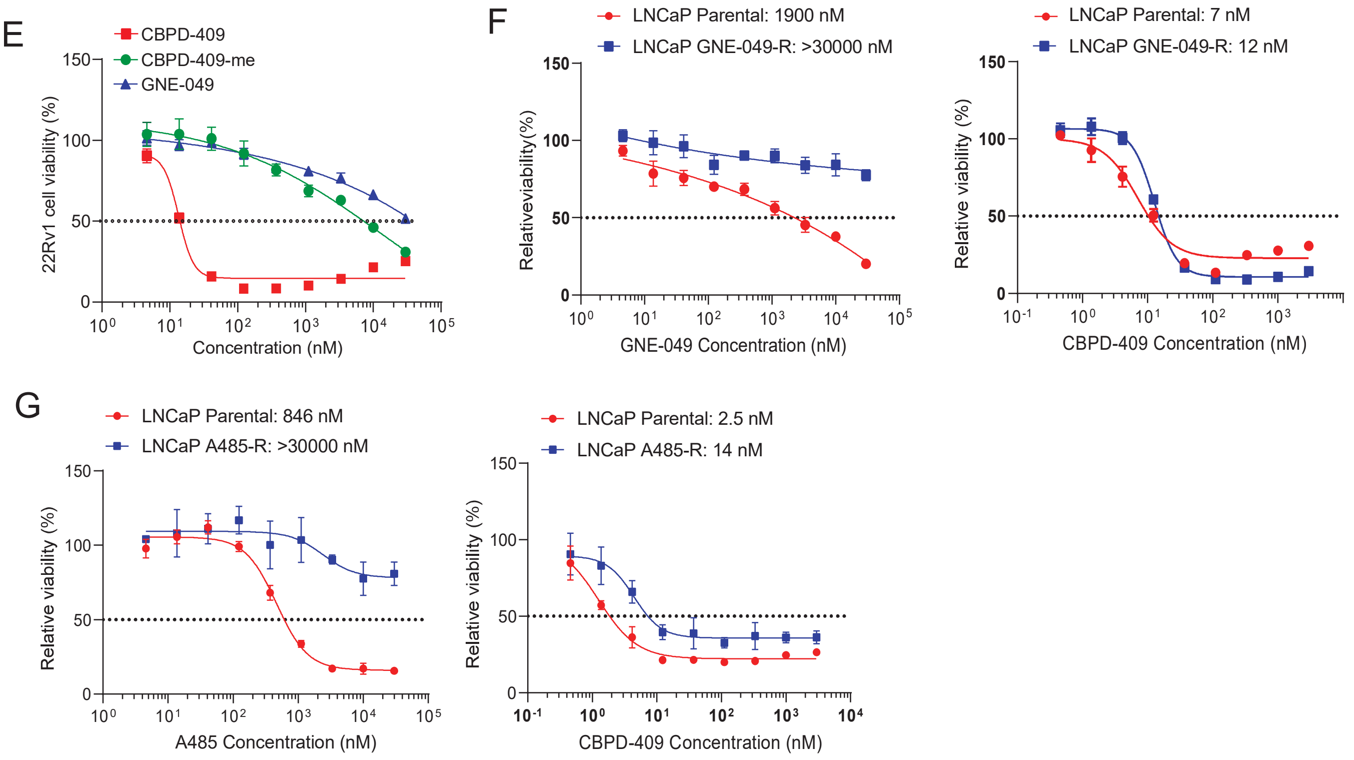

### Figure S8

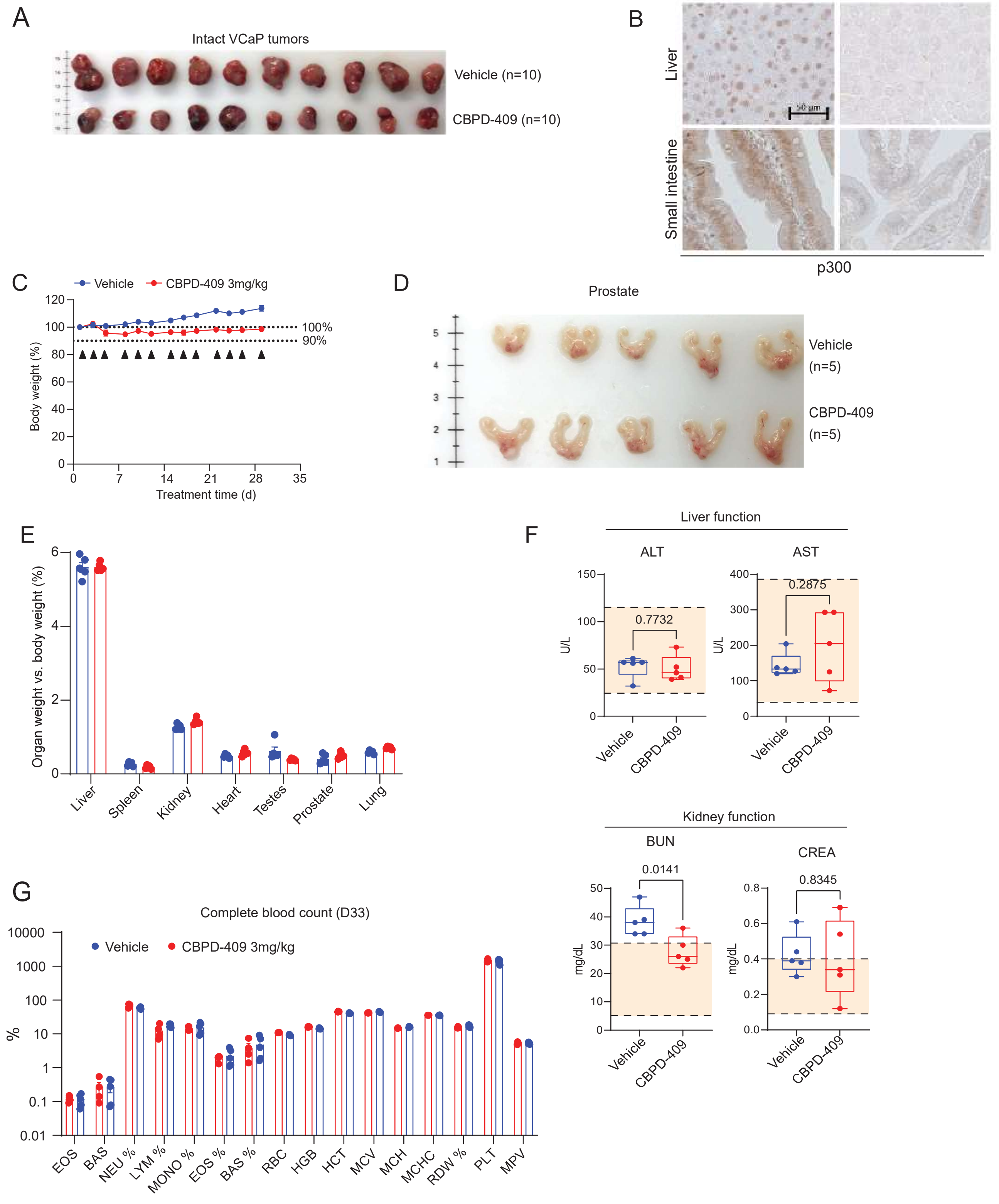
A
Intact VCaP tumors
Vehicle (n=10)
CBPD-409 (n=10)
B
Vehicle
CBPD-409
Liver
Small intestine
50 μm
p300
C
Vehicle
CBPD-409 3mg/kg
Body weight (%)
100%
90%
Treatment time (d)
D
Prostate
Vehicle
(n=5)
CBPD-409
(n=5)
E
Organ weight vs. body weight (%)
Liver
Spleen
Kidney
Heart
Testes
Prostate
Lung
F
Liver function
ALT
AST
U/L
0.7732
0.2875
Vehicle
CBPD-409
Kidney function
BUN
CREA
mg/dL
0.0141
0.8345
G
Complete blood count (D33)
Vehicle
CBPD-409 3mg/kg
%
EOS
BAS
NEU %
LYM %
MONO %
EOS %
BAS %
RBC
HGB
HCT
MCV
MCH
MCHC
RDW %
PLT
MPV

### Figure S9

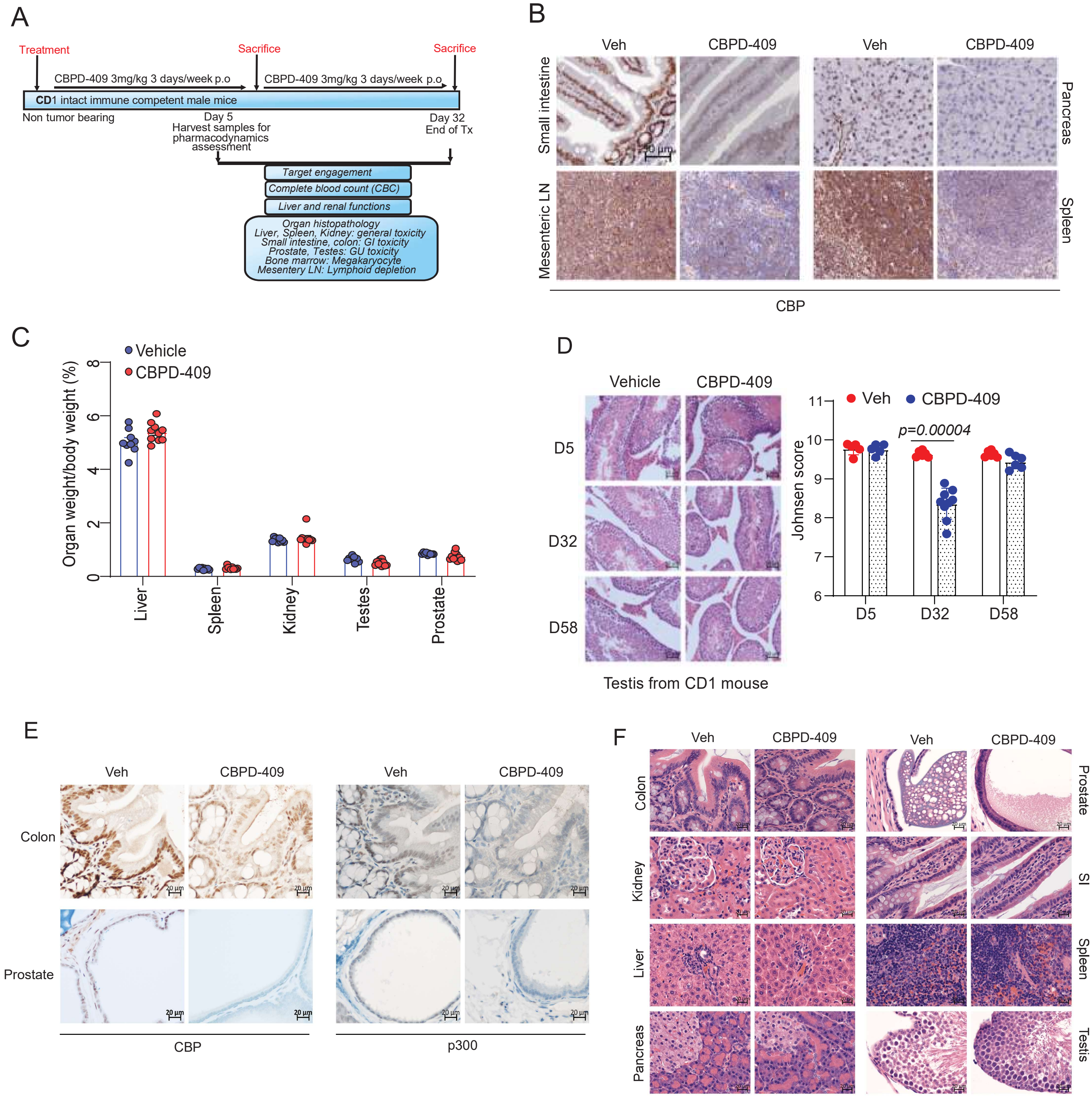

A
Treatment
Sacrifice
Sacrifice
CBPD-409 3mg/kg 3 days/week p.o
CBPD-409 3mg/kg 3 days/week p.o
CD1 intact immune competent male mice
Non tumor bearing
Day 5
Harvest samples for pharmacodynamics assessment
Day 32
End of Tx
Target engagement
Complete blood count (CBC)
Liver and renal functions
Organ histopathology
Liver, Spleen, Kidney: general toxicity
Small intestine, colon: GI toxicity
Prostate, Testes: GU toxicity
Bone marrow: Megakaryocyte
Mesentery LN: Lymphoid depletion
B
Veh
CBPD-409
Veh
CBPD-409
Small intestine
Mesenteric LN
Pancreas
Spleen
CBP
C
Vehicle
CBPD-409
Organ weight/body weight (%)
0
2
4
6
8
Liver
Spleen
Kidney
Testes
Prostate
D
Vehicle
CBPD-409
D5
D32
D58
Testis from CD1 mouse
Veh
CBPD-409
p=0.00004
Johnsen score
6
7
8
9
10
11
D5
D32
D58
E
Veh
CBPD-409
Veh
CBPD-409
Colon
Prostate
CBP
p300
20 μm
F
Veh
CBPD-409
Veh
CBPD-409
Colon
Kidney
Liver
Pancreas
Prostate
SI
Spleen
Testis

### Figure S10

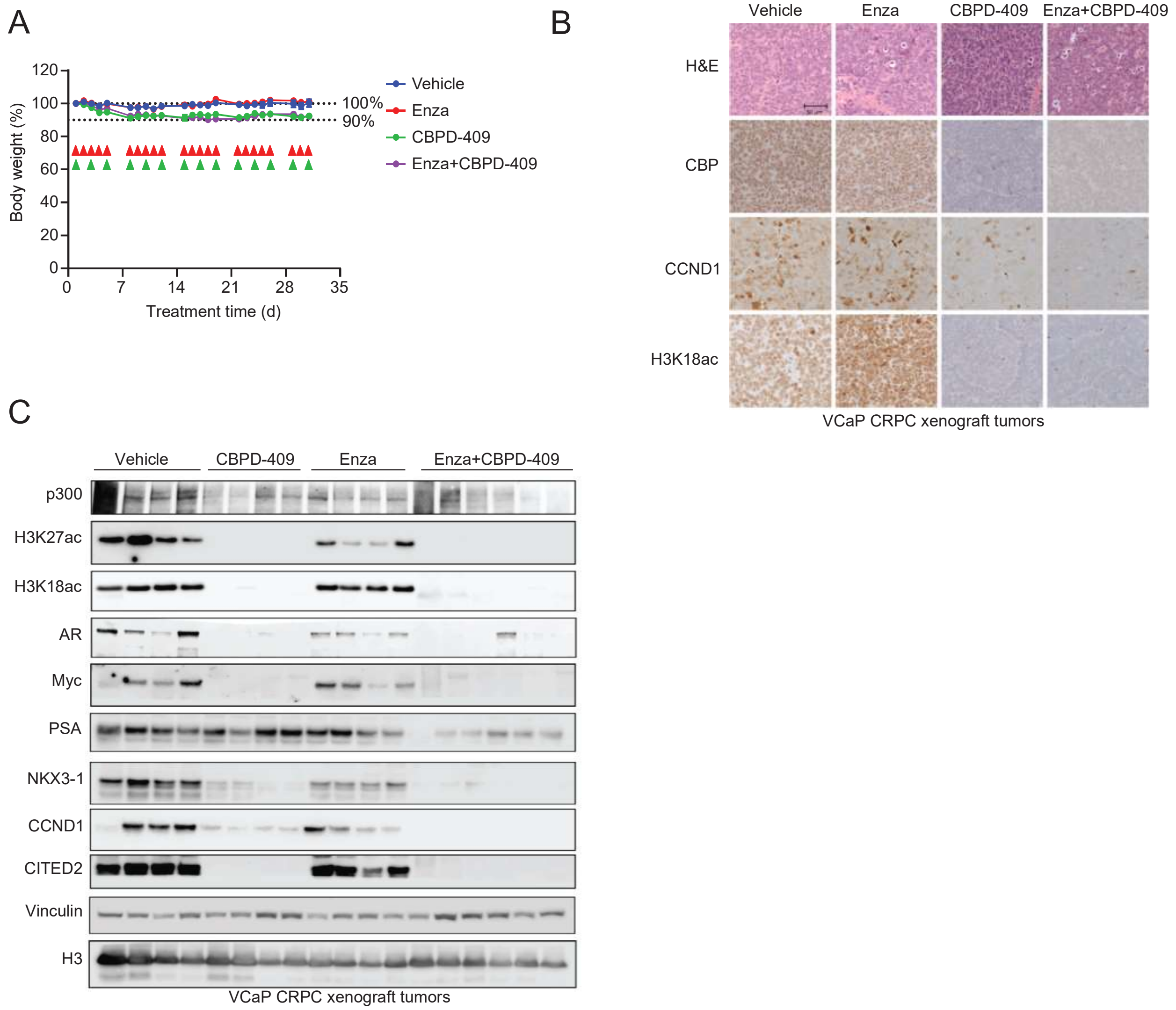
A
B
C
Body weight (%)
120
100
80
60
40
20
0
0
7
14
21
28
35
Treatment time (d)
100%
90%
Vehicle
Enza
CBPD-409
Enza+CBPD-409
Vehicle
Enza
CBPD-409
Enza+CBPD-409
H&E
CBP
CCND1
H3K18ac
VCaP CRPC xenograft tumors
Vehicle
CBPD-409
Enza
Enza+CBPD-409
p300
H3K27ac
H3K18ac
AR
Myc
PSA
NKX3-1
CCND1
CITED2
Vinculin
H3
VCaP CRPC xenograft tumors

### Figure S11

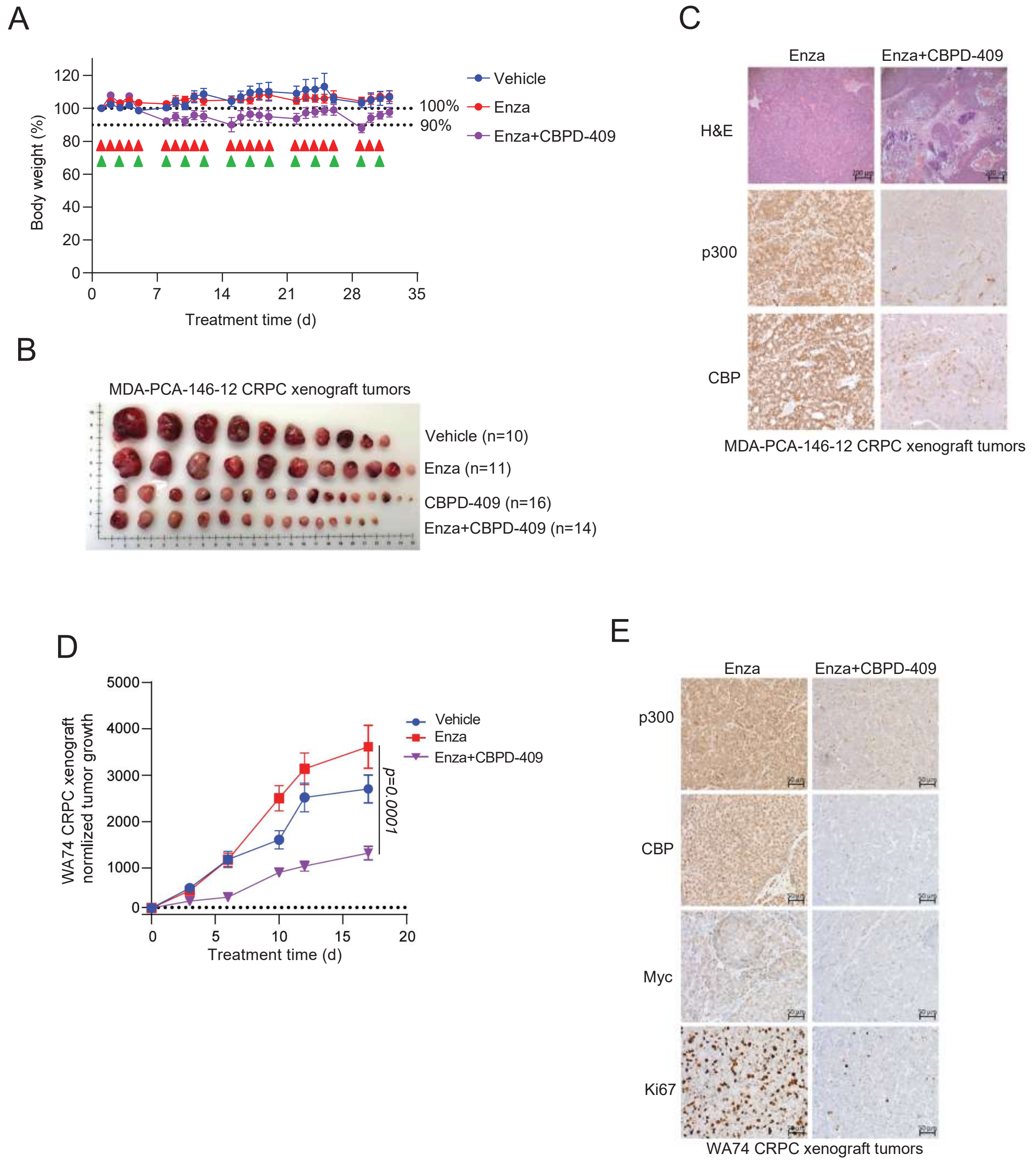

A
Body weight (%)
120
100
80
60
40
20
0
0
7
14
21
28
35
Treatment time (d)
100%
90%
Vehicle
Enza
Enza+CBPD-409
B
MDA-PCA-146-12 CRPC xenograft tumors
Vehicle (n=10)
Enza (n=11)
CBPD-409 (n=16)
Enza+CBPD-409 (n=14)
C
Enza
Enza+CBPD-409
H&E
p300
CBP
MDA-PCA-146-12 CRPC xenograft tumors
D
WA74 CRPC xenograft normlized tumor growth
5000
4000
3000
2000
1000
0
0
5
10
15
20
Treatment time (d)
Vehicle
Enza
Enza+CBPD-409
p=0.0001
E
Enza
Enza+CBPD-409
p300
CBP
Myc
Ki67
WA74 CRPC xenograft tumors
