## Supplemental Methods for "p300/CBP degradation is required to disable the active AR enhanceosome in prostate cancer"

### Supplementary Methods

#### Synthetic Chemistry for CBPD-409 and CBPD-409-Me

##### 1. General chemistry information

All commercial materials were utilized in their original form, unless explicitly stated otherwise. NMR spectra were captured using a Bruker Ascend$^{TM}$ 400 MHz spectrometer, with calibration conducted using residual solvent peaks as internal references. Spectral data were presented in the ($\delta$) chemical shift (multiplicity, J values in Hz, integration) format, employing abbreviations such as s = singlet, d = doublet, t = triplet, q = quartet, hept = heptet, dd = doublet of doublets, and m = multiplet. Low-resolution mass spectrometry (MS) analysis was performed using a Waters UPLC ACQUITY QDa mass spectrometer. High-resolution mass experiments were executed on an Agilent Technologies 6230 TOF LC/MS instrument with APCI ionization. Flash column chromatography was undertaken with a Teledyne CombiFlash RF+ using RediSep Rf silica gel flash columns. The final compounds and some intermediates underwent purification using a C18 reversed-phase preparative HPLC column (SunFire$^{TM}$ Prep C18 OBD$^{TM}$ 5 µm, 50×100 mm) with solvent A (0.1% TFA in $H_2O$) and solvent B (0.1% TFA in MeCN) as eluents at a flow rate of 60 mL/min. The purity of all final compounds was evaluated through UPLC-MS analysis (10-100% MeCN in $H_2O$ containing 0.1% formic acid in 5 min, 1.0 mL/min flow rate) with a C18 column (ACQUITY UPLC BEH C18 1.7 µm, 2.1 × 50 mm).

Abbreviations used: $CDCl_3$, deuterated chloroform; $Cs_2CO_3$, cesium carbonate; DCM, dichloromethane; DMAP, 4-dimethylaminopyridine; DMF, *N,N'*-Dimethylformamide; DMSO, dimethyl sulfoxide; DIBAL, diisobutylaluminum hydride; DIPEA, *N,N'*-Diisopropylethylamine; $Et_3N$, triethylamine; EtOAc, ethyl acetate; HCl, hydrochloric acid; $K_2CO_3$, potassium carbonate; KOH, potassium hydroxide; MeCN, acetonitrile; MeI, methyl iodide; MeOH, methanol; $N_2$, nitrogen; $Na_2SO_4$, sodium sulfate; $NaBH(OAc)_3$, sodium triacetoxyborohydride; NaH, sodium hydride; NaOH, sodium hydroxide; NBS, N-bromosuccinimide; $PdCl_2$(dppf), palladium(II) chloride-bis(diphenylphosphino)ferrocene; RuPhos, ruthenium-based ligand (Phospha-methyl-di(tert-butyl)phenylphosphine); RuPhos Pd G2, second-generation ruthenium-phosphine ligand; *t*-BuONa, sodium tert-butoxide; TFA, trifluoroacetic acid; THF, tetrahydrofuran.

#### 2. Method and procedure for the preparation of CBPD-409

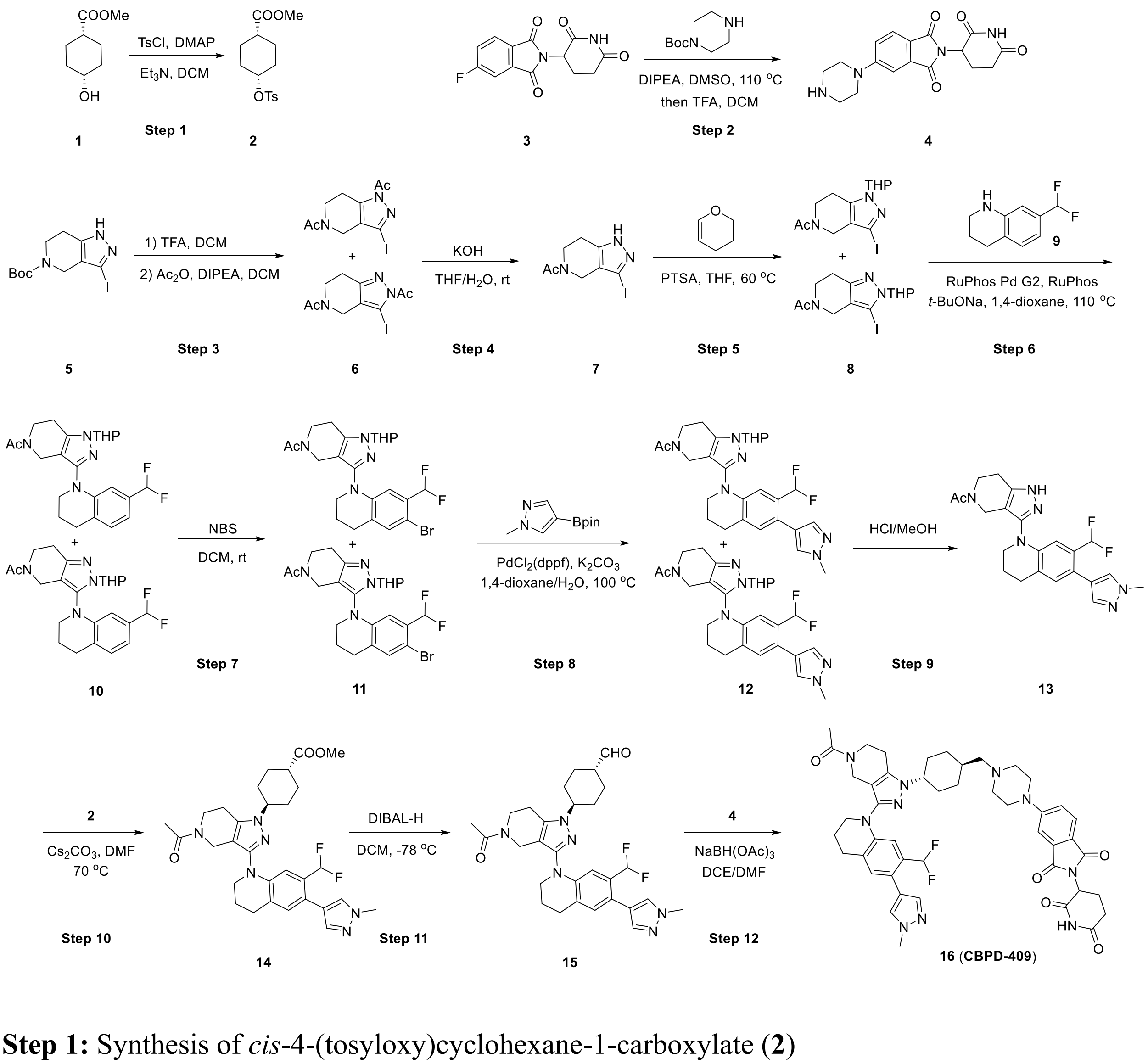


**Step 1:** Synthesis of *cis*-4-(tosyloxy)cyclohexane-1-carboxylate (**2**)

To a solution of methyl *cis*-4-hydroxycyclohexane-1-carboxylate (**1**, 5 g, 1.0 eq) in DCM (50 mL), 4-methylbenzenesulfonyl chloride (9.0 g, 1.5 eq), DMAP (0.77 g, 0.2 eq) and $Et_3N$ (13.2 mL, 3.0 eq) were sequentially added. The resulting mixture was stirred at room temperature for 16 h. Following this, water was introduced, and the resulting mixture was subjected to extraction with DCM. The combined organic layer was then washed with brine, dried over anhydrous $Na_2SO_4$, filtered, and concentrated under reduced pressure. The resulting crude residue was subsequently subjected to purification through flash column chromatography (20% EtOAc/*n*-hexane) to give *cis*-4-(tosyloxy)cyclohexane-1-carboxylate (**2**) as a yellow oil (9.3 g, 94% yield). LC-MS: *m/z* $[M+H]^+$ 334.85; $^1$H NMR (400 MHz, Chloroform-*d*) $\delta$ 7.81 – 7.75 (m, 2H), 7.35 – 7.29 (m, 2H), 4.70 (tt, J = 5.0, 2.9 Hz, 1H), 3.66 (s, 3H), 2.44 (s, 3H), 2.37 – 2.26 (m, 1H), 1.92 – 1.78 (m, 4H), 1.75 – 1.65 (m, 2H), 1.59 – 1.47 (m, 2H).

**Step 2:** Synthesis of 2-(2,6-dioxopiperidin-3-yl)-5-(piperazin-1-yl)isoindoline-1,3-dione (**4**)

To a mixture of 2-(2,6-dioxopiperidin-3-yl)-5-fluoroisoindoline-1,3-dione (**3**, 570 mg, 1.0 eq) and *tert*-butyl piperazine-1-carboxylate (769 mg, 2.0 eq) in DMSO (10 mL) was added DIPEA (1.44 mL). Upon cooling to room temperature, the reaction mixture was diluted with DCM, followed by washing with brine, drying over anhydrous $Na_2SO_4$, and concentration under reduced pressure. The resulting residue underwent purification via flash column chromatography (using a gradient of 0-5% MeOH/DCM) to yield a pure product, which was then subjected to treatment with TFA, resulting in the formation of 2-(2,6-dioxopiperidin-3-yl)-5-(piperazin-1-yl)isoindoline-1,3-dione (**4**) as a light yellow solid (850 mg, yield = 90%). UPLC-MS [M+H]+ 343.15; $^1$H NMR (400 MHz, MeOH-$d_4$) $\delta$ 7.77 (d, *J* = 8.5 Hz, 1H), 7.48 (d, *J* = 2.3 Hz, 1H), 7.35 (dd, *J* = 8.5, 2.4 Hz, 1H), 5.10 (dd, *J* = 12.5, 5.5 Hz, 1H), 3.76 – 3.65 (m, 4H), 3.43 – 3.37 (m, 4H), 2.93 – 2.81 (m, 1H), 2.80 – 2.66 (m, 2H), 2.16 – 2.08 (m, 1H).

**Step 3:** Synthesis of 1,1'-(3-iodo-6,7-dihydro-1H-pyrazolo[4,3-c]pyridine-1,5(4H)-diyl)bis(ethan-1-one) and 1,1'-(3-iodo-6,7-dihydro-2H-pyrazolo[4,3-c]pyridine-2,5(4H)-diyl)bis(ethan-1-one) (**6**)

A solution of *tert*-butyl 3-iodo-1,4,6,7-tetrahydro-5H-pyrazolo[4,3-c]pyridine-5-carboxylate (**5**, 2.5 g, 1.0 eq) in DCM (30 mL) was treated with TFA (10 mL). The reaction mixture was stirred at room temperature for 30 minutes, after which it was concentrated under reduced pressure. The resulting residue was dissolved in DCM (30 mL), and the solution was cooled to 0 °C. DIPEA (6.2 mL, 5.0 eq) was then added, followed by acetic anhydride (1.8 mL, 2.5 eq). The reaction was allowed to warm to room temperature and was stirred for 20 h. After completion, the reaction mixture was diluted with DCM, washed with brine, dried over anhydrous $Na_2SO_4$, and concentrated under reduced pressure to remove the solvent. The resulting crude product was subjected to purification via flash column chromatography (using a gradient of 0-5% MeOH/DCM) to afford 1,1'-(3-iodo-6,7-dihydro-1H-pyrazolo[4,3-c]pyridine-1,5(4H)-diyl)bis(ethan-1-one) and 1,1'-(3-iodo-6,7-dihydro-2H-pyrazolo[4,3-c]pyridine-2,5(4H)-diyl)bis(ethan-1-one) (**6**) as a white solid (2.2 g) in a yield of 92%. LC-MS *m/z* $[M+H]^+$ 334.04; $^1H$ NMR (400 MHz, Chloroform-*d*) $\delta$ 4.46 – 4.22 (m, 2H), 3.88 – 3.65 (m, 2H), 3.20 – 3.07 (m, 2H), 2.67 (s, 3H), 2.19 (s, 3H).

**Step 4:** Synthesis of 1-(3-iodo-1,4,6,7-tetrahydro-5H-pyrazolo[4,3-c]pyridin-5-yl)ethan-1-one (**7**)

A solution of compound **6** (3.4 g, 1.0 eq) in THF/$H_2O$ (40 mL/20 mL) was treated with KOH (2.9 g, 5.0 eq). The reaction mixture was stirred at room temperature for 3 h. Following this, the solvent was removed under reduced pressure, and the resulting residue was dissolved in water. The solution was then neutralized with 2 N aqueous HCl to achieve a pH of 8~9. The neutralized solution was subsequently extracted with EtOAc. The combined organic layers were washed with brine, dried over anhydrous $Na_2SO_4$, and concentrated to yield 1-(3-iodo-1,4,6,7-tetrahydro-5H-pyrazolo[4,3-c]pyridin-5-yl)ethan-1-one (**7**) as a white solid (2.7 g), with a yield of 91%. LC-MS *m/z* [M+H]$^+$ 291.89; $^1$H NMR (400 MHz, Methanol-$d_4$) $\delta$ 12.97 (s, 1H), 4.22 (s, 2H), 3.76 – 3.60 (m, 2H), 2.77 – 2.54 (m, 2H), 2.13 – 2.05 (m, 3H).

**Step 5:** Synthesis of 1-(3-iodo-1-(tetrahydro-2H-pyran-2-yl)-1,4,6,7-tetrahydro-5H-pyrazolo[4,3-c]pyridin-5-yl)ethan-1-one and 1-(3-iodo-2-(tetrahydro-2H-pyran-2-yl)-2,4,6,7-tetrahydro-5H-pyrazolo[4,3-c]pyridin-5-yl)ethan-1-one (**8**)

In a solution containing compound **7** (3.7 g, 1.0 eq) and 3,4-dihydro-2*H*-pyran (3.5 mL, 3.0 eq) in THF (60 mL), *p*-toluenesulfonic acid (2.2 g, 1.0 eq) was added. The reaction mixture was stirred at 65 °C for 24 h. After completion, the reaction mixture was concentrated, then diluted with DCM. The resulting solution was washed with brine, dried over anhydrous $Na_2SO_4$, and concentrated under reduced pressure to remove the solvent. The residue was subjected to purification by flash column chromatography (using a gradient of 0-100% EtOAc/*n*-hexane) to afford 1-(3-iodo-1-(tetrahydro-2H-pyran-2-yl)-1,4,6,7-tetrahydro-5H-pyrazolo[4,3-c]pyridin-5-yl)ethan-1-one and 1-(3-iodo-2-(tetrahydro-2H-pyran-2-yl)-2,4,6,7-tetrahydro-5H-pyrazolo[4,3-c]pyridin-5-yl)ethan-1-one (**8**) as a white solid (3.6 g, 76% yield). LC-MS *m/z* [M+H]$^+$ 376.14; $^1$H NMR (400 MHz, DMSO-$d_6$) $\delta$ 5.33 (dd, $J$ = 9.7, 2.6 Hz, 1H), 4.34 – 4.09 (m, 2H), 3.89 – 3.81 (m, 1H), 3.80 – 3.53 (m, 3H), 2.91 – 2.55 (m, 2H), 2.23 – 2.04 (m, 4H), 2.01 – 1.90 (m, 1H), 1.89 – 1.78 (m, 1H), 1.71 – 1.57 (m, 1H), 1.57 – 1.44 (m, 2H).

**Step 6:** Synthesis of 1-(3-(7-(difluoromethyl)-3,4-dihydroquinolin-1(2H)-yl)-1-(tetrahydro-2H-pyran-2-yl)-1,4,6,7-tetrahydro-5H-pyrazolo[4,3-c]pyridin-5-yl)ethan-1-one and 1-(3-(7-(difluoromethyl)-3,4-dihydroquinolin-1(2H)-yl)-2-(tetrahydro-2H-pyran-2-yl)-2,4,6,7-tetrahydro-5H-pyrazolo[4,3-c]pyridin-5-yl)ethan-1-one (**10**)

A mixture of intermediate **8** (2.0 g, 1.0 eq), 7-(difluoromethyl)-1,2,3,4-tetrahydroquinoline (**9**, 1.4 g, 1.2 eq), RuPhos Pd G2 (828 mg, 0.2 eq), RuPhos (500 mg, 0.2 eq), and *t*-BuONa (2.3 g, 4.5 eq) in dioxane (40 mL) was degassed and purged with nitrogen three times. The mixture was then stirred at 110 °C for 12 h. LC-MS analysis indicated complete consumption of **8**, with the formation of a main peak exhibiting the desired mass spectrum. Upon cooling, the mixture was diluted with DCM, filtered through Celite, and the filter cake was washed with DCM. The filtrate was concentrated under reduced pressure, and the resulting residue was purified by flash column chromatography (using a gradient of 0-100% EtOAc/*n*-hexane) to afford 1-(3-(7-(difluoromethyl)-3,4-dihydroquinolin-1(2H)-yl)-1-(tetrahydro-2H-pyran-2-yl)-1,4,6,7-tetrahydro-5H-pyrazolo[4,3-c]pyridin-5-yl)ethan-1-one and 1-(3-(7-(difluoromethyl)-3,4-dihydroquinolin-1(2H)-yl)-2-(tetrahydro-2H-pyran-2-yl)-2,4,6,7-tetrahydro-5H-pyrazolo[4,3-c]pyridin-5-yl)ethan-1-one (**10**, 1.7 g) as a light yellow foam in a yield of 74%. LC-MS *m/z* $[M+H]^+$ 431.28; $^1$H NMR (400 MHz, Chloroform-*d*) *δ* 7.12 – 7.00 (m, 1H), 6.84 – 6.74 (m, 1H), 6.67 (s, 1H), 6.61 – 6.28 (m, 1H), 5.26 – 5.16 (m, 1H), 4.26 – 4.13 (m, 1H), 4.09 – 3.98 (m, 2H), 3.94 – 3.81 (m, 1H), 3.77 – 3.59 (m, 4H), 2.94 – 2.75 (m, 4H), 2.41 – 2.28 (m, 1H), 2.17 – 1.92 (m, 7H), 1.70 – 1.56 (m, 3H); $^{13}$C NMR (101 MHz, $CDCl_3$) *δ* 169.79, 169.13, 149.54, 149.01, 143.00, 138.94, 137.29, 133.16, 132.98, 132.76, 132.73, 132.55, 129.52, 129.36, 126.64, 126.63, 126.42, 117.52, 117.39, 115.38, 115.32, 115.26, 115.15, 115.11, 115.02, 114.99, 112.78, 112.66, 111.20, 111.14, 111.08, 110.99, 110.93, 110.87, 108.01, 107.18, 85.34, 85.30, 67.88, 67.61, 49.49, 49.35, 43.28, 43.00, 38.87, 38.61, 29.44, 29.40, 27.59, 24.98, 24.94, 23.10, 22.69, 22.55, 22.47, 22.37, 22.01, 21.80, 21.53.

**Step 7:** Synthesis of 1-(3-(6-bromo-7-(difluoromethyl)-3,4-dihydroquinolin-1(2H)-yl)-1-(tetrahydro-2H-pyran-2-yl)-1,4,6,7-tetrahydro-5H-pyrazolo[4,3-c]pyridin-5-yl)ethan-1-one and

1-(3-(6-bromo-7-(difluoromethyl)-3,4-dihydroquinolin-1(2H)-yl)-2-(tetrahydro-2H-pyran-2-yl)-2,4,6,7-tetrahydro-5H-pyrazolo[4,3-c]pyridin-5-yl)ethan-1-one (**11**)

A solution of compound **10** (1.7 g, 1.0 eq) in DCM (30 mL) was treated with NBS (667 mg, 0.95 eq) in portions under ice bath conditions. After 2 h, the reaction mixture was diluted with DCM, washed sequentially with aqueous $Na_2S_2O_3$ solution followed by brine, dried over anhydrous $Na_2SO_4$, and concentrated under reduced pressure to remove the solvent. The crude product obtained was purified by flash column chromatography (using a gradient of 0-5% MeOH/DCM) to yield 1-(3-(6-bromo-7-(difluoromethyl)-3,4-dihydroquinolin-1(2H)-yl)-1-(tetrahydro-2H-pyran-2-yl)-1,4,6,7-tetrahydro-5H-pyrazolo[4,3-c]pyridin-5-yl)ethan-1-one and 1-(3-(6-bromo-7-(difluoromethyl)-3,4-dihydroquinolin-1(2H)-yl)-2-(tetrahydro-2H-pyran-2-yl)-2,4,6,7-tetrahydro-5H-pyrazolo[4,3-c]pyridin-5-yl)ethan-1-one (**11**, 1.75 g) as a white foam in a yield of 87%. LC-MS *m/z* $[M+H]^+$ 509.16; $^1H$ NMR (400 MHz, Chloroform-*d*) $\delta$ 7.20 (d, $J$ = 15.9 Hz, 1H), 6.94 – 6.57 (m, 2H), 5.27 – 5.17 (m, 1H), 4.27 – 4.13 (m, 1H), 4.10 – 3.98 (m, 2H), 3.95 – 3.80 (m, 1H), 3.78 – 3.59 (m, 4H), 2.95 – 2.77 (m, 4H), 2.40 – 2.28 (m, 1H), 2.13 – 1.90 (m, 6H), 1.72 – 1.55 (m, 5H).

**Steps 8-9:** Synthesis of 1-(3-(7-(difluoromethyl)-6-(1-methyl-1H-pyrazol-4-yl)-3,4-dihydroquinolin-1(2H)-yl)-1,4,6,7-tetrahydro-5H-pyrazolo[4,3-c]pyridin-5-yl)ethan-1-one (**13**)

In a mixture comprising intermediate **11** (1.28 g, 1.0 eq), 1-methyl-4-(4,4,5,5-tetramethyl-1,3,2-dioxaborolan-2-yl)-1H-pyrazole (1.04 g, 2.0 eq), $PdCl_2$(dppf) (367 mg, 0.2 eq), and $K_2CO_3$ (1.38 g, 4.0 eq) in 1,4-dioxane/$H_2O$ (30 mL/5 mL), the components were degassed and purged with nitrogen three times. Subsequently, the mixture was heated at 100 °C for 12 h. LC-MS analysis confirmed the complete consumption of intermediate **11**, with the formation of a primary peak displaying the desired mass spectrum. Upon cooling, the reaction mixture was quenched with water and subjected to extraction with EtOAc. The collected organic layers were washed with brine, dried over anhydrous $Na_2SO_4$, and concentrated under reduced pressure. The resulting residue underwent purification through flash column chromatography (using a gradient of 0-5% methanol/dichloromethane) to yield a crude product comprising 1-(3-(7-(difluoromethyl)-6-(1-methyl-1H-pyrazol-4-yl)-3,4-dihydroquinolin-1(2H)-yl)-1-(tetrahydro-2H-pyran-2-yl)-1,4,6,7-tetrahydro-5H-pyrazolo[4,3-c]pyridin-5-yl)ethan-1-one and 1-(3-(7-(difluoromethyl)-6-(1-methyl-1H-pyrazol-4-yl)-3,4-dihydroquinolin-1(2H)-yl)-1-(tetrahydro-2H-pyran-2-yl)-1,4,6,7-tetrahydro-5H-pyrazolo[4,3-c]pyridin-5-yl)ethan-1-one (**12**) as a brownish oil.

The crude intermediate **12** was dissolved in a solution of 3 M HCl in MeOH and stirred at room temperature for 12 h. Subsequently, the reaction mixture was concentrated under reduced pressure, dissolved in DCM, and neutralized with aqueous NaOH to achieve a pH of 7~8. The resulting mixture was extracted with DCM, washed with brine, dried over anhydrous $Na_2SO_4$, and concentrated under reduced pressure to yield a crude product. This crude product underwent further purification by flash column chromatography (using a gradient of 0-5% MeOH/DCM) to afford 1-(3-(7-(difluoromethyl)-6-(1-methyl-1H-pyrazol-4-yl)-3,4-dihydroquinolin-1(2H)-yl)-1,4,6,7-tetrahydro-5H-pyrazolo[4,3-c]pyridin-5-yl)ethan-1-one (**13**) as a light yellow solid (640 mg, 60% yield from **11**). LC-MS *m/z* $[M+H]^+$ 427.43; $^1H$ NMR (400 MHz, Methanol-$d_4$) $\delta$ 7.67 – 7.61 (m, 1H), 7.51 (s, 1H), 7.15 – 7.06 (m, 1H), 6.76 – 6.40 (m, 2H), 4.30 – 4.17 (m, 2H), 3.96 – 3.90 (m, 3H), 3.89 – 3.76 (m, 2H), 3.72 – 3.62 (m, 2H), 2.94 – 2.74 (m, 4H), 2.21 – 2.00 (m, 5H); $^{13}C$ NMR (101 MHz, MeOD) $\delta$ 172.48, 172.38, 148.28, 148.17, 143.92, 143.62, 141.83, 141.47, 139.27, 132.39, 132.27, 131.52, 131.32, 131.11, 130.90, 130.00, 129.75, 124.70, 124.37, 120.54, 118.31, 117.26, 117.20, 115.49, 114.86, 113.67, 112.96, 112.57, 112.52, 106.09, 50.89, 50.79, 44.19, 43.76, 39.84, 39.54, 38.97, 27.95, 27.91, 23.54, 23.45, 22.67, 21.64, 21.35.

**Step 10:** Synthesis of *trans*-4-(5-acetyl-3-(7-(difluoromethyl)-6-(1-methyl-1H-pyrazol-4-yl)-3,4-dihydroquinolin-1(2H)-yl)-4,5,6,7-tetrahydro-1H-pyrazolo[4,3-c]pyridin-1-yl)cyclohexane-1-carboxylate (**14**)

To a solution of compound **13** (300 mg, 1.0 eq) and compound **2** (658 mg, 3.0 eq) in DMF (5 mL), $Cs_2CO_3$ (917 mg, 4.0 eq) was added. The resulting mixture was stirred at 70 °C for 7 h. Following the reaction, the mixture was directly subjected to purification by pre-HPLC (45-100% MeCN (0.1% TFA)/$H_2O$ (0.1% TFA) in 55 min). The desired product began eluting when the MeCN/$H_2O$ ratio reached 52%. The compound *trans*-4-(5-acetyl-3-(7-(difluoromethyl)-6-(1-methyl-1H-pyrazol-4-yl)-3,4-dihydroquinolin-1(2H)-yl)-4,5,6,7-tetrahydro-1H-pyrazolo[4,3-c]pyridin-1-yl)cyclohexane-1-carboxylate (**14**, 257 mg) was isolated as a white solid, yielding 64%. LC-MS: *m/z* $[M+H]^+$ = 567.12; $^1H$ NMR (400 MHz, $CDCl_3$-*d*) *δ* 7.64 (s, 1H), 7.46 (s, 1H), 7.08 – 6.97 (m, 1H), 6.83 (s, 1H), 6.47 (td, J = 55.5, 11.6 Hz, 1H), 4.25 (s, 1H), 4.13 (s, 1H), 4.05 – 3.99 (m, 3H), 3.97 – 3.89 (m, 2H), 3.82 – 3.75 (m, 1H), 3.74 – 3.66 (m, 5H), 2.92 – 2.81 (m, 3H), 2.80 – 2.73 (m, 1H), 2.47 – 2.35 (m, 1H), 2.24 – 2.01 (m, 11H), 1.69 – 1.52 (m, 2H); $^{13}C$ NMR (101 MHz, $CDCl_3$) *δ* 175.75, 175.71, 171.80, 171.22, 148.60, 148.01, 142.27, 142.20, 137.79, 137.46, 137.30, 136.57, 131.35, 130.72, 130.50, 130.00, 129.94, 129.79, 129.73, 129.58, 129.52, 126.68, 126.60, 120.19, 120.08, 119.87, 115.93, 115.84, 113.57, 113.49, 111.18, 110.65, 105.94, 105.39, 57.65, 57.56, 51.78, 49.91, 49.78, 43.47, 43.17, 41.79, 41.73, 39.42, 39.31, 38.56, 38.49, 31.32, 28.10, 28.07, 27.38, 27.34, 22.71, 22.17, 22.08, 21.82, 21.09, 20.75.

**Step 11:** Synthesis of *trans*-4-(5-acetyl-3-(7-(difluoromethyl)-6-(1-methyl-1H-pyrazol-4-yl)-3,4-dihydroquinolin-1(2H)-yl)-4,5,6,7-tetrahydro-1H-pyrazolo[4,3-c] 9 yridine-1-yl)cyclohexane-1-carbaldehyde (**15**)

Compound **14** (232 mg, 1.0 eq) was dissolved in anhydrous DCM (15 mL), and the solution was degassed by purging with nitrogen three times. DIBAL (25% in toluene, 1.1 mL, 4.0 eq) was then added dropwise at -78 °C over 1 h. The reaction mixture was stirred at -78 °C for an additional 2 h. Subsequently, the reaction was quenched with aqueous potassium sodium tartrate and warmed to room temperature. The resulting mixture was extracted with DCM, and the organic layers were washed with brine, dried over anhydrous $Na_2SO_4$, filtered, and concentrated under reduced pressure. The crude residue was purified by pre-HPLC (35-100% MeCN/$H_2O$ in 65 min). The desired product began eluting when the MeCN/$H_2O$ ratio reached 44%. The compound *trans*-4-(5-acetyl-3-(7-(difluoromethyl)-6-(1-methyl-1H-pyrazol-4-yl)-3,4-dihydroquinolin-1(2H)-yl)-4,5,6,7-tetrahydro-1H-pyrazolo[4,3-c]pyridin-1-yl)cyclohexane-1-carbaldehyde (**15**) was isolated as a white solid (85 mg, 39% yield). LC-MS: *m/z* $[M+H]^+$ 537.10; $^1$H NMR (400 MHz, $CDCl_3$-*d*) $\delta$ 9.69 (d, $J$ = 3.5 Hz, 1H), 7.56 – 7.51 (m, 1H), 7.43 – 7.37 (m, 1H), 7.07 – 6.96 (m, 1H), 6.89 – 6.83 (m, 1H), 6.51 (td, J = 55.6, 10.9 Hz, 1H), 4.25 (s, 1H), 4.12 (s, 1H), 3.98 – 3.93 (m, 3H), 3.93 – 3.82 (m, 2H), 3.78 – 3.64 (m, 3H), 2.91 – 2.82 (m, 2H), 2.81 – 2.69 (m, 2H), 2.40 – 2.28 (m, 1H), 2.24 – 2.00 (m, 11H), 1.51 – 1.36 (m, 2H); $^{13}$C NMR (101 MHz, $CDCl_3$) $\delta$ 203.48, 171.88, 171.38, 148.46, 147.96, 142.12, 142.00, 138.07, 137.11, 137.07, 136.93, 131.39, 130.97, 130.73, 129.97, 129.76, 129.52, 126.87, 126.75, 120.09, 119.89, 115.88, 115.80, 113.53, 113.45, 111.39, 111.10, 110.77, 108.09, 105.82, 105.33, 57.74, 57.62, 53.97, 49.89, 49.78, 48.58, 48.52, 43.37, 43.07, 39.35, 39.21, 38.43, 38.38, 30.94, 27.25, 27.20, 25.01, 22.59, 22.08, 22.01, 21.73, 20.96, 20.60.

**Step 12:** Synthesis of 5-(4-(((1r,4r)-4-(5-acetyl-3-(7-(difluoromethyl)-6-(1-methyl-1H-pyrazol-4-yl)-3,4-dihydroquinolin-1(2H)-yl)-4,5,6,7-tetrahydro-1H-pyrazolo[4,3-c]pyridin-1-

yl)cyclohexyl)methyl)piperazin-1-yl)-2-(2,6-dioxopiperidin-3-yl)isoindoline-1,3-dione (**16**, **CBPD-409**)

$NaBH(OAc)_3$ (117 mg, 3.0 eq) was added in portions over 2 h to a mixture of **15** (100 mg, 1.0 eq) and **4** (110 mg, 1.3 eq) suspended in DCE/DMF (8 mL/4 mL). The reaction was then stirred at room temperature for 12 h. After completion, the mixture was concentrated under reduced pressure. The resulting residue was subjected to purification by pre-HPLC (30-100% MeCN (0.1% TFA) /$H_2O$ (0.1% TFA) in 70 min). The final compound, 5-(4-(((1r,4r)-4-(5-acetyl-3-(7-(difluoromethyl)-6-(1-methyl-1H-pyrazol-4-yl)-3,4-dihydroquinolin-1(2H)-yl)-4,5,6,7-tetrahydro-1H-pyrazolo[4,3-c]pyridin-1-yl)cyclohexyl)methyl)piperazin-1-yl)-2-(2,6-dioxopiperidin-3-yl)isoindoline-1,3-dione (**16**, **CBPD-409**), was obtained as a light yellow solid (78 mg, 48% yield). UPLC-MS: 1.69 min, purity > 95%; MS (ESI) *m/z* calcd. For $C_{46}H_{52}F_2N_{10}O_5$ $[M + H]^+$ 863.42, found 863.43; HRMS (APCI) *m/z* calcd. For $C_{46}H_{52}F_2N_{10}O_5$ $[M + H]^+$ 863.4163, found 863.4189; $^1H$ NMR (400 MHz, MeOH-$d_4$) $\delta$ 7.77 (d, $J$ = 8.5 Hz, 1H), 7.64 (s, 1H), 7.54 – 7.46 (m, 2H), 7.36 (dd, $J$ = 8.5, 2.3 Hz, 1H), 7.11 (d, $J$ = 8.9 Hz, 1H), 6.77 – 6.40 (m, 2H), 5.10 (dd, $J$ = 12.5, 5.4 Hz, 1H), 4.31 – 4.03 (m, 5H), 3.97 – 3.78 (m, 6H), 3.67 (q, $J$ = 6.7 Hz, 4H), 3.51 – 3.34 (m, 3H), 3.18 (d, $J$ = 6.3 Hz, 2H), 2.97 – 2.79 (m, 5H), 2.79 – 2.65 (m, 2H), 2.19 (s, 2H), 2.16 – 1.97 (m, 11H), 1.45 – 1.27 (m, 2H); $^{13}C$ NMR (101 MHz, MeOD) $\delta$ 174.57, 172.40, 172.26, 171.62, 168.96, 168.67, 155.71, 150.22, 149.77, 143.82, 139.82, 139.33, 139.07, 135.49, 132.28, 132.16, 131.25, 131.23, 131.00, 130.79, 127.51, 127.37, 126.12, 122.76, 122.22, 121.22, 120.89, 120.79, 120.47, 118.36, 117.39, 115.50, 115.05, 112.71, 112.64, 111.47, 111.03, 110.97, 110.41, 107.57, 63.43, 58.16, 52.98, 51.11, 51.03, 50.54, 45.86, 44.60, 43.89, 40.16, 39.66, 38.93, 32.95, 32.47, 32.19, 30.50, 28.44, 23.71, 23.38, 23.25, 22.59, 21.67, 21.38.

#### 3. Method and procedure for the preparation of CBPD-409-Me

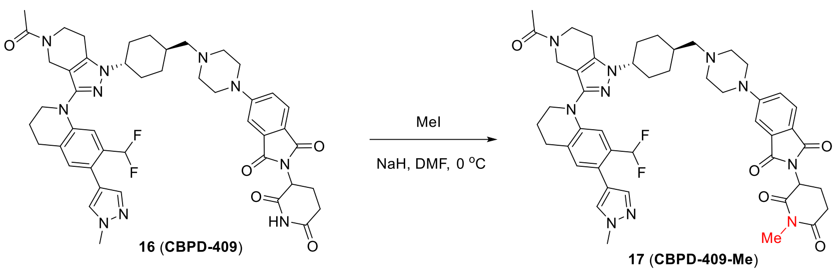


To a solution of **16** (10.8 mg, 1.0 eq) in DMF (1 mL) was added NaH (4.4 mg, 10.0 eq) under ice bath condition, followed by MeI (3.44 µL, 5.0 eq). The reaction was stirred for 30 min, then quenched with TFA and water. The resulting mixture was directly purified by pre-HPLC (30-100% MeCN/$H_2O$ in 70 min) to give product 5-(4-(((1r,4r)-4-(5-acetyl-3-(7-(difluoromethyl)-6-(1-methyl-1H-pyrazol-4-yl)-3,4-dihydroquinolin-1(2H)-yl)-4,5,6,7-tetrahydro-1H-pyrazolo[4,3-c]pyridin-1-yl)cyclohexyl)methyl)piperazin-1-yl)-2-(1-methyl-2,6-dioxopiperidin-3-yl)isoindoline-1,3-dione (**17**, **CBPD-409-Me**) as a light yellow solid (9.0 mg, 82% yield): UPLC-MS: 1.47 min, purity > 95%; MS (ESI) *m/z* calcd. For $C_{47}H_{54}F_2N_{10}O_5$ $[M + H]^+$ 877.43, found 866.80; $^1H$ NMR (400 MHz, Methanol-$d_4$) $\delta$ 7.77 (d, $J$ = 8.5 Hz, 1H), 7.64 (s, 1H), 7.54 – 7.47 (m, 2H), 7.37 (dd, $J$ = 8.5, 2.3 Hz, 1H), 7.15 – 7.07 (m, 1H), 6.79 – 6.40 (m, 2H), 5.12 (dd, $J$ = 12.9, 5.4 Hz, 1H), 4.32 – 4.02 (m, 5H), 3.97 – 3.79 (m, 6H), 3.78 – 3.59 (m, 4H), 3.56 – 3.32 (m, 3H), 3.22 – 3.11 (m, 5H), 2.98 – 2.79 (m, 6H), 2.76 – 2.63 (m, 1H), 2.19 (s, 2H), 2.14 – 1.98 (m, 11H), 1.43 – 1.32 (m, 2H).

#### 4. UPLC-MS spectra

##### The UPLC-MS spectrum for compound 16 (CBPD-409)

**SAMPLE INFORMATION**

| | | | |
|---|---|---|---|
| Sample Name: | ZXC-409-Weeks | Acquired By: | System |
| Sample Type: | Unknown | Sample Set Name: | 1 |
| Vial: | 1:D,8 | Acq. Method Set: | 10to100Bin5min_noDelay |
| Injection #: | 1 | Processing Method: | Default |
| Injection Volume: | 6.00 ul | Channel Name: | 254.0nm |
| Run Time: | 5.0 Minutes | Proc. Chnl. Descr.: | PDA Spectrum PDA 254.0 nm |
| Date Acquired: | 7/29/2022 10:08:59 PM EDT | | |
| Date Processed: | 7/29/2022 10:23:11 PM EDT | | |

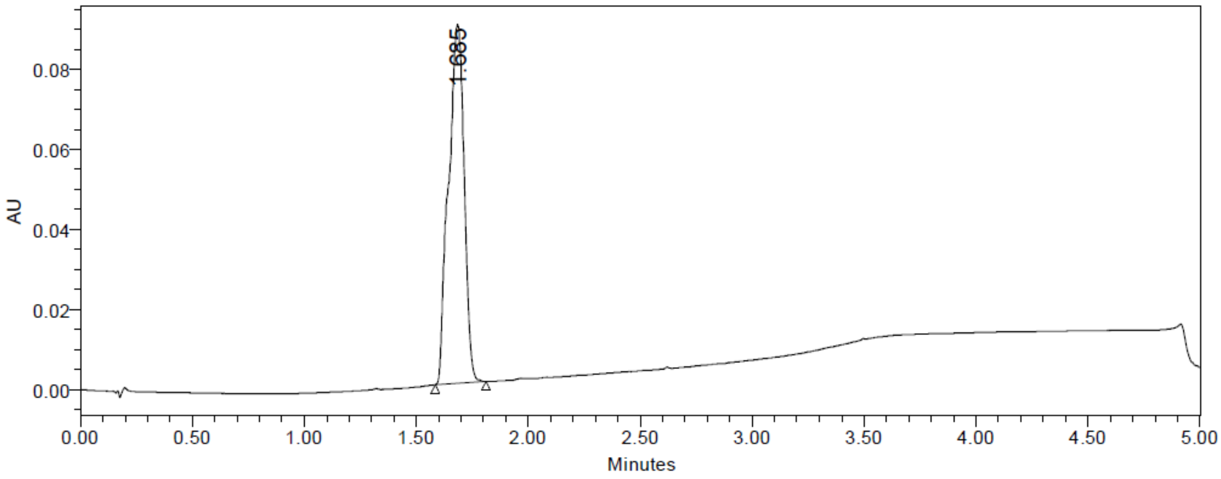


| | RT | Area | % Area | Height |
|---|---|---|---|---|
| 1 | 1.685 | 433496 | 100.00 | 89466 |

**The UPLC-MS spectrum for compound 17 (CBPD-409-Me)**

SAMPLE INFORMATION

| | | | |
|---|---|---|---|
| Sample Name: | ZXC-7-261-P | Acquired By: | System |
| Sample Type: | Unknown | Date Acquired: | 5/7/2023 1:28:57 AM EDT |
| Vial: | 1:A,4 | Acq. Method Set: | New10to 100% B 5 min_NoDelay |
| Injection #: | 1 | Date Processed: | 5/7/2023 3:14:43 AM EDT |
| Injection Volume: | 6.00 ul | Processing Method: | Bruce1 |
| Run Time: | 5.0 Minutes | Channel Name: | 254.0nm |
| Sample Set Name: | 3 | Proc. Chnl. Descr.: | PDA Spectrum PDA 254.0 nm (PDA |

Auto-Scaled Chromatogram

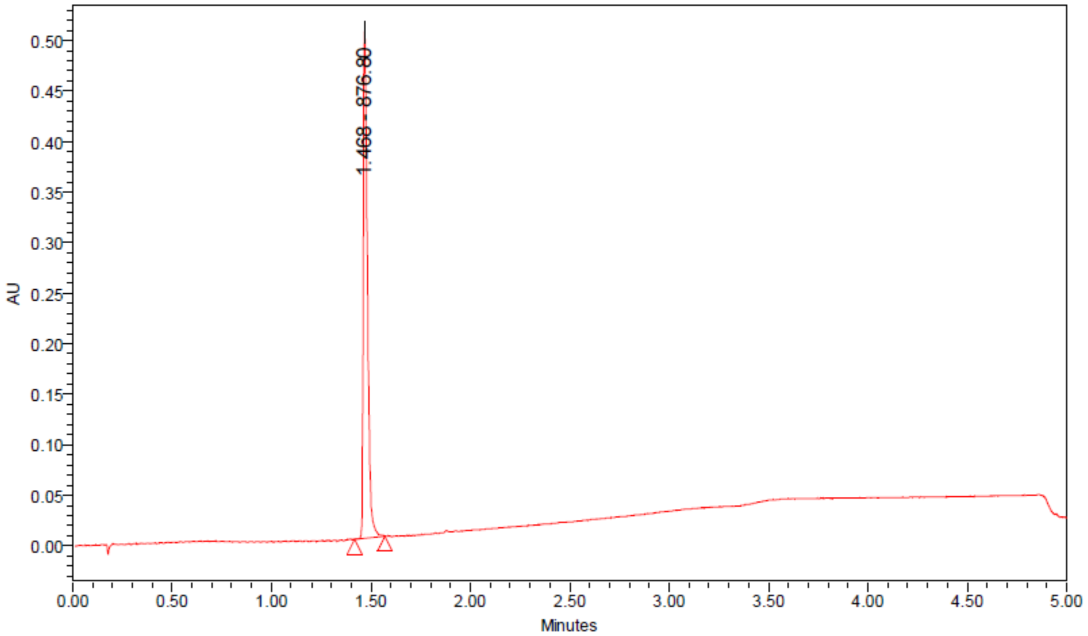


Peak Results

| | RT | Area | Height | % Area |
|---|---|---|---|---|
| 1 | 1.468 | 741778 | 501466 | 100.00 |

Peak Results

| | Base Peak (m/z) |
|---|---|
| 1 | 876.80 |

#### 5. NMR spectra

##### [1]H-NMR spectrum for compound 2

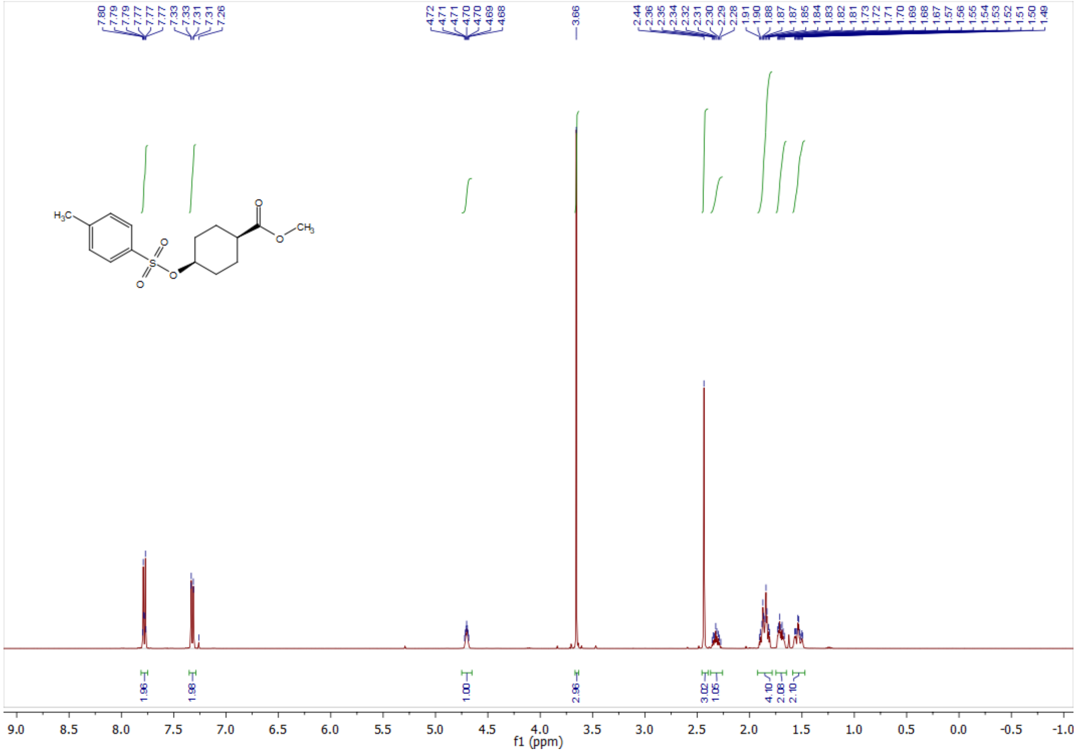

##### [1]H-NMR spectrum for compound 4

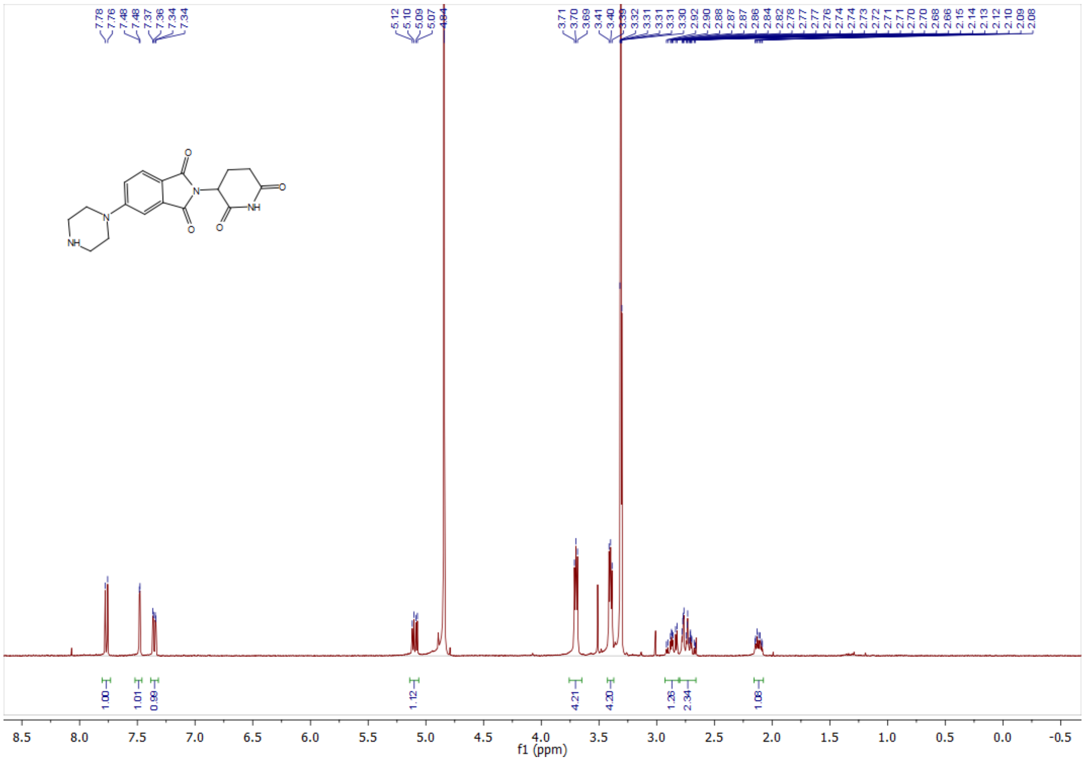

### $^1$H-NMR spectrum for compound 7

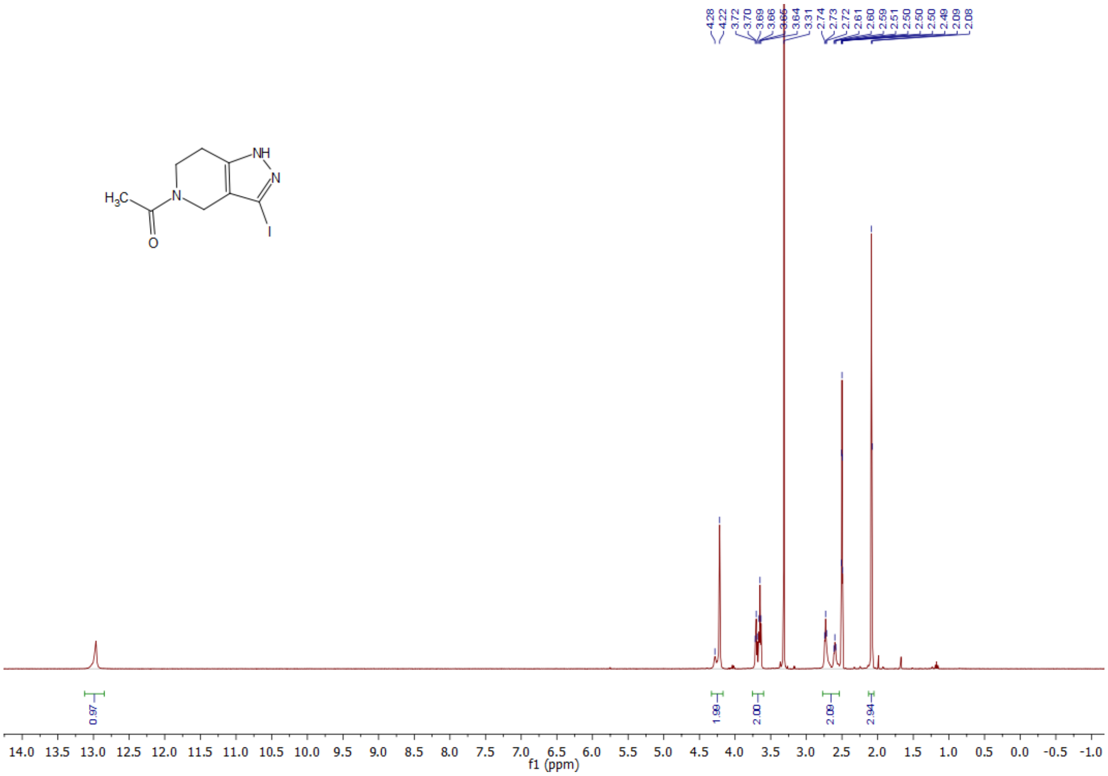


### $^1$H-NMR spectrum for compound 8

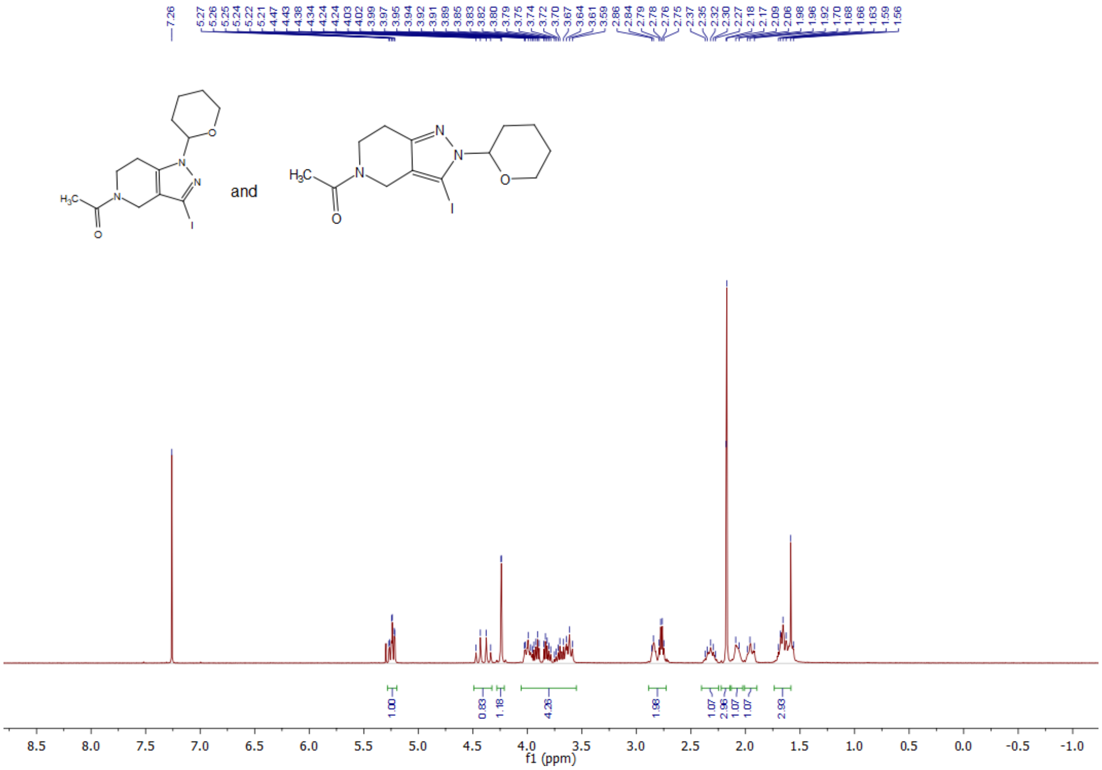

### [1]H-NMR spectrum for compound 10

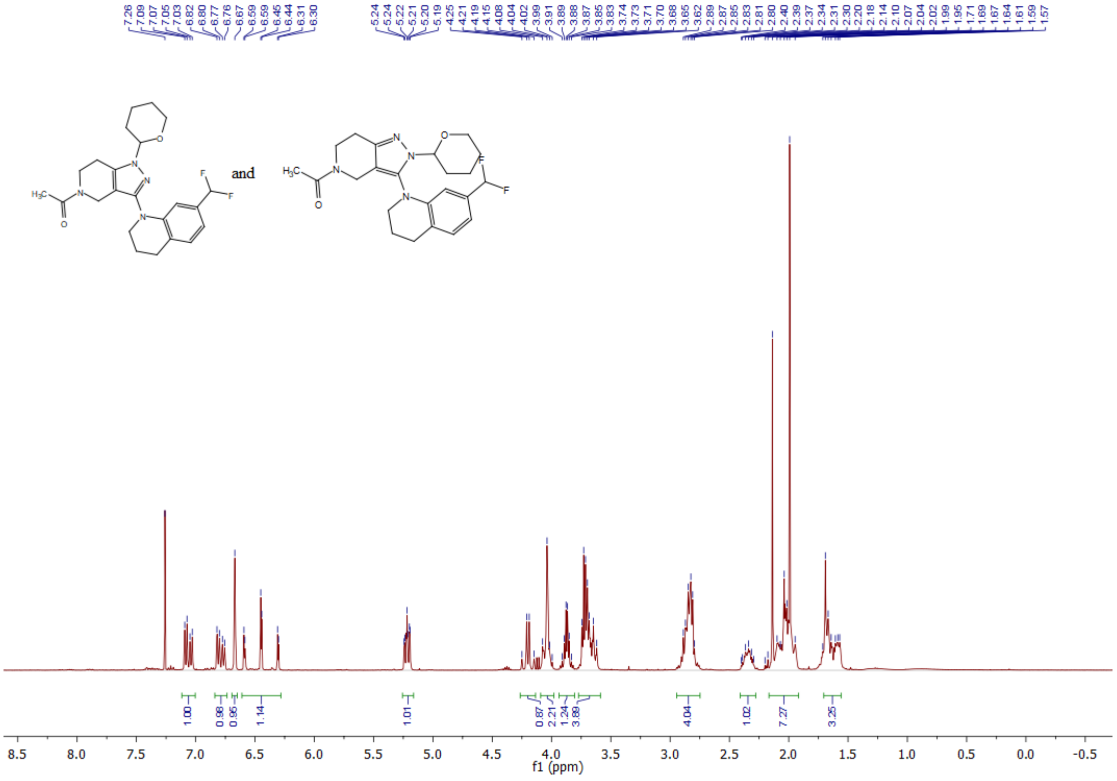


### [13]C-NMR spectrum for compound 10

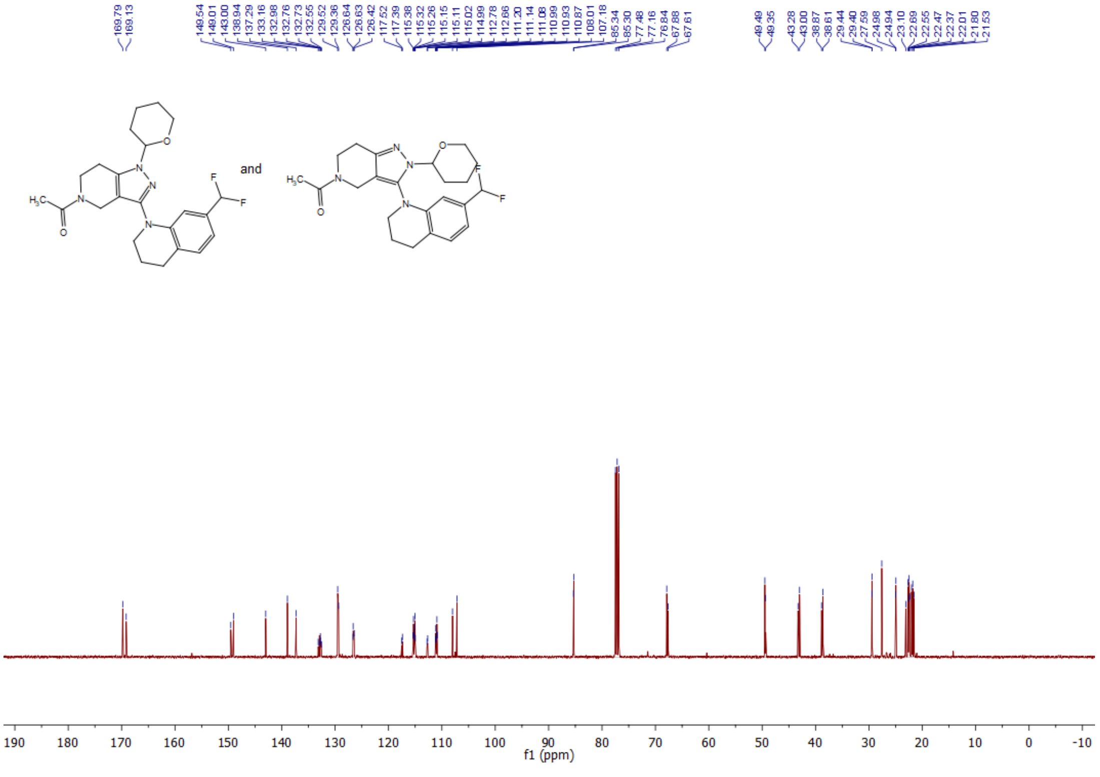

#### [1]H-NMR spectrum for compound 13

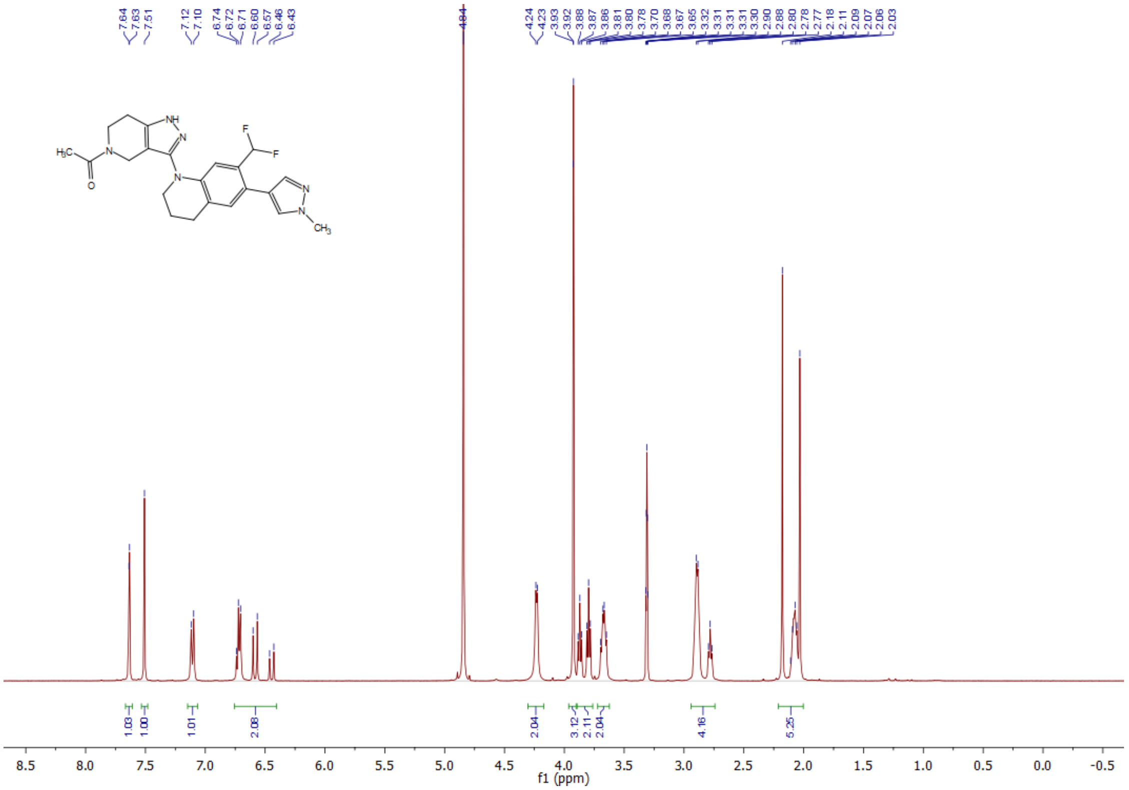

#### [13]C-NMR spectrum for compound 13

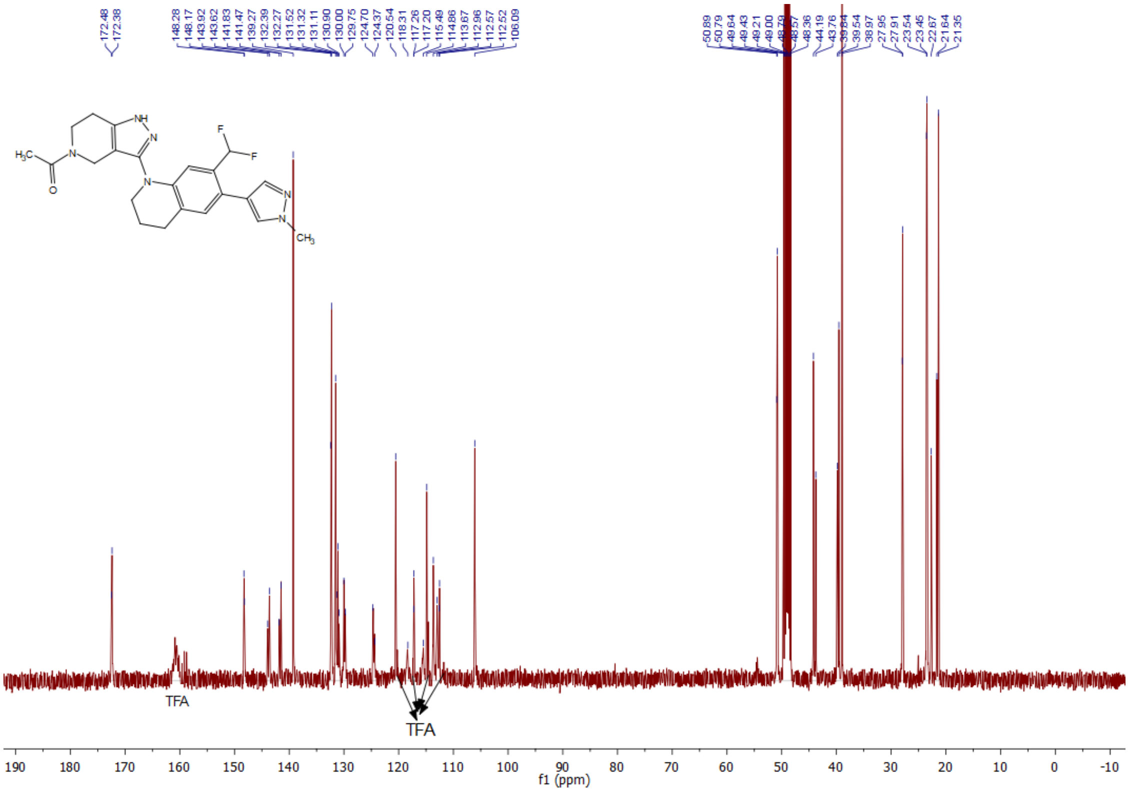

#### [1]H-NMR spectrum for compound 14

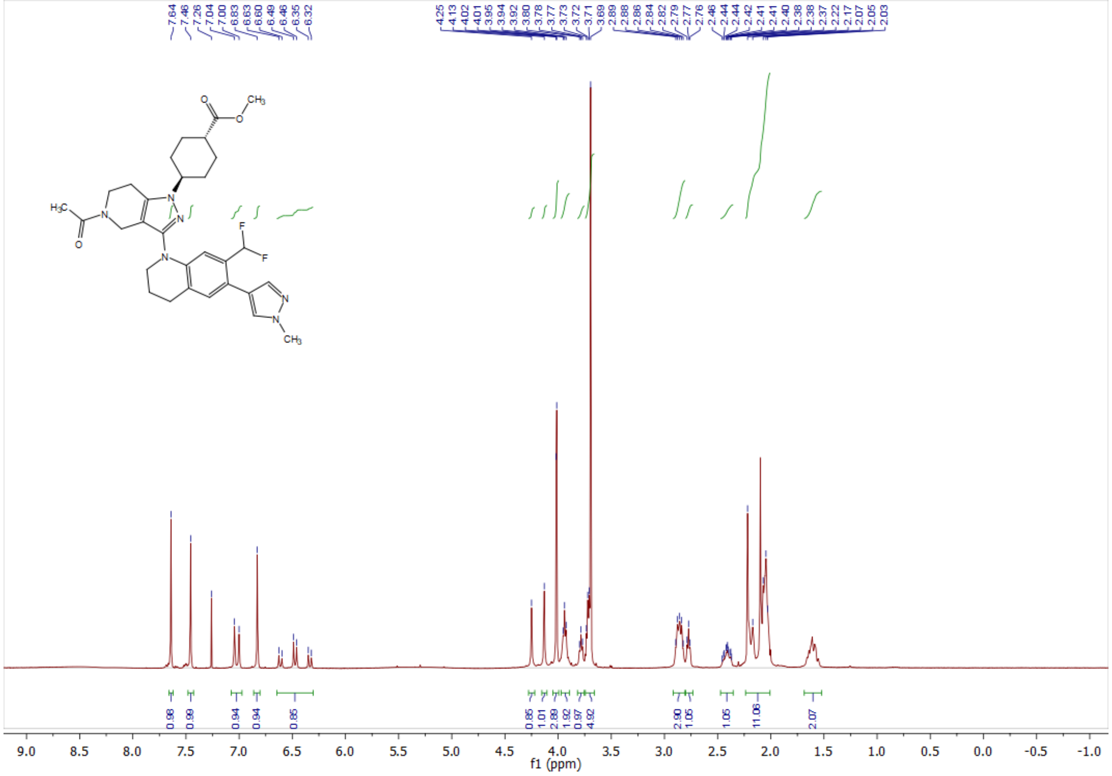

#### [13]C-NMR spectrum for compound 14

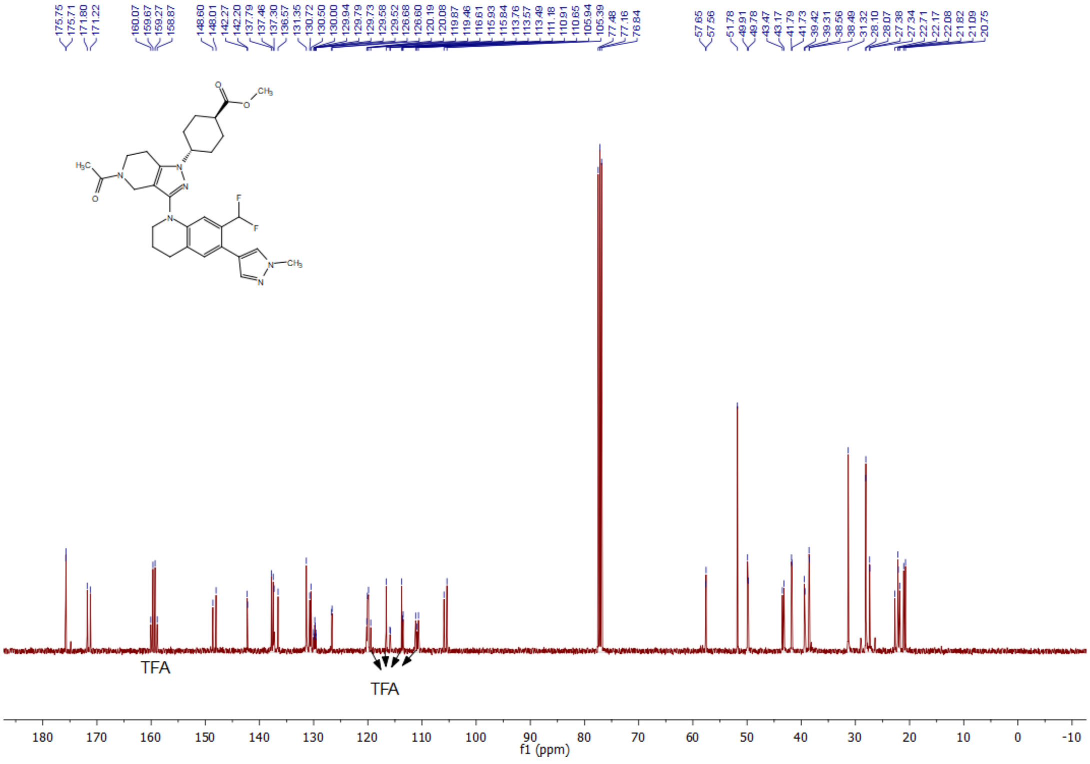

#### [1]H-NMR spectrum for compound 15

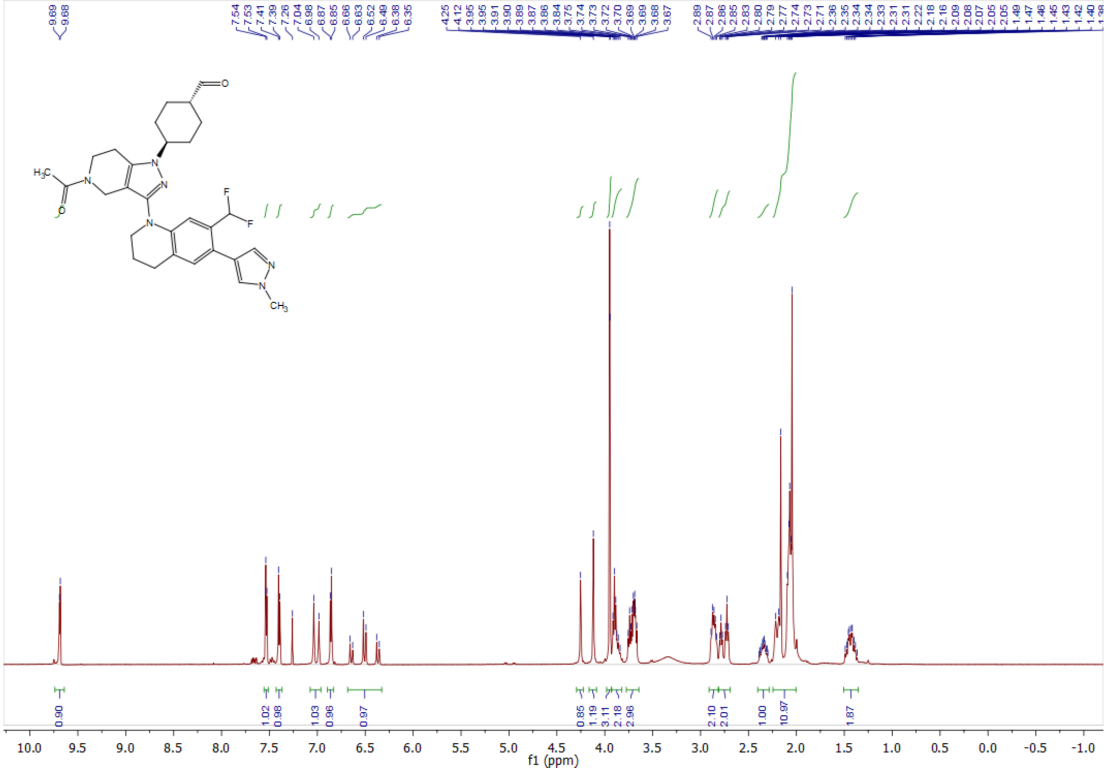

#### [1]H-NMR spectrum for compound 16 (CBPD-409)

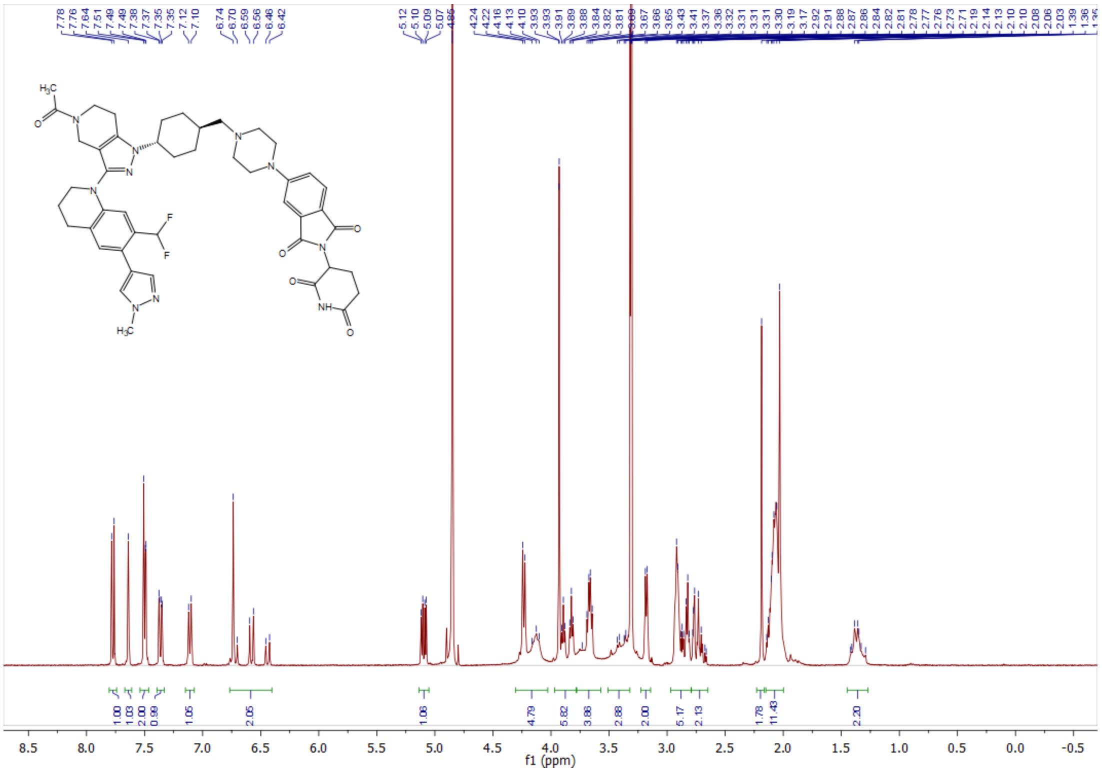

#### $^{13}$C-NMR spectrum for compound 16 (CBPD-409)

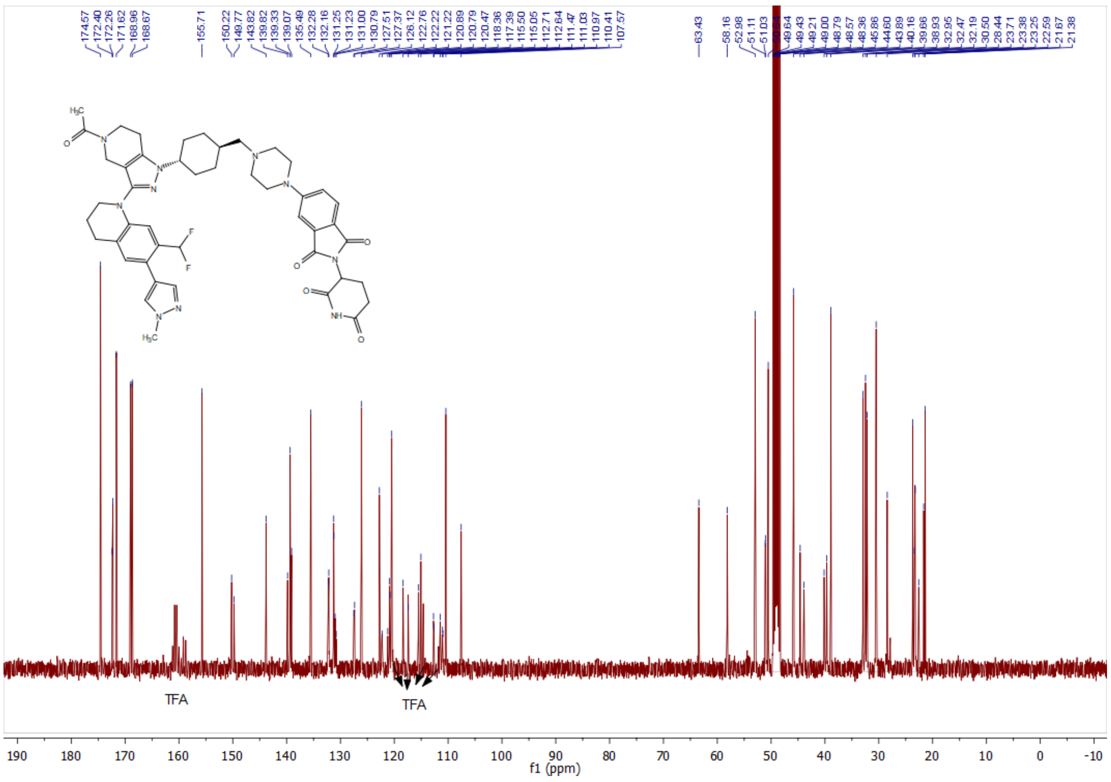


#### $^{1}$H-NMR spectrum for compound 17 (CBPD-409-Me)

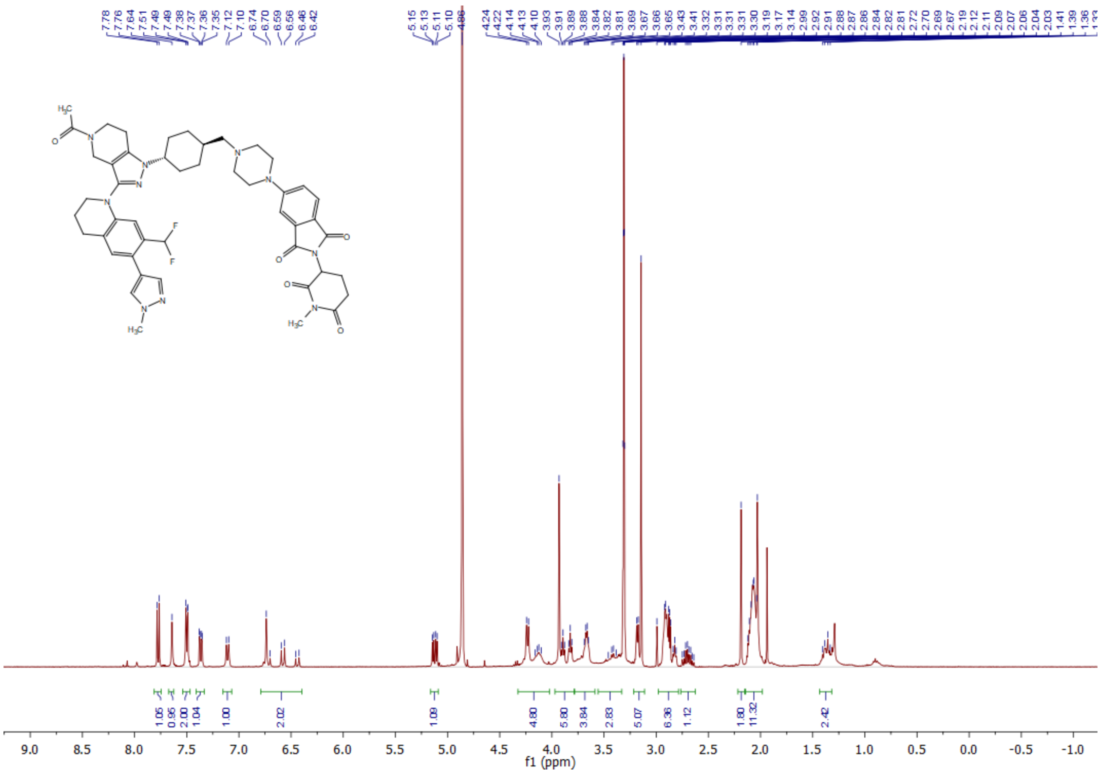
